# The connectome of an insect brain

**DOI:** 10.1101/2022.11.28.516756

**Authors:** Michael Winding, Benjamin D. Pedigo, Christopher L. Barnes, Heather G. Patsolic, Youngser Park, Tom Kazimiers, Akira Fushiki, Ingrid V. Andrade, Avinash Khandelwal, Javier Valdes-Aleman, Feng Li, Nadine Randel, Elizabeth Barsotti, Ana Correia, Richard D. Fetter, Volker Hartenstein, Carey E. Priebe, Joshua T. Vogelstein, Albert Cardona, Marta Zlatic

**Affiliations:** University of Cambridge, Department of Zoology; Cambridge, UK; Johns Hopkins University, Department of Biomedical Engineering; Baltimore, MD, USA; University of Cambridge, Department of Physiology, Development, and Neuroscience; Cambridge, UK; Johns Hopkins University, Department of Applied Mathematics and Statistics; Baltimore, MD, USA; Accenture; Arlington, VA, USA; Johns Hopkins University, Center for Imaging Science; Baltimore, MD, USA; Janelia Research Campus, Howard Hughes Medical Institute; Ashburn, VA, USA; kazmos GmbH; Dresden, Germany; Zuckerman Mind Brain Behavior Institute, Columbia University; New York, NY, USA; University of California Los Angeles, Department of Molecular, Cell and Developmental Biology; Los Angeles, CA, USA; MRC Laboratory of Molecular Biology, Neurobiology Division; Cambridge, UK

## Abstract

Brains contain networks of interconnected neurons, so knowing the network architecture is essential for understanding brain function. We therefore mapped the synaptic-resolution connectome of an insect brain (*Drosophila* larva) with rich behavior, including learning, value-computation, and action-selection, comprising 3,013 neurons and 544,000 synapses. We characterized neuron-types, hubs, feedforward and feedback pathways, and cross-hemisphere and brain-nerve cord interactions. We found pervasive multisensory and interhemispheric integration, highly recurrent architecture, abundant feedback from descending neurons, and multiple novel circuit motifs. The brain’s most recurrent circuits comprised the input and output neurons of the learning center. Some structural features, including multilayer shortcuts and nested recurrent loops, resembled powerful machine learning architectures. The identified brain architecture provides a basis for future experimental and theoretical studies of neural circuits.

**One-Sentence Summary:** We generated a synaptic-resolution brain connectome and characterized its connection types, neuron types, and circuit motifs.

## Introduction

A defining characteristic of a brain is its synaptic wiring diagram, or connectome: which neurons synapse onto each other to form circuits. A synapse-resolution connectome is therefore an essential prerequisite for understanding brain function that can inspire understanding and guide analysis by constraining modeling and experimental studies (*1–3*). To date, complete synaptic-resolution connectomes from volume electron microscopy (EM) have only been mapped for three organisms with up to several hundred brain neurons, namely the nematode *C. elegans* (*4, 5*), the larva of the sea squirt *Ciona intestinalis* (*6*), and the larva of the marine annelid *Platynereis dumerilii* (*7*). Due to technological constraints, reconstructing and proofreading circuits from larger brains has been extremely challenging. Synapse-resolution circuitry of larger brains has therefore been approached with a reductionist framework that considers select subregions in isolation (*8–14*). Yet, neural activity across whole brains of multiple species reveals widespread responses to limited stimulation (*15–17*), suggesting pervasive, strong interconnectivity between brain regions, as also reported in classical studies of the primate cerebral cortex (*18*). Large-scale recording of functional activity in invertebrates (*16, 19–21*) and vertebrates (*15, 17, 22, 23*) demonstrates that neural computations occur across spatially dispersed brain regions, further motivating brain-wide circuit studies. In the light of pervasive connectivity between brain regions, our understanding of any one brain region requires analyzing it in the context of its synaptic inputs and outputs to other regions.

We therefore sought to generate a comprehensive synapse-resolution connectivity map, including both hemispheres and all inputs and outputs, of a relatively complex brain of a small insect with several thousand neurons and a rich behavioral repertoire. We settled on an organism, the larva of the fruit fly *Drosophila melanogaster*, that has a relatively compact brain, so that it can be imaged at nanometer scale with EM and its circuits reconstructed within a reasonable time frame. Its brain structures are homologous to those of adult *Drosophila* and larger insects (*24–26*) and its neuronal connectivity is relatively stable throughout larval life (*27*). The larva has a rich repertoire of adaptive behaviors, including several modes of locomotion and many kinds of sequences of actions (*28–30*), and can form short- and long-term associative memories (*25, 31, 32*), and use these memories to guide value-computation and action-selection (*33, 34*). Furthermore, an exquisite genetic toolkit (*35–37*) enables single-cell opto- and electrophysiological recordings to experimentally test hypotheses generated by the connectome about the roles of specific circuit motifs in distinct behavioral tasks (*28, 29, 31, 34, 38*).

We mapped all neurons in the brain using computer-assisted reconstruction with CATMAID (*39, 40*) in a previously acquired nanometer-resolution EM volume of the central nervous system (CNS) of the *Drosophila* larva (*38*) and we annotated their synapses. Prior studies have used the same EM volume to map sensory and premotor circuits in the ventral nerve cord (VNC) and subesophageal zone (SEZ) (*38, 41–44*), as well specific subregions in the brain (a total of 452 sensory and 1054 brain neurons): the primary sensory neuropils (*45–49*) and the higher order center for learning and memory (the mushroom body, MB) (*25, 31, 34*). Here, we took a dense circuit mapping approach to reconstruct the remaining 1507 brain neurons. We found that the brain received axons from 477 sensory/ascending neurons and comprised 2,536 differentiated neurons (3,013 neurons total), the majority of which were not previously reconstructed in areas that have been dubbed “terra incognita” (*10*).

We performed a detailed analysis of the 3,013-neuron connectome, including the connection-types, neuron-types, hubs, brain-wide circuit motifs, and interhemispheric and brain-nerve cord interactions. We paired all uniquely identifiable, mirror-symmetric left-right homologous neuron pairs from the two brain hemispheres. 37% of neurons displayed contralateral branches that link the two hemispheres, highlighting that mapping both hemispheres is fundamentally required to understand the brain. We performed an unbiased hierarchical clustering of all neurons based solely on their synaptic connectivity and identified 90 neuron clusters. These connectivity-defined neuron types were internally consistent for other features, such as morphology and function, when those features were known. We developed analysis tools to characterize brainwide multihop pathways and used them to uncover general principles of multisensory integration, interhemispheric communication, feedback, and brain-nerve cord interactions. We found extensive feedback from descending neurons back to other brain neurons, suggesting efference copy signals may heavily influence brain processing. Overall, the brain had a highly recurrent architecture with 41% of neurons participating in polysynaptic recurrent loops. We found that the dopaminergic neurons (DANs) that drive learning were amongst the most recurrent in the brain, suggesting recurrence may be especially important for learning. Structural features identified in the brain, including multilayer shortcuts and prominent nested recurrent loops, were reminiscent of powerful machine learning architectures (*50–52*). The connectome presented here will guide analysis and provide a basis for many future theoretical and experimental investigations (*12, 53, 54*) of the structure-function relationships of neural circuits (*29, 38, 55–57*), and may provide inspiration for new machine-learning architectures.

## Results

### Reconstruction of the Drosophila larva brain in a full-CNS electron microscopy volume

We previously generated a synaptic-resolution EM volume of the CNS of the first instar *Drosophila* larva (*38, 40*). This volume contains all CNS neurons, as well as sensory neuron axons and motor neuron dendrites, enabling reconstruction of all neural pathways from sensory input to motor output. Previous studies have used this EM volume to reconstruct most sensory inputs to the brain (452 input neurons), their downstream partners, and the higher-order learning center (total 1,054 brain neurons). Here, we reconstructed the remaining 1,507 brain neurons, resulting in a total of 3,013 neurons and approximately 544,000 synaptic sites (Fig. 1A, B; Fig. S1A, B). The vast majority of neurons (>99%) were reconstructed to completion and the majority of annotated synaptic sites in the brain (75%) were linked with a neuron (Fig. 1B). The remaining 25% were mostly composed of small dendritic fragments, which are labor intensive to reconstruct and provide diminishing returns. Prior studies have shown that neurons make multiple connections with the same partner on different dendritic branches (*40*), so orphaned synapses may affect synaptic weights of known connections but are unlikely to add entirely new strong connections or change conclusions about strongly connected pathways.

**Fig. 1:**
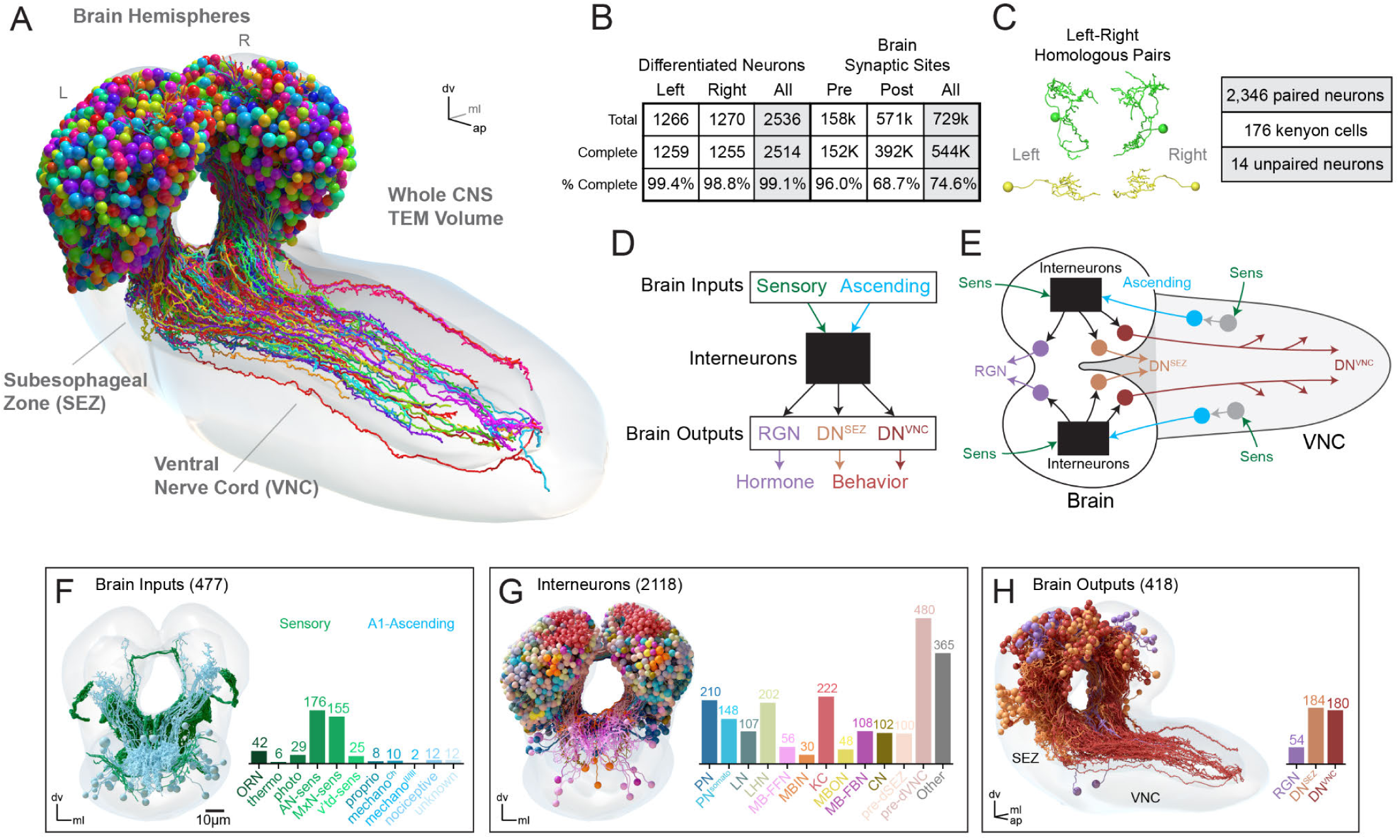
Comprehensive reconstruction of a *Drosophila* larva brain. (**A**) Morphology of differentiated brain neurons in the central nervous system (CNS) of a *Drosophila* larva. All neurons were reconstructed in both hemispheres, including all sensory neurons (SNs) to the brain and axons of brain descending neurons (DNs), including those entering the Subesophageal Zone (SEZ) and the Ventral Nerve Cord (VNC). The CNS is approximately 105 μm x 120 μm x 240 μm (medial-lateral [ml], dorsal-ventral [dv], anterior-posterior [ap] axes, respectively). (**B**) Characterization of brain neurons and their synaptic sites. >99% of neurons were reconstructed to completion, defined by reconstruction of all leaf nodes in the skeleton (see Methods) and no data quality issues preventing identification of axon and dendrite. Synaptic sites in the brain and associated with brain neurons were quantified. Pre- and postsynaptic sites were considered complete when connected to a brain neuron or to the arbor of a neuron outside the brain. (**C**) Left/right homologous neuron pairs were identified using automated graph matching with manual proofreading. 14 neurons displayed no clear partner based on this workflow (unpaired), along with 176 unpaired kenyon cells (KCs) in the learning/memory center. (**D** and **E**) Schematic overview of brain structure, including input neurons, interneurons, and output neurons. Brain inputs include SNs, which directly synapse onto brain neurons, and ascending neurons (ANs) from VNC segment A1, which receive direct or multi-hop input from A1 sensories (see Fig. S2). Brain interneurons transmit these input signals to output neurons: DNs to the SEZ (DN^SEZ^), DNs to the VNC (DN^VNC^), and ring gland neurons (RGN). (**F** to **H**) Cell classes in the brain. Some interneurons belong to multiple classes, but are displayed in the paper as mutually exclusive for plotting expedience (see Fig. S4).

Most neurons in *Drosophila* are mirrored across hemispheres, such that each neuron has a hemilateral homolog in the opposite hemisphere (*40*). We identified all homologous hemilateral partners using automated graph matching (*58–60*) followed by manual review. These pairings were robust across a variety of independent morphological and connectivity metrics (Fig. S1E, F). Our current data suggests that 93% of brain neurons have hemilateral homologous partners in the opposite hemisphere (Fig. 1C), while the Kenyon cells (KC, 176 mature neurons) in the learning/memory center comprise the vast majority of unpaired neurons (*25*).

These homologous partners were used to identify potential reconstruction errors, namely instances of morphological asymmetries between partners, and to target review to such neurons (Fig. S1D). To assess the effectiveness of this targeted review, we randomly selected ten brain interneurons and fully reviewed them according to previously described methods (*38, 40*). We found that most (74%) neuron→neuron connections, or edges, remained unchanged. Edges that did change after review mostly displayed a modest increase in synaptic strength, suggesting errors of omission, the most common type of error as described in previous connectomics studies (*10, 40*) (Fig. S1G, H).

In the following sections, we investigate neuron and connection types, the flow of information from inputs to outputs, multisensory integration, cross-hemisphere interactions, feedback from outputs to inputs, the level of recurrence in the brain and brain-nerve cord interactions.

### Identification of all brain input neurons, interneurons, and output neurons

To facilitate the analysis of the connectome, we defined and identified a curated set of broad neuron classes based on available prior information about some brain neurons. All brain neurons were divided into three general categories: input neurons, output neurons, and interneurons (Fig. 1D, E). Brain input neurons (Fig. 1F) comprise two broad classes: 1) sensory neurons (SNs) with axons in the brain; many were reconstructed and characterized previously (*45–47, 61*), and 2) ascending neurons (ANs) that transmit somatosensory signals from the VNC; several were previously reconstructed (*38, 41, 44, 62–64*). We reconstructed an additional set of A1 ANs, resulting in the most complete set of ANs from a nerve cord segment (A1) to date (Fig. S2). Brain output neurons comprise three broad classes: those with axons that terminate in the ring gland (RGNs), descend to the SEZ (DNs^SEZ^), or descend into the VNC (DNs^VNC^) (Fig. 1H). The full set of RGNs have been previously described (*41, 47, 49, 65, 66*), while DNs^SEZ^ and DNs^VNC^ were reconstructed and identified here based on axon projections (Fig. S3).

Brain interneurons comprised all neurons with cell bodies and axons/dendrites in the brain. We subdivided interneurons into neuron classes based on previously known function or direct connectivity with neurons of known function (Fig. 1G, Fig. S4). We started with sensory input neurons and identified all of their projection neurons (PNs) in the primary sensory neuropils and the neurons postsynaptic of these PNs in the brain center for encoding innate valences (the Lateral Horn, LH). We used the previously characterized neurons of the learning center, the MB: the KCs that sparsely represent stimulus identities; MB output neurons (MBONs) that represent learnt valences of stimuli; MB modulatory input neurons (MBINs, mostly dopaminergic, DANs) that provide teaching signals for learning; and their input neurons (MB feedforward neurons, MB-FFN (*31*)); MB feedback neurons (MB-FBNs that connect MBONs and MBINs (*31*)); and convergence neurons (CN) that integrate learnt and innate valences from the MB and LH (*34*). We also identified all presynaptic partners of the three output neuron types.

### Identification of all axons and dendrites in the brain

To better understand individual neuron morphology, we identified all axons and dendrites. In *Drosophila*, axons and dendrites contain the majority of a neuron’s presynaptic and postsynaptic sites, respectively, and are separated by a linker domain devoid of synapses. All linker domains were identified using synapse flow centrality (*40*). This data was manually proofread, and an axon-dendrite split point was placed for each neuron. Based on this analysis, we determined that 95.5% of the brain (2,421 neurons) are polarized with an identifiable axon and dendrite, 0.5% (13 neurons) are unpolarized with no definable axon, and 4.0% (102 neurons) were immature (Fig. 2A). These immature neurons were not developmentally-arrested SU neurons that differentiate into adult neurons (*67*). Their nuclei were not heterochromatin-rich like SU neurons, despite their general lack of arborization or synaptic sites. It is likely that these immature neurons started to differentiate, but are still in the process of neurite outgrowth and polarization. This population includes 78 immature KCs as previously reported in the memory/learning center (*25, 68*). However, there were also 24 non-KC immature neurons, revealing some neurogenesis of larval neurons outside of the MB.

**Fig. 2:**
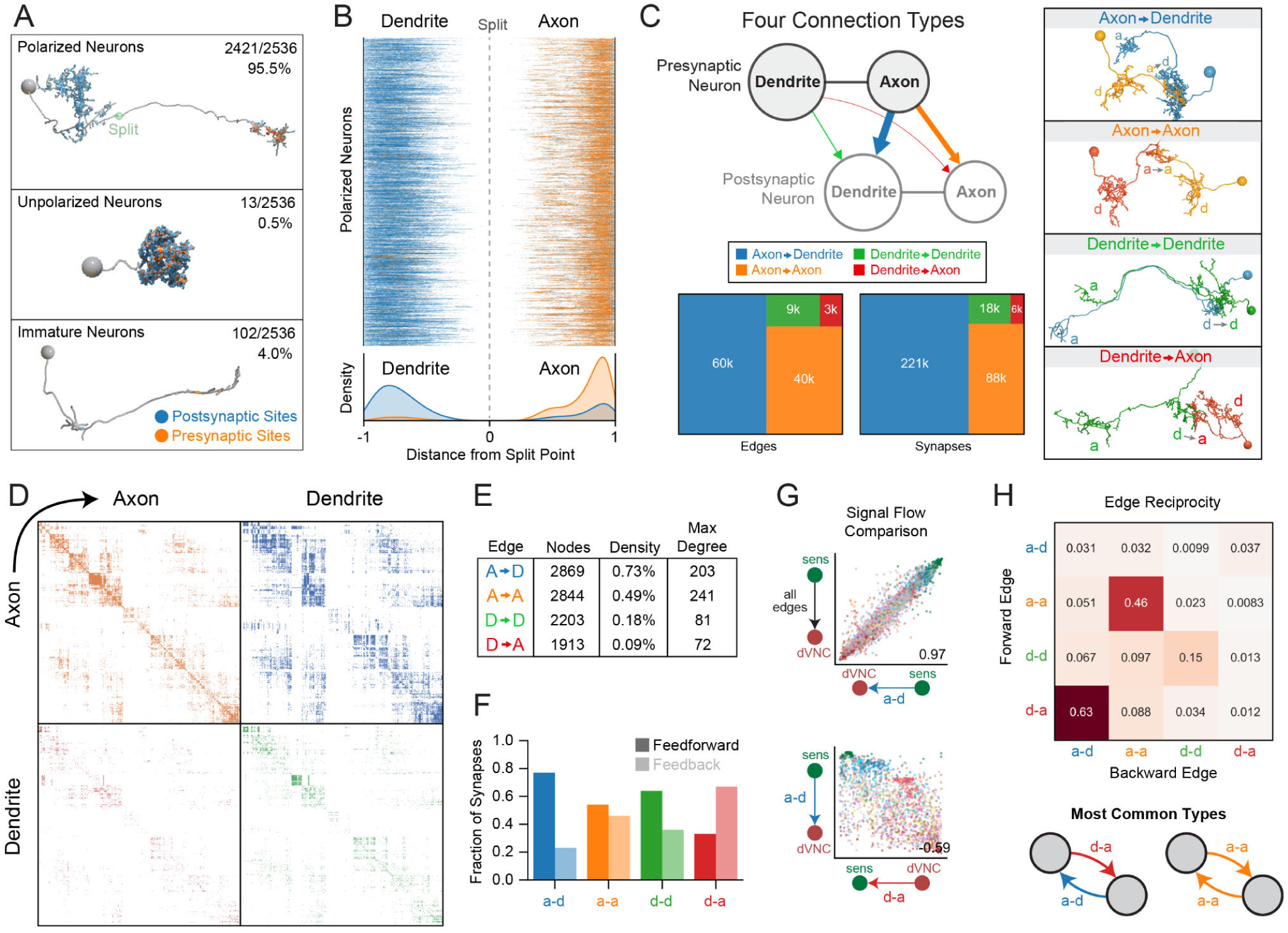
Identification of all brain axons and dendrites revealed four connection types. (**A**) Axons and dendrites were identified in all brain neurons. >95% of neurons contained fully differentiated axons and dendrites (i.e. were polarized), but a few unpolarized neurons and immature neurons were identified. (**B**) Presynaptic sites (orange) and postsynaptic sites (blue) in all polarized brain neurons, displayed as normalized distances from the axon-dendrite split point. Axons contained mostly presynaptic sites, while dendrites contained mostly postsynaptic sites, but pre- and postsynaptic sites were observed in both compartments. (**C**) Synaptic connections were categorized as axo-dendritic (a-d), axo-axonic (a-a), dendro-dendritic (d-d), or dendro-axonic (d-a). A-d connections were the most numerous, but many non-canonical connections were observed. Examples of each connection type are depicted in the right panel. (**D**) Adjacency matrices displaying all connection types between brain neurons. Each quadrant represents a different connectivity type between each presynaptic neuron (row) and postsynaptic neuron (column) in the brain. (**E**) Graph metrics for subgraphs comprising each connection type: number of nodes participating in each connection type, graph density (number of connections observed / all possible connections), and max degree (maximum number of connections from a single neuron). (**F**) Fraction of feedforward and feedback synapses per connection type, defined based on anterograde or retrograde direction compared to the overall neuron sorting from sensory to output (Fig. S5E-F). (**G**) Comparison of the direction of information flow in the indicated connection types. Individual neurons in each graph type were sorted using the signal flow algorithm (see Methods) and the correlation between these node sortings was quantified. A-d sorting best matched the summed graph sorting (all edge types together). The d-a sorting was negatively correlated with a-d (−0.59). (**H**) Edge reciprocity between different edge types, i.e. fraction of forward edges that were coincident with different backward edge types. The most overrepresented reciprocal edge types were d-a/a-d and a-a/a-a, while other types were uncommon.

All polarized neurons segregated pre- and postsynaptic sites within axons and dendrites, respectively (Fig. 2B). However, despite this general feature, we observed that axons contained postsynaptic sites and dendrites contained presynaptic sites. Thus, axons can be directly modulated by other neurons (*69*) and dendrites can directly synapse onto other neurons, which has been previously observed in the fly olfactory system (*45*) and in the visual system in mouse (*70*). We now report that this is a common phenomenon throughout the brain.

### Four connection types: axo-dendritic, axo-axonic, dendro-axonic and dendro-dendritic

While axo-dendritic connections are well established in the literature, other non-canonical interactions such as axo-axonic connectivity (*71–73*) and dendritic output (*70, 74–77*) have been observed but are not as well studied and their prevalence was unknown. We therefore identified all axo-dendritic (a-d), axo-axonic (a-a), dendro-dendritic (d-d), and dendro-axonic (d-a) connections in the brain. We found that the majority of connections are either a-d or a-a, however there are still a large number of d-d and d-a connections (Fig. 2C).

The connectome can be thought of as four graphs (Fig. 2D), where nodes represent individual neurons and each graph contains edges between these nodes, which are either a-d, a-a, d-d or d-a. Graph metrics were quantified for each graph type (Fig. 2E). We found that the axo-dendritic graph had the highest density and node participation (i.e. the most neuron-neuron connections and highest number of neurons participating in a-d connectivity), while the axo-axonic graph had the highest max degree (i.e. the max number of synaptic partners observed in an individual neuron). These results suggest that there are fundamental network differences between connection types. We next wondered whetherneurons were connected by just one edge (singleton) or multiple edge types (multiplexed). We found that the majority of edges were singletons, connecting neuron partners in only one way (a-d, a-a, d-d, or d-a). However, we also observed many multiplexed edges (Fig. S5D). The most common examples were a-d / a-a edges, however many different combinations were observed, including rare combinations of three or four edge types between the same partner neurons. We found that all multiplexed edge type combinations were observed more often than expected by a null model, although multiplexed edges were still less frequent than singleton ones (a-d: 93% singleton; a-a: 89% singleton; d-a: 79% singleton; d-d: 76% singleton). Four-edge connections were mostly found in local neurons (66%) (LNs, i.e. neurons involved in local processing in a specific neuropil) of the antennal lobe (*45*) and visual neuropils, while three-edge connections were found dispersed throughout the brain with a focus in local neurons (17%) and pre-descending neurons (14%). Multiplexed connections may grant presynaptic neurons post- and presynaptic regulatory control of their partner, as has been observed in a-d/a-a triad motifs in mammals (*78*). It will be interesting to explore the roles of these non-canonical connections in future experimental studies.

### Numerically strong connections are reproducible across brain hemispheres

We investigated the distribution of edge strengths for each connection type (Fig. S5A, B). We found that axo-dendritic connections displayed the highest fraction (a-d: 23%) of strong synapse edges (≥5 synapses between the same neuron pair). However, there were also strong connections among non-canonical connection types (a-a: 10%, d-d: 10%, d-a: 3%). As observed in both invertebrates (*10, 79*) and the mammalian cortex (*80, 81*), the majority of edges were weak (1 or 2 synapses) for all connection types (a-d: 60%, a-a: 75%, d-d: 80%, d-a: 91%). Yet, the majority of axo-dendritic synapses were contained in strong edges (61%; Fig. S5B). Overall, across all connection types, the brain comprised 17.9% strong edges and 65.8% weak edges. Strong edges (≥5 synapses) contained 54% of synaptic sites, while weak edges (1 or 2 synapses) contained 28% of synaptic sites between neurons. Based on these results, we can confirm that the connectome contains “a skeleton of stronger connections… immersed in a sea of weaker ones” (*80*); however, this statement should be tempered with the observation that most synaptic sites are contained within strong edges and therefore most developmental effort is spent on these stronger connections.

We next investigated edge symmetry across the two brain hemispheres. We found that edge strength correlated with interhemispheric symmetry (Fig. S5C); if there was a strong edge between two neurons in one hemisphere, a homologous edge in the opposite hemisphere was usually found. The four edge types (a-d, a-a, d-d, d-a) displayed similar patterns: weak edges were mostly asymmetrical while strong edges were highly conserved between hemispheres. With edge strengths of at least 5 and 10 synapses, the majority of edges (>80% and >95%, respectively) were symmetrical across all edge types.

Our analysis revealed that the majority of connections in the brain are weak and non-reproducible (not symmetric between left and right hemispheres). However, the majority of synaptic sites in the brain are contained in the strong symmetrical connections. It is interesting to speculate that the numerically strong edges might be important for reproducible aspects of behavior, whereas weak non-reproducible edges might contribute to stochastic aspects of behavior and idiosyncratic differences between individuals.

### Distinct connection-types differentially contribute to feedforward and feedback pathways

We devised a methodology to assay the contribution of different edge types to either feedforward or feedback signal throughout the brain. We developed a new algorithm, Walk-Sort (see Materials and Methods), which uses random walks through the network based on user-defined start and stop points to determine the relative position of a neuron within this particular flow of information. We applied Walk-Sort to the summed graph (with all edge types combined, a-d, a-a, etc.) to sort nodes according to the flow from sensory to descending neurons, and then categorized edges in the resulting sorted graph. We defined feedforward edges as those connections which project from a higher-ranked (closer to sensory periphery) to a lower-ranked (further from sensory periphery) neuron, while feedback edges are pointed in the opposite direction, connecting neurons lower in the sorting with those closer to the sensory periphery. We found that the a-d graph displayed the most feedforward; a-a and d-d graphs displayed roughly equal feedforward and feedback; while the d-a graph displayed a bias towards feedback edges (Fig. 2F; Fig. S5E, F).

We next compared how information flowed through the four networks without assumptions about the brain input-output axis. We utilized the signal flow algorithm (*82, 83*) because it sorts neurons within each graph according to the overall information flow without explicitly defining sensory and output neurons (required in Walk-Sort). We compared these signal flow sortings between edge types (Fig. 2G, Fig. S6). We found that the sorting of a-d graph best matched the summed graph (graph with all edge types combined), i.e. sorted from sensory periphery to brain output neurons. The a-a and a-d graphs displayed a similar flow from sensory to output, despite the details of the sorting being different (Spearman’s rho correlation coefficient = 0.44 between the signal flow sorting of the a-a and a-d graphs). Interestingly, the d-a graph resulted in a sorting that was the inverse of the a-d graph (Spearman’s rho correlation coefficient = −0.61), i.e. starting at brain output neurons and ending at the sensory periphery. We found that the majority of d-a edges were the inverse of a-d edges (i.e. there was a high edge reciprocity; Fig. 2H), which explains the inverse relationship between these graphs. Such reciprocal a-d/d-a connectivity could be important for input regulation, as suggested in mammals (*69*).

We also observed that a-a edges displayed high edge reciprocity, meaning many neurons engaging in a-a connectivity displayed reciprocal loops. Note that because a-a connections are directional, such reciprocal loops were not guaranteed to occur. Consistent with this finding, previous studies have observed a-a reciprocality between KCs (*25, 26*), between KCs and MBINs (*25, 26, 72*), and between olfactory PN axons in the adult fly (*73*). A-a connections have been shown to gate activity (*84*), implement divisive normalization (*85*), or result in reciprocal positive feedback loops (*72*).

### Hierarchical clustering reveals 90 connectivity-based brain neuron types

Next, we wanted to subdivide brain neurons into types based on their synaptic connectivity. Neuron types are often defined based on gene expression (*86–88*), morphology (*10, 89*), function (*28, 31, 90–92*), or a combination of features (*10, 13, 93*). These properties are likely all correlated with synaptic connectivity which is defined by the genome, requires appropriate neuronal morphology and is, in turn, necessary to support specific computations and behaviors (*55, 94, 95*). Thus, an unbiased clustering of neurons based on their synaptic connectivity could potentially reveal both morphologically and functionally defined neuron types.

We used the graph structure of all four connection types to spectrally embed all brain neurons in a shared space and clustered them using this representation (see Methods). This resulted in nested sets of clusters that can be tuned based on the desired granularity, from large groups of neurons to 90 fine-grained cell types (Fig. 3A; Fig. S7). In contrast with community detection algorithms that assume dense connectivity within communities and may reveal entire processing modules rather than individual neuron types, our clusters are not necessarily composed of groups of neurons which communicate more densely within a cluster (*96*). Instead, our clustering strategy groups neurons with very similar connectivity to other neurons even if little direct intracluster connectivity was present—for example, olfactory PNs from the antennal lobe which are known to function as parallel input channels and whose activity is regulated as a group (*45*). Our clustering methodology was also designed to ensure that left-right homologous neurons are placed in equivalent left-right hemisphere clusters, allowing straightforward subsequent analysis. To confirm that this connectivity-based clustering reveals neuron types that share other attributes besides connectivity we asked whether the morphology and function of neurons within the same cluster were similar. Indeed, we found the morphology of neurons within clusters was remarkably similar (mean within-cluster NBLAST score of 0.82 ± 0.16 SD), even though clustering was based solely on connectivity and no morphological data was used (Fig. 3B; Fig. S7A, B). Furthermore, neurons with similar known functions were usually found in the same or in related clusters (e.g. clusters of olfactory PNs, clusters of KCs, cluster of MBINs, clusters of MBONs, clusters of MB-FBNs, etc.; Fig. 3A, Fig. S7D).

**Fig. 3:**
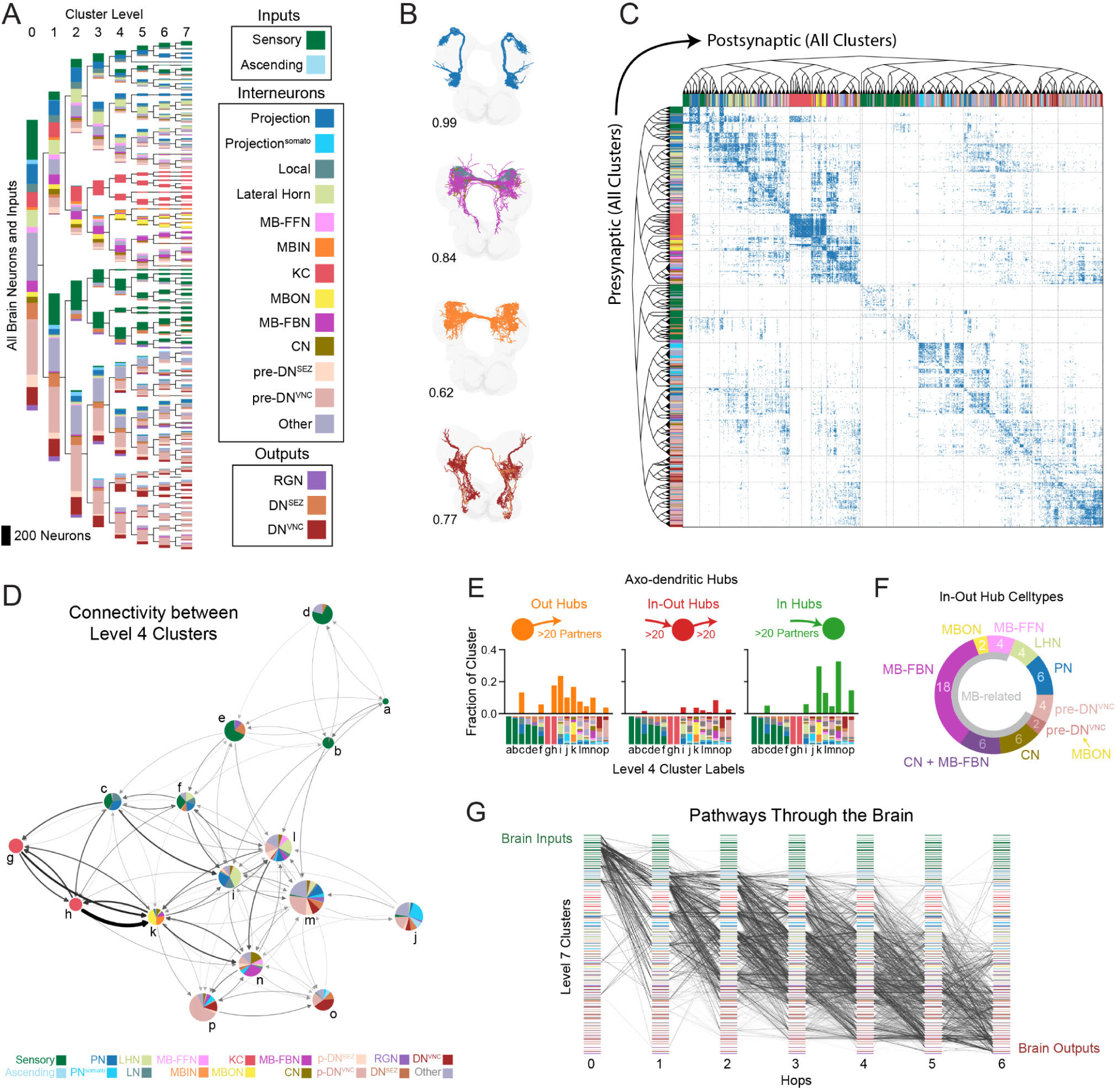
Hierarchical clustering and analysis of brain structure. (**A**) Hierarchical clustering of brain neurons using a joint left-right hemisphere spectral embedding based on connectivity. Clusters were colored based on cell classes (Fig. 1G, Fig. S4), but this information was not used for clustering. Clusters were sorted from SNs to DNs using the Walk-Sort algorithm. (**B**) Example clusters with intracluster morphological similarity score using NBLAST (see Methods). Most clusters displayed remarkable morphological similarity despite being clustered based on connectivity (Fig. S7A, B). (**C**) Adjacency matrix of the brain sorted by hierarchical cluster structure with color-coded neuron classes. (**D**) Network diagram of level 4 clusters displays coarse brain structure. Colored pie charts display cell types within clusters; size indicates number of neurons in each cluster. Arrow thickness is scaled to number of synapses between groups. (**E**) Fraction of a-d hub neurons in level 4 clusters. Cell types of each cluster are depicted as x-axis marginal plot and annotated to match clusters in (D). Hubs were defined as having ≥20 in- or out-degree (≥20 presynaptic or postsynaptic partners, respectively; based on the mean degree plus 1.5 standard deviations). (F) Cell classes of in-out hubs (a-d). The majority of neurons were downstream or upstream of the memory/learning center (gray semi-circle, *MB-related*). Note that *CN + MB-FBN* indicates neurons that were both CNs and MB-FBNs. One pair of pre-DN^VNC^ neurons received direct MBON input. (**G**) Pathways from SNs to output neurons with 6 or fewer hops, using a pairwise ≥1% input threshold of the a-d graph. Plot displays a random selection of 100K paths from a total set of 3.6 million paths.

The connectivity within and between all clusters is displayed in Fig. 3C. Many (but not all) clusters displayed strong intragroup connectivity and shared output to similar postsynaptic clusters. A coarser granularity can be also selected (Fig. 3D) and used to explore connectivity between larger groups of related neuron types. Altogether, we present a hierarchical clustering of the brain, which is robust across multiple independent metrics and designed to facilitate downstream analysis. These clusters represent connectivity-based cell types and are also internally consistent for both morphology and known function.

### The majority of brain hubs are pre- or postsynaptic to the learning center

Hubs are thought to play key roles in brain computations and behavior (*97–99*). We therefore identified brain hubs for all connection types. In order to focus on the strongest hubs, reproducible across hemispheres, we filtered the graph to include only strong edges observed in both hemispheres (using a ≥1% input threshold, see Methods). Brain hubs were defined as having ≥20 pre- or postsynaptic partners, respectively i.e. an in- or out-degree of ≥20 (this threshold is the network mean plus 1.5 standard deviations (SD)). We distinguished between in-hubs (over the in-degree threshold), out-hubs (over the out-degree threshold), and in-out hubs (over both thresholds). Using these criteria, we identified 506 a-d, 100 a-a, 10 d-d, and 8 d-a hubs (Fig. 3E, Fig. S8). A-d out-hubs were often observed in clusters closer to the sensory periphery, notably PNs (31%), while a-d in-hubs were more often closer to output clusters, including pre-output and output neurons (41%). The majority (73%, 19/26 pairs) of a-d in-out hubs were postsynaptic to the learning center output neurons (MBONs) and/or presynaptic to its modulatory neurons that drive learning (MBONs, CNs, MB-FBNs, MB-FFNs, and one pre-DN^VNC^ pair postsynaptic to MBONs; Fig. 3F). Several in-out hubs (23%, 12 pairs) were convergence neurons (CNs), receiving input from both the MB, that encodes learnt, and the lateral horn (LH), that encodes innate values (*31, 92*). One such in-out hub is the CN-MBON-m1, shown to functionally integrate learnt and innate values and bidirectionally control approach and avoidance (*34*). Together, these findings suggest that many of the brains’ in-out hubs may play a role in computing predicted values of stimuli (based on both learnt and innate values) and in regulating actions and/or future learning (via feedback to MB modulatory neurons).

### Identification of all brain local neurons

Brain neurons are often divided into local neurons (LNs), involved in local processing within a specific brain neuropil or layer, and PNs, which carry information to other brain regions. To systematically identify all brain LNs, we developed two connectivity-based definitions (Fig. S9A, B). Type 1 LNs provided a majority of their output to neurons in their sensory layer (defined by the number of hops from SNs of a particular modality), and/or to the sensory layer directly upstream of them (Fig. S9A). Type 2 LNs received a majority of their input and sent most of their output to any sensory layer, to which it did not belong (Fig. S9B). In this way, we identified all previously published LNs (*25, 45, 46*) and many new putative LNs (Fig. S9C, D). We then defined all 2nd order PNs by exclusion, i.e. all neurons that were not local but were directly downstream of SNs (Fig. S9E). Non-LN neurons that are higher order (i.e. not directly downstream of SNs) are usually termed output neurons from a specific neuropile (*12, 25, 90, 100*) rather than PNs, but we refrain from labeling them in a specific way and leave them undefined, as non-LNs. Although our LN definitions were connectivity-based, they provided results that matched morphological expectations. Namely, the Euclidean distance between the axon and dendrite of local neurons was small, while for PNs the axon-dendrite distance was large (Fig. S9D-F). Interestingly, we also found that LNs engaged in more noncanonical connectivity than PNs, including a-a, d-d, and d-a connections (Fig. S9G), perhaps allowing LNs to regulate multiple aspects of activity in both the axon and dendrite.

Interestingly, the majority of LNs (98 neurons) that meet the above definition were either 2nd order neurons directly downstream of SNs (i.e. one-hop from SNs) or 3rd order neurons (2-hops downstream of SNs, Fig. S9C). A very small number of 4th-order LNs were also identified (6 neurons, Fig. S9C-D). Two of the three pairs were pre-DN^VNC^ neurons and one was downstream of neurons that integrate learnt and innate valence, suggesting some level of local processing in the pre-DN^VNC^ layer and in post-MB layer. Overall, progressively fewer LNs were found further from the sensory periphery, perhaps suggesting that computations in higher-order neuropils are less local, involving integration and communication with a diversity of neuron types (rather than from a single sensory processing layer).

### Identification of all brain sensory pathways

Next, we systematically characterized brainwide pathways from distinct types of SNs to all other brain neurons. We note that, for the remainder of the paper, we will focus our analysis on a-d connections because they are the most abundant and best understood in terms of functional effects. We generated all possible a-d pathways from brain input neurons to all other brain neurons and ending at output neurons in fewer than 6 hops (Fig. 3G). We classified input neurons based on their known sensory modalities. Olfactory (*45*), gustatory (*47, 66, 101*), thermosensory (*48, 102*), visual (*46*), gut (*47, 103, 104*) and respiratory state SNs (*105*) project directly to the brain. Somatosensory ANs from the nerve cord received direct or indirect input from mechanosensory (*29, 38, 106*), nociceptive (*38, 107–109*) and proprioceptive SNs (*110, 111*) (Fig. S2, Table S1) and projected to the brain.

We identified all 2nd-, 3rd-, 4th-, and 5th-order brain neurons downstream of each input modality (Fig. 4A-C). For the purpose of this analysis, we defined the order of a neuron according to its lowest order input from any input neuron type. However, we note that neurons can receive multipath input from the same input neuron type, via distinct paths of different lengths (e.g. they can be both 2nd- and 3rd-order, etc.). Many brain neurons (539; 21%) were 2nd order, but the majority of brain neurons (1,403; 55%) were 3rd order (received input from a SN in two-hops). A considerable number were 4th order (386; 15%), but only 12 neurons (<1%) were 5th order (Fig. 4C). Note that 204 brain neurons (8%) were either immature or received only input from neurons in the SEZ of unknown modality and were therefore not categorized. Thus, of those neurons analyzed, no brain neuron was more than 4-hops removed from at least one input neuron and the vast majority were only 2- or 3-hops removed.

**Fig. 4:**
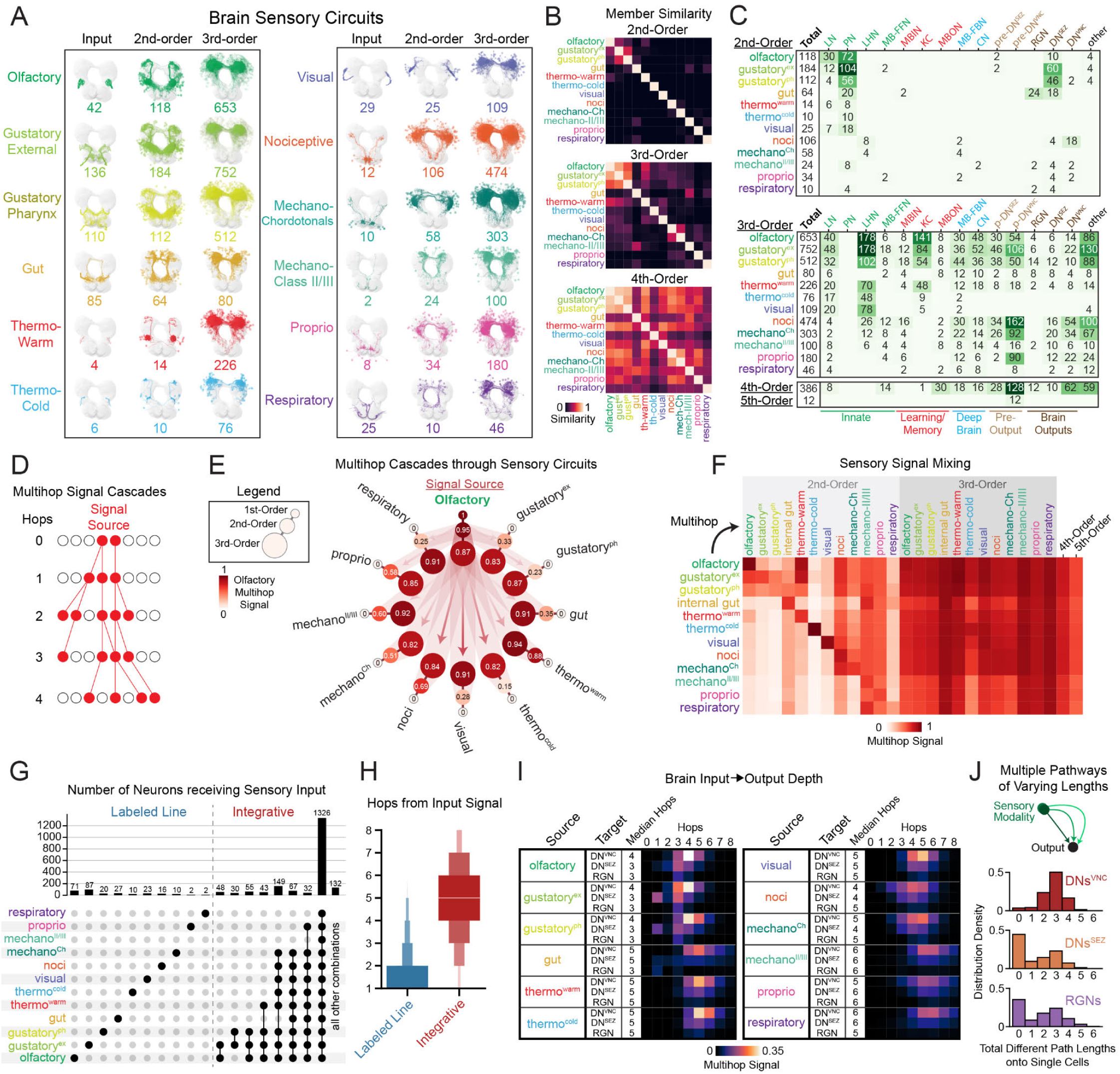
Multimodal sensory integration across the brain. (**A**) Morphology of brain sensory circuits, identified using multihop a-d connectivity from SNs or ANs. (**B**) Neuron similarity across sensory circuits using the Dice Coefficient. Most 2nd-order neurons were distinct between modalities, while 3rd-order neurons were more similar and 4th-order neurons displayed widespread similarity. (**C**) Cell classes in each sensory circuit. Note that row identities are not mutually exclusive. (**D**) Schematic of a multihop signal cascade, which probabilistically propagates signal from a user-defined source and endpoint using synaptic weights between neurons along the path. Multiple iterations are performed to determine how many times cascades visit particular neurons. Signal cascades were based on a-d connectivity throughout this study. (**E**) Signal was assessed in all sensory circuits after running cascades from olfactory SNs to output neurons (1000 independently run cascades, see Methods for normalization details). Olfactory signal traveled robustly to all 3rd-order neuron types and to some 2nd-order neurons. (**F**) Signal cascades from each sensory modality (rows) to all sensory circuit layers (columns). Multimodal integration is widespread in 3rd-, 4th-, and 5th-order neurons. 4th- and 5th-order neurons were not divided by modality due to extensive overlap of their members across modalities. The first row displays the same data from (E). (**G**) Combinations of sensory integration at the single-cell level (UpSet plot). Neurons were considered to receive sensory input when visited in most cascade iterations. The majority of brain neurons integrate from all sensory types, but a few neurons integrated from only one sensory modality (labeled line) or from a particular combination of modalities. This analysis was performed on fully differentiated neurons in the brain (2412 neurons). 315 neurons did not receive an over-threshold signal; many of these neurons displayed dendrites in the SEZ and therefore received weak signal from brain neurons. (**H**) The distance from sensory input in labeled line or integrative cells from was quantified. Labeled line neurons were generally very close to the sensory periphery, while integrative cells were deeper in the brain. (**I**) Signal cascades from sensory modalities to brain outputs (row normalized), including distribution of hops from sensory to output. The median sensory to DN^VNC^ path lengths were between 4-6 hops, with a maximum value of 8 hops; we consider this the maximum brain depth. (**J**) Number of pathways of different lengths from individual sensory modalities to individual output neurons. Only pathways contributing substantial cascade signal per hop were considered (>0.1 multihop signal). Individual sensory modalities sent multihop signal to output neuron types through pathways of multiple different lengths. Note that not all DNs^SEZ^ and RGNs received signal from each sensory modality (thus the peaks at 0).

We found that most 2nd-order neurons received direct input from a single SN type (Fig. 4B), with some exceptions, including olfactory local neurons that also received from gustatory and thermo^warm^ SNs, as previously reported (*45, 48*). 3rd-order neurons were more often shared across modalities and, by the 4th-order, most neurons were shared across modalities (Fig. 4B). However, we note that even neurons that are exclusively 2nd or 3rd order for one modality, can receive input from other modalities via longer paths (discussed in a future section).

Most sensory modalities exhibited a large expansion of neuron numbers in 3rd-order, compared to 2nd-order layers (Fig. 4A, Fig. S10A, Table S2), indicating prominent divergence, i.e. they broadcast their signals to very many different downstream partners. Generally, the number of neurons downstream of 2nd-order PNs (divergence) was higher than the number of PNs upstream of the 3rd-order neurons (convergence). Nevertheless, convergence was also prominent, with most 3rd-order neurons receiving input from multiple 2nd order PNs.

The expansion-contraction architecture is thought to be an adaptation of learning circuits to sparsify dense neural codes and enhance stimulus discrimination by increasing the dimensionality of representation (*112–114*). Interestingly, in the 1st instar larva, we observed a similar amount of divergence and convergence onto 3rd-order LH (innate center) and MB (learning center) neurons, suggesting both 3rd-order neuropils may discriminate between stimuli to a similar extent. Thus, olfactory PNs synapsed onto 15 ± 5 KCs and 23 ± 10 LHNs (average ± SD). KCs and LHNs received input from 4 ± 3 and 4 ± 5 uPNs, respectively. In the adult, the olfactory signal divergence onto KCs is 2.5 times greater than onto LHNs (*100*). During larval life, the number of KCs increases 3-fold, whereas the number of LHNs remains constant. Thus, in a 3rd instar larva divergence of information in the MB compared to the LH may be greater, potentially enabling better odor discrimination (*114*).

### Sensory information can reach output neurons within one to three hops

We investigated the cell type identities of neurons at different processing layers, i.e. at different hops from SNs or ANs (2nd, 3rd, 4th and 5th order neurons) within each sensory circuit (Fig. 4C, Fig. S10A). We found that sensory information reached all cell classes within a couple hops. A surprising percentage of brain output neurons were 2nd order, i.e. postsynaptic (one-hop) of SNs or ANs (Fig. 4C; *DNs^VNC^*: 16%, *DNs^SEZ^*: 54%, *RGNs*: 61%), or 3rd order, i.e. two hops from SNs or ANs (*DNs^VNC^*: 54%, *DNs^SEZ^*: 37%, *RGNs*: 11%). The remaining 30% of DNs^VNC^, 9% of DNs^SEZ^ and 28% of RGNs were 4th order (three-hops from SNs/ANs, Fig. 4C). Thus, all output neurons can receive sensory information within a maximum of three-hops. However, we found that while these direct (one-hop), two-hop, or three-hop connections represent the shortest paths to output neurons, most output neurons also received longer multihop input from SNs.

The highest-order neurons in the brain (5th-order) were not output neurons, but 12 pre-output neurons, presynaptic to DNs^VNC^. These neurons received input from and output to other pre-DNs^VNC^ (the most numerous group of 4th-order neurons) and share some upstream and downstream partners, suggesting complex, multilayered connectivity between pre-DNs^VNC^ (Fig. S11). This suggests that, even though DN^VNC^ neurons can receive sensory input in very few hops, they also receive the most processed information in the brain via longer paths. We observed multiple parallel pathways from each sensory modality to DNs (Fig. S12A, B). However, we also found extensive connectivity between neurons within these parallel pathways, suggesting they likely form a distributed processing network (Fig. S12C). We found that a majority of pathways and the vast majority of individual neurons within paths were not unique to a particular sensory modality and were instead shared by multiple modalities (Fig. S12D, E).

We found that different sensory modalities targeted different types of output neurons (Fig. S10C, D). For example, gustatory and gut sensory signals targeted more DNs^SEZ^ than DNs^VNC^, whereas other modalities targeted more DNs^VNC^ than DNs^SEZ^. This is consistent with the proposed role of SEZ motor neurons in driving feeding behavior (*66, 115*) and VNC motor neurons in driving locomotion (*42, 43, 116*). Generally, sensory pathways to DNs^SEZ^ were shorter compared to pathways to DNs^VNC^. The majority of DNs^SEZ^ were 2nd-order (receiving direct inputs from SNs), whereas the majority of DNs^VNC^ were 3rd-order. This raises the possibility that general action selection may require more processing steps than feeding behavior.

### Brain output neurons receive the same sensory input via multiple paths of different lengths

While characterizing the shortest paths from SNs to output neurons, we observed that output neurons also receive sensory information via longer paths. We therefore wanted to systematically analyze both short and long pathways without a bias towards shortest ones.

For this purpose, we developed a computational tool, the signal cascade, that propagates multihop signals through the brain based on the numbers of connections between neurons at each hop (Fig. 4D; see Methods). This tool captures all pathways with reasonably strong connections along their length and not just the shortest ones. Signals can be started and terminated at pre-defined neurons to explore all pathways that link them. We note that this algorithm makes no assumption about the excitatory or inhibitory sign of connections, only about the likelihood of signal propagation from one neuron to the next. Throughout this paper, we will use brain output neurons as end points unless otherwise mentioned. In cascades started at SNs, we found that the signal generally reached DNs^VNC^ in 3-6 hops and rarely more than 8 hops, which we therefore considered the maximum depth of the brain. 5-hop pathways were shown to be functional in the larva (specifically, MD class IV neurons to MB DANs (*31*)), but no studies have yet functionally tested 6, 7, or 8-hop pathways. We therefore stop the cascades at either 8 or 5 hops, using 8-hops to not miss long paths and 5-hops to determine which aspects of architecture are apparent with a pathway-length for which functional connectivity has been confirmed.

Using signal cascades, we identified all multihop pathways between SNs or ANs and output neurons (Fig. 4I). Individual sensory modalities had different median pathway depths to output neurons. Overall, olfaction and gustation displayed the shortest pathways to output neurons, while the ascending somatosensory modalities displayed the longest. This is even more striking, because somatosensory ANs are already carrying processed sensory information and are themselves one- to three-hops removed from SNs (*29, 38*).

We found that output neurons received sensory inputs from the same modality via multiple paths of different lengths. For example, some paths from the same sensory modality reached DNs^VNC^ in 2 hops, while others displayed as many as 6 hops (Fig. 4I). This was also true on the single cell level, with the vast majority of individual DNs^VNC^ receiving multipath input. DNs^VNC^, on average, received input from at least three distinct pathways of three different lengths from individual sensory modalities (Fig. 4J). When DNs^SEZ^ and RGNs received sensory input, it was also mostly multipath. Thus, convergence of shorter and longer feedforward paths from individual sensory modalities onto single output neurons appears to be a general architectural feature of the brain feedforward circuits, for all sensory modalities and for all output neuron types.

### More than half of all brain neurons integrate information from all sensory modalities

We next investigated the multimodal character of sensory circuits and the brain as a whole, while taking into account both short and long pathways. We therefore started signal cascades at different sensory modalities and assayed which sensory circuits (i.e, which 2nd, 3rd, 4th or 5th-order neurons as defined based on shortest paths, from Fig. 4A) received these signals. For example, we assayed how much signal from olfactory SNs traveled to all other sensory circuits (Fig. 4E) and found it is integrated by some 2nd-order and most 3rd-order neurons of distinct sensory modalities.

To assess global sensory integration, cascades were initiated from each sensory modality and signal assayed at all sensory circuits (Fig. 4F). 2nd-order neurons displayed some multimodal mixing, while 3rd, 4th, and 5th-order neurons received robust input from many sensory modalities, suggesting most brain regions are multimodal. Many modalities converged at the earliest stages of sensory processing on 2nd-order PNs (Fig. 4F). Strikingly, only 35% of 2nd-order neurons were unimodal. However, this varied by sensory modality; for example, 100% of thermo-cold and 92% of visual 2nd-order neurons were unimodal, in contrast to 36% of olfactory and 6% of proprioceptive 2nd-order neurons. Only 4% of 3rd-order and no 4th- or 5th-order neurons were unimodal. Consistent with these findings, we observed direct connectivity between different sensory circuit types (Fig. S10B), providing avenues for multimodal mixing.

We next investigated multimodal integration for each brain neuron, independent of its location in sensory circuits. We started signal cascades from each sensory modality and reported the combinations of sensory input each neuron received (Fig. 4G, Fig. S10C). We found that a small minority of neurons (12%) received signal from only one modality, so-called labeled line neurons, while a majority of neurons were integrative (88%), receiving signal from multiple modalities. Surprisingly, the majority of neurons integrated information from all sensory modalities (62% of neurons). Most labeled line neurons were close to the sensory periphery (Fig. 4H). Based on these data, we postulate that true labeled line pathways are rare and, without comprehensive connectivity data, may only appear as such in a particular experimental context.

We also quantified sensory integration at the level of brain output neurons. We found that different output types displayed different patterns of integration (Fig. S10D, E). Most DNs^VNC^ integrated signal from all sensory modalities (80%), with only a few labeled line DNs^VNC^ (2%). DNs^SEZ^ and RGNs were more often labeled line (26% and 7%, respectively). These findings are consistent with previous publications demonstrating that larval feeding circuits utilize DNs^SEZ^ and RGNs (*47, 49, 65, 66*) that are close to the sensory periphery.

Finally, we analyzed sensory integration in MB DANs. DANs have been implicated in learning, motivation, and action-selection across the animal kingdom (*117–120*) and understanding the type of sensory information they receive is essential for understanding their function. DANs are known to receive a-d input from sensory systems that sense rewards and punishments (*31, 121–123*), but the extent to which they receive input from other modalities was less clear. Interestingly, we found that all DANs integrated input from all sensory modalities, including from those that normally sense conditioned stimuli in learning tasks (e.g. olfactory) and from proprioceptive neurons (Fig. S20A). In contrast to the DANs, other MB modulatory neurons, namely octopaminergic neurons (OANs) and MBINs with unknown neurotransmitter, were not as integrative: only 33% of OANs and 60% of other MBINs integrated all modalities. In summary, a comprehensive view of all sensory inputs into DANs revealed that they receive multi-hop input from all sensory modalities, suggesting that multimodal integration may be important for computing teaching signals that drive learning.

### Identification of multihop signal cascade hubs

Our analysis of multihop pathways suggests that signal propagation is not all-to-all, despite extensive multimodal integration. For example, some neurons do not receive signal from certain sensory modalities (even with 8-hop cascades). To systematically explore the extent to which signal from any neuron reaches other brain neurons, we initiated cascades from each homologous pair and asked what fraction of neurons is reached by their signal within 5- or 8-hops (Fig. S13A). On average, signal from any pair of SNs reached only 13% (± 20% SD) and 31% (± 30% SD) of brain neurons within 5- and 8-hops, respectively. Similarly, signal from any pair of brain neurons reached 16% (± 15% SD) and 41% (± 25% SD) of other neurons within 5- and 8-hops, respectively.

We wondered whether there are neurons whose signal reaches a greater fraction of neurons than others. We quantified the polysynaptic in- and out-degree of individual neurons, i.e. the number of upstream and downstream partners connected by 5-hop cascades. We identified cascade hubs (Fig. S13B-D) using the same definition as for direct-connectivity hubs (Fig. 3E), namely all neurons with more polysynaptic partners than the network mean plus 1.5 SD. We found many cascade in- (480 neurons) and out- (122 neurons) hubs, but only a single pair of in-out hubs. Cascade in-hubs received input from 59% (± 9% SD) of the brain, cascade out-hubs output to 46% (± 4% SD) of the brain, and the cascade in-out hub received from 46% and output to 49% of the brain. There was a fair amount of overlap between cascade and direct-connectivity hubs: 56% of direct-connectivity hubs were also cascade hubs (73% of in-hubs, 23% of out-hubs, Fig. S13C). Cascade out-hubs were either part of the sensory periphery (PNs, LNs) or learning/memory center (KCs, MBONs), while cascade in-hubs belonged to a variety of cell types (Fig. S13D), including MB-FBNs, CNs, pre-DN^VNC^, and DNs^VNC^, MBINs, and MBONs. Interestingly, the only cascade in-out hub was CSD, a serotonergic neuron (*124, 125*), thought to single-handedly regulate hunger-based internal state (*126*).

### Identification of all ipsilateral, bilateral, and contralateral neurons

A fundamental property of brains is their bilateral symmetry, i.e. the presence of two hemispheres. To better understand how the brain hemispheres interact, we identified all neurons that engaged in interhemispheric communication via contralateral edges (Fig. 5A, B) and categorized them based on their axonal and dendritic projections.

**Fig. 5:**
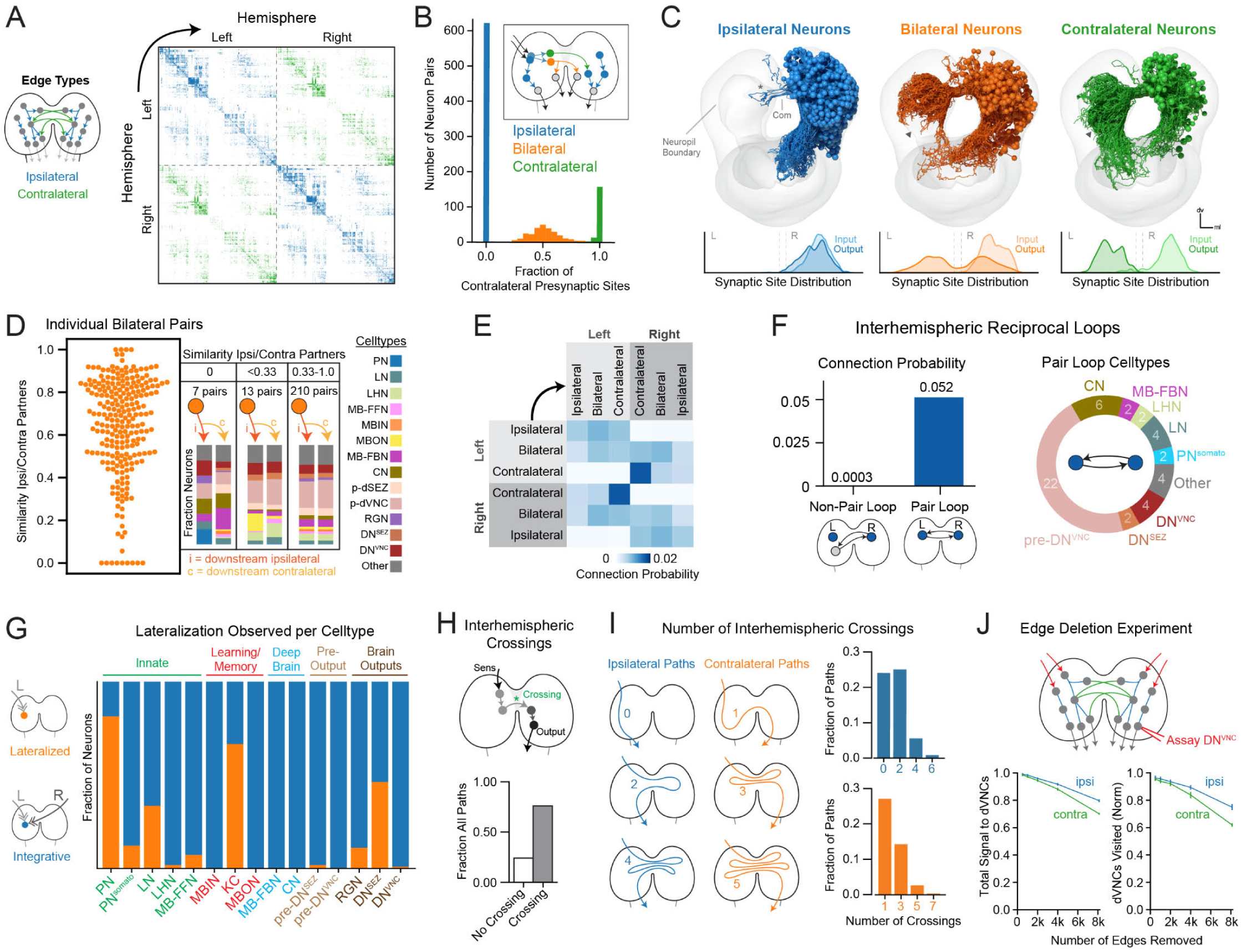
Characterization of interhemispheric communication by bilateral and contralateral neurons. (**A**) Adjacency matrix depicting connectivity between left and right brain hemispheres, sorted within each hemisphere by the cluster structure (Fig. 3A, Fig. S7D). Blue edges indicate ipsilateral connections, green edges indicate contralateral connections. (**B**) Fraction of contralateral a-d presynaptic sites per neuron. The trimodal distribution corresponded to ipsilateral, bilateral, and contralateral axon neuron types. (**C**) Morphology of ipsilateral, bilateral, and contralateral axon neurons with a-d synaptic distribution (right-side neurons displayed so that contralateral arbors can be seen). Most dendrites were ipsilateral, but there were some rare neurons with bilateral or contralateral dendrites (Fig. S14C). (**D**) Most individual bilateral axon neurons synapsed onto homologous neurons in the left/right hemispheres, as indicated by high cosine similarity of their a-d connectivity to ipsilateral and contralateral downstream partners (*left*). A bilateral axon neuron received a cosine similarity score of 1 if it synapsed onto homologous neuron pairs in both hemispheres. A score of 0 indicates that none of its ipsi- and contralateral partners are homologous pairs. A minority of bilaterals communicated with different left- and right-hemisphere neurons. Three bins of cosine similarity values and the cell type memberships of the downstream partners are displayed (*right*). (**E**) Connection probability between left and right cell types using a-d edges. Bilateral neurons were equally likely to connect to each hemisphere, but with a bias towards bilateral or contralateral axon neurons. Contralateral axon neurons displayed a preference to synapse onto other contralateral neurons in the opposite hemisphere. (**F**) Interhemispheric reciprocal loops. Neurons with contralaterally projecting axons tended to form reciprocal a-d connections between left and right homologous pairs, in contrast to the low probability of forming loops with non-homologous neurons. The cell classes involved in pair loops are reported on the right. (**G**) Quantification of sensory signal lateralization per cell class. Signal cascades were generated from left- or right-side SNs independently; neurons were considered to receive signal if they were visited in more than half of simulated cascades. All neurons that received signal from both hemispheres were classified as integrative (blue), while those that received signal from only one hemisphere were classified as lateralized (orange). (**H**) Interhemispheric crossings were observed in a majority of a-d pathways through the brain (SNs to DNs). (**I**) Interhemispheric crossings in ipsilateral and contralateral a-d pathways. Ipsilateral pathways were defined as starting and ending in the same hemisphere, regardless of any interhemispheric crossings in between. Contralateral pathways started and ended on opposite hemispheres. Most ipsilateral paths contained interhemispheric crossings, while most contralateral paths displayed only one crossing. (**J**) Signal cascades were generated from left/right SNs simultaneously and total signal was assayed in DN^VNC^ output neurons. Deletion of random sets of contralateral edges resulted in a decrease in total a-d signal to DNs^VNC^ (*left*) and a reduction in the number of DNs^VNC^ that received signal at all (*right*). Metrics are normalized to results from the unmanipulated control graph.

We found three major populations of neurons based on their axonal projections: neurons with ipsilateral (60%), bilateral (24%), or contralateral (16%) axons (Fig. 5C). We found that the vast majority (98%) of neurons displayed ipsilateral dendrites (Fig. S14). A small set of neurons (1%) had contralateral dendrites as well as contralateral axons (Fig. S14C), such that their cell body and neurites were located in the opposite hemispheres. Interestingly, a small population of neurons (1%) had bilateral dendrites that extend into both hemispheres with either ipsilateral, bilateral or contralateral axons. These neurons were only observed in the learning center (MBONs) and brain output network (pre-DNs^VNC^, DNs^VNC^, DNs^SEZ^) (Fig. S15). Neurons with bilateral dendrites are well suited for integrating sensory drive from both hemispheres to promote non-lateralized behaviors, such as backwards crawling as observed in the MDNs, which display bilateral dendrites (*42*).

### Some neurons with bilateral axons target distinct partners in the two hemispheres

Neurons with bilateral axons output to both hemispheres, but do they communicate with homologous postsynaptic partners in both hemispheres? To answer this question, we calculated the cosine similarity between postsynaptic partners of individual bilaterally projecting neurons in the left and right hemispheres (Fig. 5D, left). We found that most bilateral neurons generally connected to homologous partners in both hemispheres, i.e. had high partner similarity scores, but there were some neurons that had low scores. We binned these neurons into three categories based on their partner similarity scores and analyzed their downstream partners further (Fig. 5D, *right*; Fig. S16).

Strikingly, we found 7 pairs of bilateral neurons with completely different postsynaptic partners on the ipsi- and contralateral brain hemispheres, and 13 pairs of bilateral neurons with mostly non-overlapping sets of ipsi- and contralateral postsynaptic partners (Fig. S16). All of these neurons had unilateral dendrites. Most asymmetric bilateral neurons synapsed onto pre-DNs or DNs in only one hemisphere, but not the other, or onto different DNs or pre-DNs across the two hemispheres. It is interesting to speculate that these neurons could be involved in controlling asymmetric motor patterns that require activation of different subsets of muscles on the left and right side of the body. Indeed, several DNs that receive input from asymmetric bilateral neurons (Fig. S16C) have presynaptic sites in thoracic and early abdominal segments, perhaps indicating a role in turning (*127–129*).

### Reciprocal contralateral loops

To better understand information flow between brain hemispheres, we asked how ipsilateral, bilateral, and contralateral neurons communicate with each other and calculated their connection probability (Fig. 5E). We found that ipsilateral neurons synapsed approximately equally onto ipsilateral, bilateral and contralateral neurons in the ipsilateral hemisphere. Bilateral neurons had a slight preference for bilateral and contralateral neurons in both hemispheres. Surprisingly, contralateral neurons displayed a striking preference for other contralateral neurons, both in terms of input and output. Individual contralateral neurons directly synapsed onto 3.4 other contralateral neurons on average (34% of their downstream partners), compared to ipsilateral neurons, which only synapsed onto 1.5 contralateral neurons on average (15% of their downstream partners).

Given the striking preference of contralateral neurons for each other and the fact that each contralateral neuron has a homolog in the opposite hemisphere, we wondered whether homologous left-right contralateral neuron pairs tended to directly synapse onto each other. To test this, we calculated the connection probability between homologous neuron pairs compared to non-homologous neurons. We found the connection probability onto a homologous contralateral partner was much higher than onto a non-homologous neuron (Fig. 5F). We identified 24 neuron pairs that engage in homologous pair loops (10% of contralateral and 2% of bilateral neurons; Fig. S17). More than half were pre-DNs^VNC^ or DNs^VNC^; a third were postsynaptic of the learning center outputs (MBONs) and/or provided feedback onto the MB DANs (Fig. S18). Many pair loops interacted amongst themselves, forming double loops or super loops between pair loops (Fig. S18B). Double loops and super loops occured between neuron pairs with relatively similar morphology and/or connectivity. One super loop involved four neurons downstream of the in-out hub, MBON-m1, that integrates input from other MBONs and from the LH (*92*) and computes predicted values of stimuli. This super-loop output onto pre-DNs^VNC^ and indirectly sent feedback onto MB DANs via MB-FBNs (Fig. S18C). The other super loop involved five neurons that output onto DNs^VNC^. Thus, the reciprocal pair loops, double and super loops appear to be prevalent in brain areas that potentially play a role in action-selection (downstream of MBONs and upstream of DNs^VNC^) and learning (upstream of MB DANs). Indeed, reciprocal connectivity and interhemispheric communication has been implicated in action selection (*29, 130*) and working memory (*131–133*) in a variety of organisms.

### Interhemispheric integration occurs across most of the brain

Our finding that 37% of brain neurons have contra- or bilateral axons suggests that the two hemispheres are heavily interconnected and their information could be integrated at many sites. To systematically investigate where interhemispheric convergence occurs, we generated signal cascades from either left- or right-side SNs and observed the resulting signal propagation through both brain hemispheres (Fig. S19A-C). We found that signal crossed to the opposite hemisphere within 2 hops and was robustly found in both hemispheres by 3 hops (Fig. S19A). We assessed simultaneous overlap between left- and right-side sensory signals to find interhemispheric integration sites. The cell types of all integrative ipsilateral, bilateral, and contralateral types were identified (Fig. S19B).

Next, we investigated the lateralization status of single cells across the brain (whether they integrate signals from only one or both hemispheres). We quantified the lateralization of each neuron based on the ratio of left and right signal they received via signal cascades (Fig. S19C). We found that a majority of the brain (81%) integrated signals from both left- and right-side SNs, while a minority (19%) received only same-side lateralized signal from their own hemisphere. We found very similar results using 5-hop cascades: 79% of neurons integrated signal from left and right-side SNs and 21% received only signal from their own hemisphere. When plotted on a per cell type basis (Fig. 5G), it became clear that most PNs were lateralized, suggesting that interhemispheric mixing primarily occurs in higher-order processing centers. A large fraction of DNs^SEZ^ also received lateralized input, while the vast majority of DNs^VNC^ did not (Fig. 5G). This is consistent with previous work suggesting that the SEZ may be involved in lateralized behavior, such as orientation events/turning (*134*).

### Interhemispheric crossings increase the number of pathways through the brain

Interhemispheric communication may increase the processing potential of the brain by opening up a larger pool of computational nodes and pathways. To test this hypothesis, we generated a comprehensive list of multihop pathways from brain input to output neurons (maximum 6 hops for computational expedience, using ≥1% input threshold). We quantified whether these pathways ended in the same hemisphere that they started in or whether they crossed between hemispheres. We found that 75% of pathways crossed between hemispheres at least once (Fig. 5H). More than 50% of pathways began and ended in the same hemisphere (but they may have crossed and crossed back). 25% of paths never crossed. We characterized the distribution of interhemispheric crossings for all sensory-to-output pathways and found that multiple interhemispheric crossings were surprisingly common (Fig. 5I). The majority of ipsilaterally-ending pathways engaged in multiple crossings (median of 2 crossings), while most contralaterally-ending pathways only crossed hemispheres once (although a large minority crossed 3 times).

We investigated the extent to which contralateral or ipsilateral edges were important for signal propagation from sensory to output neurons. We removed either ipsilateral or contralateral edges, ran signal cascades from both left- and right-side SNs simultaneously, and quantified how much signal reached brain output neurons. We found that excision of contralateral edges reduced the total amount of signal that reached DNs^VNC^, as well as the number of individual DNs^VNC^ that received strong sensory signal (Fig. 5J). Removing contralateral edges had a stronger effect than a similar ipsilateral edge removal. In contrast, removal of contralateral edges did not reduce signal to DNs^SEZ^ more than ipsilateral edge excision and ipsilateral edges seemed to play a bigger role for RGNs (Fig. S19D). Contralateral and ipsilateral edges were equally important to transmit signal to interneurons generally, other than DNs^VNC^ (Fig. S19D). Thus, contralateral paths may be bottlenecks for information flow from SNs to DNs^VNC^. Consistent with this idea, we found that 67% of a-d in- and in-out hubs (Fig. S7A) had either contra- or bilateral axons (19% of contralateral and 19% of bilateral neurons were in or in-out hubs).

We also compared how ipsilateral and contralateral edges connect cell types in the brain. We found that while patterns of connectivity appeared similar (Fig. S19E, F), sometimes contralateral edges provided categorically new types of connectivity (Fig. S19G).

Overall, contralateral edges increase the number of pathways through the brain, help transmit signal to DNs^VNC^, and provide new cell-cell connectivity not observed in the ipsilateral networks of each hemisphere. We speculate that the increased number of pathways provided by interhemispheric crossings could increase the processing power of the brain, by increasing the depth of the neural network and providing more steps of information processing.

### Analysis of brainwide pathways reveals a nested recurrent architecture

The dominant synaptic network of the brain comprised a-d connections (Fig. 2C), many of which provide feedforward signal from sensory to output systems (Fig. 2F). However, recurrence is an important feature of brain circuits (*31, 135–137*) and can improve computational power in artificial neural nets (*138*). We therefore characterized the reverse signal in the a-d network, from output neurons back towards the sensory periphery. To do this, we generated independent signal cascades starting at each level-7 brain cluster (Fig. 3A). Because these clusters were sorted from brain inputs to outputs, we could track the extent to which signals propagated up or down this brain structure to other clusters. We kept these cascades short (ending after 2 hops) to initially limit our analysis to the shorter paths of reverse signal and identify its lower bound. Cascade signal that traveled up the brain cluster structure towards the sensory periphery was considered backward, while the signal that traveled down the cluster structure towards the output neurons was considered forward (Fig. 6A). We found that robust forward and backward signal originated from nearly all brain clusters (Fig. 6B). We found that deeper brain clusters (closer to brain outputs) received mostly forward signal, while shallower clusters (closer to sensory periphery) received a mixture of forward and backward signal (Fig. 6C). Most brain clusters provided forward and backward signal to multiple other clusters simultaneously; this was observed even for single neurons within each cluster (Fig. 6D).

**Fig. 6:**
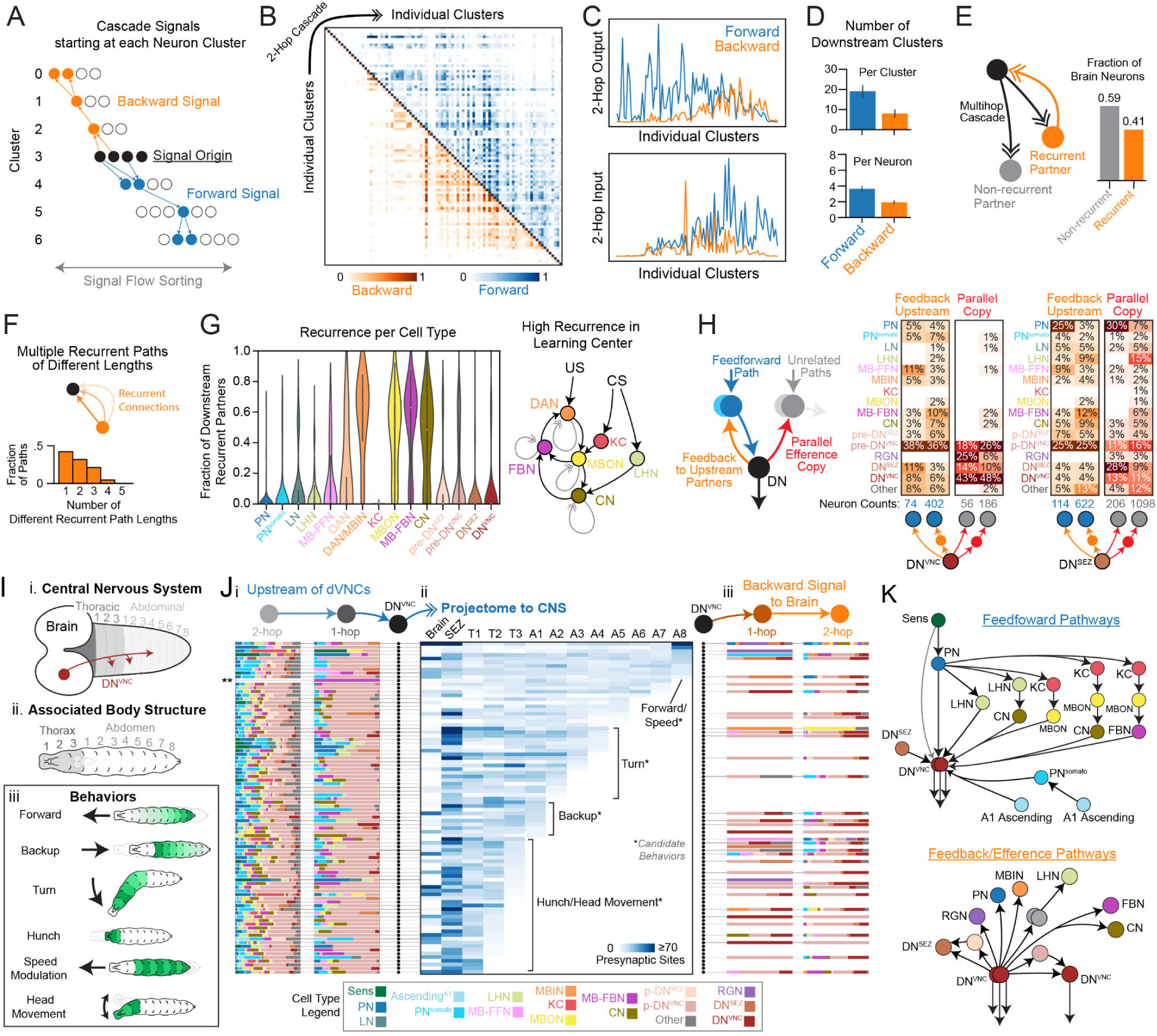
Comprehensive feedforward and feedback pathways through the brain. (**A**) Schematic of signal cascades starting from each cluster. Forward signals travel up the cluster sorting towards SNs; backward signals travel down the cluster sorting towards DNs. (**B**) Signal cascades originating at each level-7 cluster (along the diagonal) travel in both forward (above the diagonal) and backward (below the diagonal). Signal cascades were based on a-d connectivity and contained 2-hops maximum to restrict analysis initially to the lower bound of backward signal. (**C**) Quantification of forward and backward signal per cluster (along the x-axis), including cascade output signal (*top*) and cascade input signal (*bottom*). (**D**) The number of clusters or single cells that received cascade forward or backward signals from clusters or single cells within clusters, respectively. For the cluster to cluster analysis, non-negligible forward or backward signal (>0.05) was used. For the single cell analysis, the usual cascade threshold was used (receiving signal from a majority of cascade iterations). (**E**) Recurrence in brain neurons. Polysynaptic downstream partners of each brain neuron were identified with a-d cascades (up to 5 hops). Recurrent partners sent multihop signal back to the source neuron, forming a recurrent loop (*left*). 41% of brain neurons engaged in at least one such recurrent loop (*right*). (**F**) Quantification of recurrent pathways of different length between individual neurons. We found that recurrent signals usually utilized multiple pathways of different lengths. (**G**) Recurrence was quantified for each cell class. To do this, we quantified the fraction of individual neurons’ downstream partners that were recurrent for each cell type, indicating how many recurrent loops each neuron was involved in. On the right, a schematic of the most recurrent cell types in the brain and their relation to conditioned stimulus (CS) and unconditioned stimulus (US) during associative learning. Note, the MBIN category was split into OANs (octopaminergic neurons) and DAN/MBIN (dopaminergic neurons and MBINs of unknown neurotransmitter), as they displayed different distributions of recurrence. (**H**) Characterization of cells receiving feedback or parallel efference copy signal from DNs^VNC^ or DNs^SEZ^ in 1- or 2-hops. Feedback was defined as a backward connection from a specific output neuron onto a neuron in its own feedforward pathway. Parallel efference copy signals output onto neurons in pathways unrelated to the source output neuron. All connectivity reported here is a-d. (**I**) Schematic of the *Drosophila* larva (i) and how this topology corresponds to different body segments (ii), involved in a diverse set of behaviors (iii). Because the axons of brain output DNs^VNC^ were reconstructed, axonal outputs to different VNC segments could be quantified (Fig. S22). (**J**) Each row represents an individual DN^VNC^ pair with its associated upstream and downstream a-d connectivity in the brain and its projections to the rest of the CNS. Upstream and downstream partner plots (i, iii) depict the fraction of cell types 1-hop and 2-hops from each DN^VNC^ (color legend, *bottom*). The projectome plot (ii) reports the number of DN^VNC^ presynaptic sites in each CNS region. Candidate behaviors are suggested based on known behaviors described in (H, iii). **one DN^VNC^ pair has no strong 2nd-order partners in the brain. (**K**) Schematic of common feedforward and efference copy a-d pathways observed in the brain with a focus on DN^VNC^ connectivity.

We wondered to what extent individual neurons provide feedback to their own upstream partners, thereby forming recurrent loops. We therefore used multihop signal cascades from individual neurons to identify their direct and indirect downstream partners throughout the brain (up to 5 hops). We then determined which of these downstream partners sent recurrent signal back to the source neuron. When analyzing the whole brain in this way, we found 41% of brain neurons were recurrent, i.e. sent signal back to at least one of their upstream partners (Fig. 6E). Furthermore, downstream neurons often sent recurrent signal to upstream neurons using paths of multiple different lengths (Fig. 6F). On average, recurrent communication between a single downstream neuron and its upstream partner used polysynaptic paths of multiple different lengths (on average 1.9 ± 0.9 SD). Such a nested architecture with multi-stage recurrent processing has been suggested as a model for the visual cortex (*138*) and may explain how shallow architectures observed in biology (here, up 8-hops from sensory to brain outputs) can compete with the ultra deep networks often used in machine learning (up to a thousand layers (*139*)).

### Input and output neurons of the learning center are among the most recurrent in the brain

We next analyzed which brain cell classes were the most recurrent (Fig. 6G, Fig. S20B). We define recurrence for individual neurons as the fraction of their polysynaptic downstream partners (using cascades of up to 5-hops) that sent signal back to that source neuron (also using 5-hop cascades) with a-d connections. Therefore, neurons with high and low recurrence scores are engaged in many and few recurrent loops, respectively.

We observed that the fraction of recurrent partners varied widely between distinct neuron classes (Fig. 6G). PNs and the intrinsic neurons of the learning center (KCs) had virtually no recurrent partners (on average, 1.2% and 0.1%, respectively). Interestingly, other neurons associated with the learning center were amongst the most recurrent in the brain: DANs (57%), the modulatory neurons that drive learning; MB-FBNs (51%), presynaptic to DANs and implicated in computing predicted value and regulating learning (*31, 34*); MBONs (45%), the outputs of the learning center and presynaptic to MB-FBNs; and CNs (42%), presynaptic to both MBONs and LHNs, which integrate learnt and innate signals (*92*) (Fig. 6G, Fig. S20B). Together, these four sets of neurons implicated in learning (*25, 31*) and in memory-based action-selection (*34*) form a set of interconnected recurrent loops (Fig. 6G, Fig. S20C). It will be interesting to determine what role this extensive recurrent architecture plays in distinct types of learning tasks.

### Descending neurons provide efference copy to learning center dopaminergic neurons

Many deep brain clusters far from the sensory periphery (Fig. 6C), including many DNs, provided backward signal to many brain neurons. We found that the axons of some DNs^VNC^ (37%) and most DNs^SEZ^ (66%) synapsed onto other brain neurons before descending to the VNC and SEZ, thus providing putative efference copy signals (i.e. copies of motor commands (*140, 141*)). Single DNs broadcasted signal to neurons that were directly or indirectly upstream of themselves (feedback signal) or onto parallel pathways, namely neurons upstream of other output neurons (parallel efference copy signal; Fig. 6H). DNs synapsed onto many different brain neurons (Fig. 6H), including 130 postsynaptic partners and 588 partners 2-hops downstream of DNs^VNC^ and 320 postsynaptic partners and 1284 partners 2-hops downstream of DNs^SEZ^. Of those DNs that synapsed onto brain neurons, we found that individual DNs^VNC^ synapsed on average onto 6 postsynaptic neurons and indirectly (via 2-hops) onto 43 neurons. Individual DNs^SEZ^ synapsed on average onto 8 neurons directly and onto 79 neurons in 2 hops.

We investigated the cell type identities of brain neurons receiving DN^SEZ^ and DN^VNC^ input (Fig. 6H). The most prominent DN^SEZ^ targets werePNs (including direct connections to an olfactory uniglomular PN [uPN 67b], 5 pairs of multi-glomerular PNs, 24 pairs of gustatory PNs) and pre-DN^VNC^ neurons. This suggests that DNs^SEZ^ might modulate upstream sensory processing, as well as locomotor commands that are sent to the VNC. The most prominent DNs^VNC^ targets were pre-DN^VNC^ neurons and MB-related neurons that are thought to play a role in memory-based action-selection (CNs (*34*)) and in driving learning: MBINs (mostly dopaminergic, DANs) and FBNs that integrate MBON input and feed it back onto the MBINs (*31*) (Fig. 6H). DNs^VNC^ also synapsed onto a few PNs (2 nociceptive and 2 gut/mechanosensory PN pairs) and 4 pairs of MB-FFNs (which carry sensory signal to DANs and OANs) (Fig. 6H), suggesting that ongoing locomotor commands could potentially modulate early stages of sensory processing as has been previously described (*140, 142*).

Signal cascades revealed that all DANs and most of their upstream MB-FBNs (90%) receive feedback signal from DNs^VNC^ (Fig. S21A-D), forming larger recurrent loops. DANs even received direct or 2-hop input from DNs^VNC^. DNs^VNC^ also sent robust feedback to MB-FBNs, that are presynaptic to MBINs/DANs (Fig. S21C). Because DNs^VNC^ are often directly or indirectly downstream of MBONs and thought to control locomotor behavioral output (*42, 134, 143, 144*), feedback connections from DNs^VNC^ to the MB may be important for evaluation of behavioral responses and comparing them with actual outcomes to compute error signals that could drive learning (*145, 146*).

### Brain - nerve cord projectome provides a basis to study how the brain controls actions

Our EM volume contains the complete CNS (brain, SEZ and nerve cord), allowing us to assess communication between the brain and the rest of the CNS. Because the majority of motor neurons (MNs) are located in the VNC, understanding brain-nerve cord communication is essential to understanding how behavior is generated. Towards this goal, we reconstructed axons of brain DNs that send feedforward signal outside of the brain. We divided the CNS into 13 regions based on stereotyped landmarks(*147*), including all VNC segments, and determined how many DN presynaptic sites were located in each CNS region (Fig. 6I-i, Fig. S22). This resulted in a brain-VNC projectome directly linked to the connectome. Each VNC segment contains MNs, which innervate muscles in stereotyped positions throughout the body (Fig. 6I-ii). Previous studies have identified body segments involved in specific behaviors (Fig. 6I-iii), such as forward and backward locomotion (*20, 127, 148, 149*), turning (*127–129*), hunch (*29, 150*), speed modulation (*151*), and head movement (*152, 153*). Using this information and the projectome, one can generate hypotheses about which DN^VNC^ might control which behavior.

Using this linked projectome-connectome data, we generated an overview plot that displays, for each DN^VNC^, i) its upstream partners; ii) the location of its outputs throughout the CNS, and iii) all its downstream partners in the brain (Fig. 6J). We annotated the projectome plot with candidate behaviors that each DN^VNC^ might produce (Fig. 6J-ii). Using this data, we grouped DNs^VNC^ based on proposed roles in behavior and examined the pre- and postsynaptic partners of each group (Fig. S23B-C). We found that DNs^VNC^ that may play a role in aversive behaviors (turn, backup, and hunch/head movement) received input from a higher fraction of innate center neurons (LHNs). Meanwhile, DNs^VNC^ that may play a role in appetitive behavior (forward crawl) received input from a higher fraction of learning center output neurons (MBONs). Some members of both types of DN^VNC^ sent feedback to the MB, with direct connections to MB-FFNs and DANs/MBINs (Fig. S23C). These findings suggest that direct paths from the innate center may be more important for aversive behavior, which could require a faster response time (*107, 154*). It should be noted that despite these differences, the primary pre- and postsynaptic partners for all DNs^VNC^ were pre-DN^VNC^ neurons (Fig. S23A), regardless of proposed role in behavior.

As mentioned earlier, multiple feedforward pathways of different kinds and different lengths converged onto DNs^VNC^ (Fig. 6K). There were many short paths via PNs directly onto DNs^VNC^, longer paths through the LH, and even longer ones through the MB. Specifically, 19% and 65% of DNs^VNC^ received direct one-hop input from PNs, respectively. 11% and 66% received both direct and one-hop input, respectively, from both PNs and LHNs. A few DNs^VNC^ received direct or one-hop input only from innate pathways (14%) or only from learning pathways (3%). However, the majority of DNs^VNC^ (80%) received direct or one-hop input both from neurons that encode innate (PNs and LHNs) and learnt valences (MBONs, CNs, MB-FBNs). This suggests that learnt and innate pathways converge at multiple levels in the brain: at the CNs that are directly downstream of LH and MB (*92, 155, 156*), and again at the DNs^VNC^. Such a multilevel convergence architecture that allows representations of distinct valences to be mixed at multiple levels may generate high-dimensional neural representations that enable more complex input-output relationships and offer better discrimination (*38, 157*).

### Most descending neurons target pre- and pre-premotor neurons in the nerve cord

The brain projectome reveals which segments DNs^VNC^ project to, but not the way in which the brain communicates with the VNC circuitry in each segment. To address this question, we analyzed how the brain communicates with the most completely reconstructed VNC segment (A1), in which all motor (*44*) and many sensory circuits (*29, 38*) have been reconstructed. We identified A1 ascending neurons to the brain (Fig. S2) and therefore have all links from the brain to the A1 (DNs^VNC^) and from A1 to the brain (ANs^A1^, Fig. 7A).

**Fig. 7:**
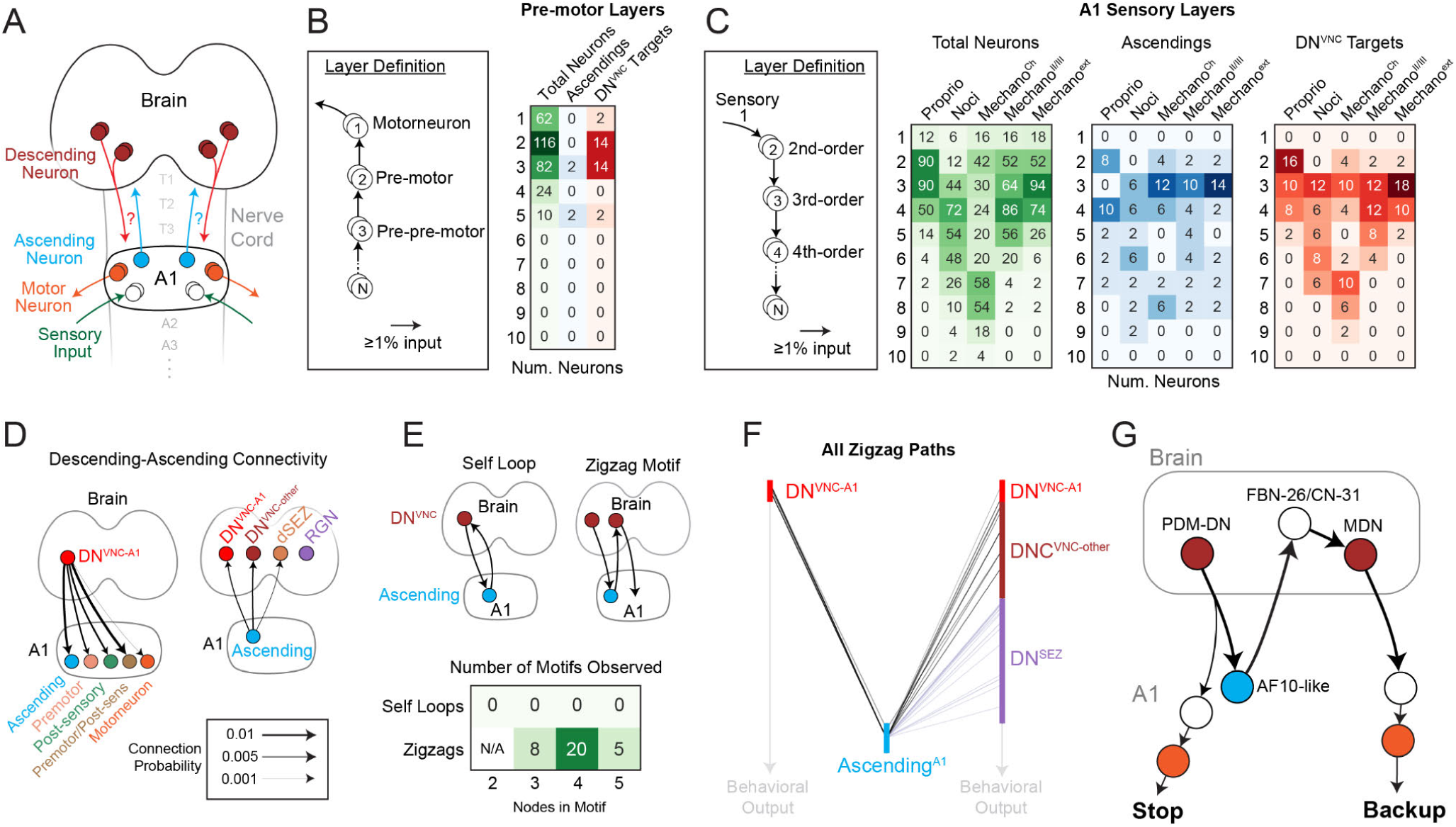
Investigation of brain-nerve cord interactions revealed direct connectivity between ascending and descending neurons. (**A**) Schematic of avenues of interaction between the brain and VNC, namely DNs^VNC^ (red edges) and ANs (blue edges). Analysis focuses on the A1 segment, which is currently the most comprehensively reconstructed VNC segment. (**B**) Pre-motor neuron layers in A1. Layers are identified based on a pairwise 1% a-d input threshold (*left*). Number of interneurons and ANs in each layer are reported (*right*). DN^VNC^ targets refer to neurons in A1 that are postsynaptic to a DN^VNC^. (**C**) Schematic of sensory layers in A1 (*left*). Total number of interneurons (green) and ANs (blue) are reported for each sensory layer and location of DN^VNC^ targets (red). (**D**) Connection probability (a-d) between DNs^VNC^ and A1 cell types (*left*) and between ANs^A1^ and brain output neurons (*right*). ANs did not synapse onto RGNs. (**E**) Quantification of a-d motifs involving DNs^VNC^ and ANs in A1. DN^VNC^-AN reciprocal loops were not observed, while DN^VNC^-AN-DN^VNC^ zigzag motifs were observed. Simplest version of each motif is depicted as a schematic (*top*), but motifs involving 3, 4, and 5 nodes were also assayed. These motifs were allowed to contain additional A1 interneurons or pre-output neurons in the brain. (**F**) Visualization of all zigzag motifs observed. Each bar represents the number of neurons in each cell type and lines represent paths originating and ending at individual cells in each category. (**G**) A zigzag motif with previously characterized DNs^VNC^ on either side. This motif starts at PDM-DN, whose acute stimulation drives a stopping behavior, and ends at MDN, whose acute stimulation causes animals to back up. Stop-backup is a common behavioral sequence observed in the *Drosophila* larva.

First, we characterized the motor and sensory layering in the A1 segment to determine where DNs^VNC^ input onto this structure (Fig. 7A-C). To do this, we quantified how many hops upstream of MNs (for motor layering, Fig. 7B) or downstream of SNs (for sensory layering) each A1 interneuron was (Fig. 7C). 232/342 A1 interneurons (68%) had direct or indirect connections to MNs, whereas 110 (32%) did not. Of those that did, most (198 neurons, 85%) were either directly or 2-hops upstream of MNs, indicating A1 motor circuits are relatively shallow (Fig. 7B). Premotor and pre-premotor neurons were the most prominent DN^VNC-A1^ targets (Fig. 7B). Out of the 42 DNs^VNC^ inputting to A1 (DNs^VNC-A1^), 30 (71%) synapsed onto pre- or pre-premotor neurons (Fig. 7B). 2 DNs^VNC-A1^ (1 pair, 5%) synapsed onto an MN, while 10 DNs^VNC-A1^ (24%) synapsed onto sensory, rather than motor circuit neurons (directly or indirectly downstream of A1 SNs, Fig. 7C).

Interestingly, we found that individual DNs^VNC^ synapsed onto relatively few A1 interneurons, with 1.9 (± 1.4 SD) neurons downstream of each DN^VNC^ and only 48/342 A1 neurons (only 14%) downstream of all DNs^VNC^. Similarly, only a small fraction of premotor (14 neurons, 12%) and their upstream pre-premotor neurons (14 neurons, 17%) were direct targets of DNs^VNC^. Many (71%) of these pre- and pre-premotor DN^VNC^ targets also received direct or indirect A1 sensory input, sometimes from multiple modalities. Finally, we also identified hub neurons in A1 (with ≥10 up- or downstream partners based on A1 network mean plus 1.5 SD) and asked whether DN^VNC^ targeted these hubs. Indeed, DN^VNC^ targeted two hubs, namely A03o (in-hub) and A18b (out-hub).

Premotor neurons have been shown to act combinatorially and flexibly in multiple behaviors (*44*). We found that DNs^VNC^ targeted a small fraction of pre- and pre-premotor neurons, including a couple of hubs, suggesting these DNs^VNC^ targets could play a key role in pushing the combinatorial dynamics of premotor circuits towards particular behaviors.

### Some descending neurons target sensory circuits in the nerve cord

We found that the depth of sensory circuits was varied, with a depth of 3-hops (proprioceptive) to 7/8-hops (nociceptive and chordotonals) from SNs within A1 (Fig. 7C). DNs^VNC^ mostly targeted 3rd or 4th-order SNs (2 or 3-hops downstream of SNs), many of which were also pre- or pre-premotor neurons (31% and 39%, respectively). A notable exception were the proprioceptive circuits. DNs directly synapsed onto several 2nd-order proprioceptive neurons (Fig. 7C). 50% of these were also pre- or pre-premotor neurons. This result is in line with the importance of proprioception in motor control (*158, 159*) and suggests that some behavioral and proprioceptive signals converge immediately downstream of proprioceptive SNs.

We categorized DNs^VNC^ into three types based on their direct downstream targets. The first group of DNs^VNC-A1^ (10 neurons, 24%) preferentially targeted motor circuits, i.e. motor, premotor, or pre-premotor neurons that were not part of A1 sensory circuits. These motor-targeting DNs^VNC^ synapsed onto 8 A1 neurons that had axonal outputs restricted to T3-A1 (3 pairs) or T3-A4 (1 pair). The second group of DNs^VNC-A1^ (12 neurons, 29%) preferentially targeted sensory circuits, i.e. 2nd- or 3rd-order SNs that were not part of A1 motor circuits. These sensory-targeting DNs^VNC^ synapsed onto 12 A1 neurons, including ANs (2 pairs), neurons restricted to A1/A2 (2 pairs), and long-range VNC neurons that output to either all thoracic segments (1 pair) or most abdominal segments (A2-A7; 1 pair). These results suggest that DN^VNC^ modulation of post-sensory cells is propagated across the CNS, including back to the brain via ANs, within A1 itself, and across nearly all VNC segments (T1-T3, A2-A7). This is in contrast to targets of motor-DNs^VNC^ that have more restricted output to T3-A4. The remaining DNs^VNC-A1^ (20 neurons, 48%) targeted both premotor and 2nd-order SNs (Fig. S24), which displayed both broad and restricted arborization patterns. In summary, our data suggests that some DNs^VNC^ may play a bigger role in direct control of motor output, while others may primarily modulate sensory processing in the nerve cord. Nearly half of DNs^VNC-A1^ likely contribute to both activities.

### Direct descending-ascending connectivity reveals novel brain-nerve cord zigzag motifs

To better understand reciprocal brain-nerve cord communication, we analyzed neurons upstream and downstream of A1 ANs. Strikingly, we observed many instances of direct DN^VNC^→AN and AN→DN^VNC^ and AN→DN^SEZ^ connectivity (but no AN→RGN, Fig. 7D, Fig. S25A). Specifically, 12 DNs^VNC-A1^ (30%) synapsed onto 4 ANs in A1 (11%), while 24 ANs in A1 (57%) synapsed onto 22 DNs^VNC^ (12%) and 12 DNs^SEZ^ (7%) in the brain. To test if AN-DN and DN-AN connections were a general feature present in other segments, we assayed connectivity between DNs^VNC^ and all currently reconstructed ANs from all VNC segments. Strikingly, individual DNs^VNC^ received 3.6% (± 5.2% SD) of their input from ANs (1.3% of which was from A1 ANs and 2.3% from other ANs), while DNs^SEZ^ received 1.3 (± 3.0% SD) input from ANs. It should be noted that this is an underestimate because most ANs have not yet been reconstructed. Conversely, individual ANs across the VNC received 4.1 (± 9.7% SD) input from DNs^VNC^. Overall, these results suggest that DN^VNC^-AN interactions may be a general feature of the CNS.

Interestingly, reciprocal loops between DNs^VNC^ and ANs were never observed. Instead, we found zigzag motifs, *DN^VNC^_1_ → AN → DN^VNC^*, with different DNs^VNC^ on each side (Fig. 7E, F). Similar zigzag motifs were also observed involving DNs^SEZ^ (Fig. S25B, C). We hypothesized that these motifs could encode behavior sequences if the DNs promote specific actions and if the ANs provide some sort of feedback about the action that has just been completed. For example, given the motif *DN^VNC^ _1_→ AN → DN^VNC^ _2_*, DN^VNC^_1_ may generate one behavior, which activates the AN both by the motor command and by proprioceptive feedback, which in turn activates DN^VNC^_2_ to generate a second behavior.

To test this hypothesis, we analyzed the sensory information carried by the two pairs of ANs in A1 that participate in zigzags. We found that one pair was presynaptic to proprioceptive SNs, while the other was highly multimodal and 2-hops downstream of most SNs (Fig. S24, see asterisks), including chordotonals, which also play a role in proprioception (*160*). Next, we checked whether DN^VNC^ _1_ and DN^VNC^_2_ in any of the zigzag motifs drive behaviors in known behavioral sequences. Unfortunately, we only know the behavioral roles of a small fraction of DNs^VNC^ (because the driver lines for most have not yet been generated). However, we did find one motif with known behavioral roles for both DNs (Fig. 7G). This motif contained PDM-DN (DN^VNC^ _1_) and the MDNs (DN^VNC^_2_), which promote stop (*43*) and backup (*42*), respectively. Stop-backup is, indeed, a common behavioral sequence in the larva (*30*). Future studies will be required to test this hypothesis further and determine the roles of the newly discovered brain-nerve cord DN_1_-AN-DN_2_ zigzag motifs.

## Discussion

We present a synaptic-resolution connectivity map of the entire brain of the *Drosophila* larva and a detailed analysis of brain structure, including connection types, neuron types, hubs, circuit motifs and brain-nerve cord interactions. Each neuron was split into two compartments, axon and dendrite, resulting in a rich multiplexed network with four connection types, facilitating hierarchical clustering and investigation of signal transduction. To characterize long-range brainwide anatomical pathways, we developed an algorithm that utilizes synapse numbers between neurons to generate and track probabilistic signal propagation across multihop connections. Using this tool, we analyzed feedforward (from sensory to output) and feedback pathways and cross-hemisphere interactions. We found extensive multilevel multisensory integration and extensive interhemispheric communication. The brain had a highly recurrent architecture with 41% of neurons participating in recurrent loops. We found that brain output neurons broadcast their signal (presumably about motor commands) to a wide variety of upstream neurons, including those very close to sensory periphery (2nd-order PNs) as well as DANs that drive learning. Interestingly, the input and output neurons of the learning center were amongst the most recurrent in the brain. Below, we discuss the potential significance of identified architectural features, comparisons with other organisms, and parallels with machine learning architectures.

### Different types of neuronal connections in the brain

We found that the connectome comprised four connection types: axo-dendritic (66.4%), axo-axonic (26.4%), dendro-dendritic (5.4%), and dendro-axonic (1.8%). Axo-dendritic synapses made up the majority of feedforward and feedback connectivity. Axo-axonic synapses tended to form reciprocal connections between neurons, perhaps indicating reciprocal inhibition or gating of axonal output (*84*). Axo-axonic connectivity has also been observed in the adult fly olfactory system (*73*), which may be important for divisive normalization (*85*). While dendro-dendritic and dendro-axonic connections have been previously observed in mammals(*74, 77, 161*), their prevalence in the brain has never been investigated and their functional role is poorly understood. We found that a majority of dendro-axonic connections were the inverse of an axo-dendritic connection, suggesting they may provide instantaneous feedback onto the upstream axon for input regulation (*69*). We also observed multiplexed connections between neurons, including up to all four types simultaneously. The most common multiplexed connection, axo-dendritic with axo-axonic, may grant the presynaptic neuron post- and presynaptic control of the downstream neuron, as has been observed in triad motifs in mammals(*78*). Overall, our comprehensive analysis revealed non-canonical connectivity throughout the brain. Future experimental studies are needed to elucidate their roles in neural computation.

### Connectivity-based clustering reveals 90 distinct types of brain neurons

Neuron types in various organisms have been classified based on their function (*28, 31, 90, 91, 162*), morphology (*10, 89*), gene expression (*86–88*), or combinations of features (*10, 13, 93*), such as morphology and connectivity. While it is reasonable to expect that all these features are correlated, it is still unclear which one is ideal for defining neuron types and how neuron types defined based on different features correspond to each other. Here, we used synaptic connectivity alone to perform an unbiased hierarchical clustering of all neurons and identified 90 types. We found that the morphology of neurons within clusters was remarkably similar, even though clustering was based solely on connectivity and no morphological data was used. Furthermore, neurons that had similar known functions in behavior were usually found in the same or related clusters. Thus, clustering neurons based on synaptic connectivity resulted in clusters that were internally consistent for other features, when those features were known. However, many of the identified clusters contained previously uncharacterised neurons with unknown physiology and function. The comprehensive atlas of neuron types and individual neurons generated by our reconstruction can be used in the future to guide the development of tools for their selective targeting and characterizing their functional properties and patterns of gene expression.

### Pathways from sensory to output neurons form a multilayered distributed network

Sensory information is thought to be processed both serially, through multiple successive processing steps that extract progressively more complex features (*163, 164*), and in parallel, with different features processed simultaneously in parallel pathways (*165, 166*). However, the exact architecture of sensory circuits is not fully understood: how many processing steps from sensory inputs to brain outputs; how many parallel pathways for each sensory modality; how much cross-communication between distinct pathways of a single modality and between distinct modalities? The comprehensive nature of our connectome enabled us to address these questions by systematically analyzing all multihop pathways from all sensory modalities to brain output neurons. We observed multiple parallel pathways of varying depths downstream of each modality, albeit with extensive interconnectivity between different pathways. This architecture suggests that distinct features may not be processed independently, but that each feature may potentially influence the computation of most other features in a distributed network.

We found that the shortest paths from sensory neurons to output neurons are surprisingly shallow. All output neurons receive input from sensory neurons within a maximum of 3 hops. However, most output neurons also received input from the same modality via multiple longer pathways. Thus, output neurons receive both relatively direct, as well as highly processed sensory information, transformed at multiple prior steps. Convergence of shorter and longer pathways onto command-like neurons has been previously observed in the nerve cord (*38, 44*). The shorter pathways could potentially enable quicker responses when sensory input is strong and unambiguous and may be evolutionarily older than the longer pathways that enable more complex action-selection based on multisensory context and learning. Consistent with this, in zebrafish, circuits regulating increasingly complex behaviors are layered on top of pre-existing circuits for more basic responses that are built early in development (*167*).

Architectures with longer and shorter paths that skip layers have also been utilized in prominent machine learning networks, including deep residual learning in ResNets (*50*) and DenseNets (*168*) and sequential shortcuts in U-Net convolutional architectures (*169*). While the predictive accuracy of artificial neural nets can be improved by simply increasing network depth (*170*), accuracy can saturate and degrade if too many layers are added. Increasingly complex and abstract features are integrated with each additional layer (*171*), but if these features become too abstract, learning degrades. Shortcuts between layers can solve this problem (*50, 172*) by combining lower level features as an additional teaching signal in addition to the more abstract high level features. Adding shortcuts to shallow networks increases their computation capacity, allowing them to compete with or outperform deep networks lacking shortcuts. Taken together, these lessons from ANNs suggest that, perhaps, the layer skipping we observed allows the brain network to increase computational capacity given a finite depth/number of neurons constrained by evolution and development.

### The majority of brain neurons integrate input from all sensory modalities

Integrating information from multiple modalities improves signal-to-noise and reduces ambiguity. Thus, many higher-order neurons involved in learning and action selection are known to be multisensory (*173, 174*). Consistent with this, here we found that 80% of brain output neurons that control locomotion (DN^VNC^) receive input from all sensory modalities and only 2% are unimodal.

Some studies have also shown that multimodal integration can start early in sensory processing and occur at multiple levels (*38, 63, 175*). However, whether early and multilevel multimodal integration are a general principle was unclear. Here we found that the vast majority (88%) of brain neurons are multimodal and a striking 62% integrate input from all modalities. Multisensory integration was pervasive not only in known action-selection neurons and output neurons (e.g neurons downstream of MB, pre-DN^VNC^ and DN^VNC^), but also in neurons close to sensory periphery. Most unimodal neurons were 2nd order PNs postsynaptic of sensory neurons. However, even amongst PNs, only a minority (35%) were unimodal. Thus, brain feedforward circuits that link sensory inputs to brain outputs form a large distributed network with extensive convergence between modalities at multiple levels. Such multilevel convergence architecture may enable more complex input-output relationships offering better discrimination between multisensory events (*157, 176*).

### Recurrent architecture of the brain with multiple nested loops

Feedback has been observed in many brain circuits and implicated in a range of computations, including normalization, object detection, comparing expected and actual outcomes, working memory and others (*135, 177–179*). However, the exact architecture of feedback pathways and the amount and nature of feedback that each neuron receives is still poorly understood. Here, we used signal cascades to systematically identify all connected pairs of brain neurons (with up to 5-hops) that had a reciprocal connection (of up to 5 hops). We found that a striking 41% of brain neurons received recurrent input from at least one of their downstream partners. Furthermore, most pairs of connected neurons were connected via multiple recurrent pathways of varying lengths, forming multiple nested loops. Recurrent nested structure has been shown to compensate for a lack of network depth in artificial neural networks (*138*) and is thought to allow neural nets to use arbitrary computation depth depending on the task (*51*).

### Learning center dopaminergic neurons are amongst the most recurrent in the brain

DANs were amongst the most recurrent neurons in the brain with almost all DANs reciprocally connected with more than 50% of their partners (via multihop connections). DANs are central for learning, motivation, and action across the animal kingdom (*117–120*) and are implicated in a range of human mental disorders (*180, 181*). The highly recurrent connectivity of DANs could potentially deliver high dimensional feedback (*182*), enabling them to encode a range of distinct features and flexibly engage in many parallel computations. Recurrent loops could also play roles in working memory (*31, 132, 133, 183*).

We provide, for the first time, a comprehensive list of multihop (up to 5 hops) feedback and feedforward inputs onto DANs. Previous studies have reported that DANs receive extensive feedback from neurons downstream of the MB that integrate learnt and innate values (*31*). Here, we find that DANs also receive extensive feedback from brain output neurons, DNs^VNC^ and DNs^SEZ^, which likely encode motor commands for locomotion and feeding. Furthermore, DANs receive extensive polysynaptic feedforward inputs from all sensory modalities. In addition to the previously known inputs from sensory neurons that encode unconditioned stimuli (rewards and punishments), DANs also received polysynaptic input from sensory neurons that normally encode conditioned stimuli in learning tasks, including olfactory (also observed in the adult fly (*100*)), visual, and thermosensory, and from proprioceptive neurons. Recent studies, in the adult fly, have shown that DAN activity correlates with movement (*184*). These movement-related signals could be explained by efference copy input from DNs^VNC^ and/or by input from proprioceptive sensory neurons. Thus, in principle, DANs could compare motor commands with proprioceptive feedback on actual movement.

In summary, the observed architecture gives DANs access to multiple kinds of information that is thought to be critical in driving reinforcement learning: actual rewards (*122*), predicted values (*145*), selected actions (*185*), and proprioceptive feedback about ongoing actions (*184*). Future functional studies will reveal how all of this incoming information is used by the DANs during different kinds of learning tasks.

### The majority of the brain hubs are directly downstream or upstream of the learning center

In C.elegans, in-out hub neurons have been shown to play essential roles in behavior (*186, 187*). Here, we found that the majority (73%) of the larval brain’s in-out hubs were postsynaptic to the learning center output neurons (MBONs) and/or presynaptic to the learning center modulatory neurons (DANs, OANs, MBINs). Many (23%) were also postsynaptic to the center that mediates innate behaviors (the LH), thus integrating both learnt and innate values (*92*). A prior functional study of one of these hubs, the MBON-m1, has shown it computes an overall predicted value of stimuli by comparing excitatory and inhibitory input from neurons that encode positive and negative value, respectively. MBON-m1 bidirectionally promotes actions, promoting approach and avoidance when its activity is increased and decreased, respectively. Several additional hubs identified here (CN-12, CN-13, CN-37) have similar patterns of input to MBON-m1, raising the possibility that they also play a similar role in computing predicted values. These hubs also provide direct feedback to the MB DANs and could therefore play roles in regulating learning. Future studies of the brain in-out hubs could therefore shed important insights into the mechanisms of value-computation, action-selection and learning.

### Cross-hemisphere interactions

Why have two brain hemispheres instead of just one? In primates and humans, this redundancy facilitated evolution of computational lateralization in the cortex (*188, 189*). Brain lateralization can occur in insects (*190, 191*), but it is rare and developmental differences between hemispheres are even compensated for (*192*), suggesting there are other advantages to a bilateral brain. Here, we found that the brain utilized neurons from both hemispheres in a majority of pathways from sensory to output neurons. Contralateral connections were particularly important to transmit sensory signal to descending neurons (DNs^VNC^), suggesting that contralateral neurons may be bottlenecks for propagating information. Consistent with this idea, the majority of brain in-out hubs (88%) had contralateral axons. Previously, it has been postulated that interhemispheric communication increases the computational power of the brain by providing access to additional computational units (neurons) and pathways (*193*). Our current work suggests this hypothesis is plausible and provides the ability to target specific contralateral hubs to test their role in future experimental studies.

We also found a striking propensity for contralateral neurons to connect to each other. Furthermore, we discovered a group of neurons that form reciprocal loops between left-right homologous partners (called pair loops). The interpretation of pair loops depends on their sign. If inhibitory, they may be important for interhemispheric comparisons. If excitatory, they may be involved in signal perpetuation/short-term memory (*130, 132*). Indeed, study of split-brain patients with commissurotomy demonstrated that interhemispheric communication is important for short-term memory in humans (*131*). Consistent with this idea, many pair loops occurred between neurons implicated in memory formation, the MB-FBNs.

### Brain and nerve cord interactions

To provide a basis for understanding how the brain controls locomotion and regulates somatosensation, we investigated the connectivity patterns between brain output neurons (DNs^VNC^) and the most reconstructed nerve cord (VNC) segment (A1) (*29, 38, 44, 194*). We found that relatively few DNs^VNC^ (42 DNs^VNC^, 23%) directly targeted a small fraction of A1 neurons (48 neurons, 14%). These DNs^VNC^ were categorized into three different groups: 1) those that target pre- and pre-premotor neurons, 2) those that target low-order post-sensory neurons, and 3) those that target both. Premotor-targeting DNs^VNC^ are likely involved in biasing behavioral output, while the post-sensory-targeting group is likely involved in modulating sensory processing. These different DN types targeted A1 neurons with distinct arborization patterns. The post-sensory neurons innervated most VNC segments, while the premotor neuronal arbors were more restricted. Thus, motor-DNs^VNC^ appear to target a few key premotor control elements within each segment to switch between locomotor states. Meanwhile, DNs^VNC^ that modulate sensory processing appear to have global effects across the VNC through widely projecting A1 interneurons.

We also found that DNs directly synapsed onto ascending neurons (ANs) and ANs directly synapsed onto DNs, a general phenomenon observed in multiple segments throughout the VNC. Some of these DN-AN and AN-DN connections formed zigzag motifs (DN_1_→AN→DN_2_), but never reciprocal loops. ANs within zigzag motifs carry proprioceptive information, raising the possibility that such motifs could encode behavioral sequences. The AN in zigzag motifs may signal the completion of a first action promoted by DN_1_ and facilitate the initiation of the next action promoted by DN_2_. Consistent with this idea, recent studies have identified examples of direct connections from ANs to DNs, and these ANs mediate specific actions (*159, 195, 196*). Investigating hypotheses about the roles of the newly identified zig-zag motifs and DN-AN connectivity generally will require a full characterization of the behavioral roles of the DNs and the signs of their connections with ANs.

### Conclusions

In summary, we have provided a synaptic-resolution map of an insect larval brain and developed a methodology for comprehensive analysis of large-scale connectomes. We characterized connection types, neuron types, network hubs and brainwide circuit motifs. The connectome provides a valuable resource and a basis for many future theoretical and experimental studies. The genetic tools available in this tractable model system (*35–37*) will allow selective visualization and manipulation of individual neuron types and testing the functional roles of specific neurons and circuit motifs revealed by the connectome. Furthermore, although the details of brain organization differ across the animal kingdom, many circuit architectures are conserved (*25, 45, 90, 197–200*). We therefore expect that the architectural features described here will be useful in future studies of both invertebrate and vertebrate brains as well as in network science (*201*). We found structural features, including multilayer shortcuts and robust recurrence, that were reminiscent of prominent machine learning architectures. Future analysis of these similarities and differences will be useful to learn more about brain computational principles and perhaps inspire new machine learning architectures. We therefore expect that this connectome will become a continued source of hypotheses and inspiration for future research.

## Acknowledgements

The authors thank HHMI Janelia Research Campus for funding and support, the Janelia Fly EM Project Team for imaging the EM volume, and the Janelia Visiting Scientist Program for outstanding support over the years. We thank Elise C. Croteau-Chonka for providing the base image for larva illustrations in Fig. 6.

The neuronal and synaptic reconstruction presented in this paper was made possible through the collaborative efforts of 86 individuals. We would like to acknowledge the work of all those who contributed to the reconstruction of the larval brain, either directly for this paper or through past publications:

> *Akira Fushiki, Ingrid Andrade, Michael Winding, Avinash Khandelwal, Javier Valdes Aleman, Feng Li, Philipp Schlegel, Albert Cardona, Ivan Larderet, Kathi Eichler, Volker Hartenstein, Anton Miroschnikow, Timo Saumweber, Jennifer Lovick, Matthew Berck, Casey Schneider-Mizell, Nadia Riebli, Larisa Maier, Alex S. Bates, Laura Herren, Ilona Brueckmann, Bruno Afonso, Andreas S. Thum, Lindsey Claus, Haluk Lacin, Alex MacLachlan, Sebastian Hückesfeld, Andreas Schoofs, Simon Sprecher, Aref Arzan Zarin, Hanbo Chen, Elizabeth Barsotti, Ana Correia, Julie Tran, Chris Q. Doe, Nadine Randel, Suguru Takagi, Liria Masuda-Nakagawa, Chris Barnes, Maarten Zwart, Nusreen Imambocus, Eri Hasegawa, Chris Wreden, Andrey Stoychev, Xinyu Tang, Aravi Samuel, Anita Burgos, Julius Jonaitis, Keira Turner, Atsuki Hiramoto, Tomoko Ohyama, Scott Wilson, Sam Qian, Jim Truman, Hiroshi Kohsaka, Julia Meng, Brittany Kemp, Qianhui Zhao, Ellie Heckscher, Waleed Osman, Davi Bock, Jamie Macleod, Andrew Champion, Tihana Jovanic, Rebecca Arruda, Eisuke Imura, Brandon Mark, Mason Klein, Laura Lungu, Marc Corrales, Claire Julliot de La Morandière, Tara Guillorit, Matthieu Louis, Stephan Gerhard, Atit Patel, Li Guo, Yinhui He, Katerina Karkali, Joao Picao Osorio, Maxime Lehman, Tom Kazimiers, Thuc To, Akinao Nose, Edward O’Garro-Priddie, André Ferreira Castro, Basel El Galfi*

## Funding

Howard Hughes Medical Institute Janelia Research Campus (MZ, AC, MW, AF, NR, IVA)

Howard Hughes Medical Institute Visiting Scientist Program (AV)

Wellcome Trust grant 205038/Z/16/Z (AC)

Wellcome Trust grant 205050/Z/16/Z (MZ)

ERC grant ERC-2018-COG: 819650 (MZ)

MRC LMB Core Funding (AC, MZ)

NSF GRFP grant DGE1746891 (BDP)

NSF CAREER Award 1942963 (JTV)

NIH BRAIN Initiative RF1MH123233 (JTV, CEP)

DARPA D3M contract FA8750-17-2-0112 (HGP, YP, CEP)

Air Force Research Laboratory contract FA8750-18-2-0035 (HGP, YP, CEP)

DARPA contract FA8750-20-2-1001 (HGP, YP, CEP)

NIH Grant 2 R01 NS054814 (IVA, VH)

The U.S. Government is authorized to reproduce and distribute reprints for Governmental purposes notwithstanding any copyright notation thereon. The views and conclusions contained herein are those of the authors and should not be interpreted as necessarily representing the official policies or endorsements, either expressed or implied, of the Air Force Research Laboratory and DARPA or the U.S. Government.

## Author Contributions

Conceptualization: MW, BDP, CEP, JTV, MZ, AC

Neuron Reconstruction: MW, AF, IVA, AK, JVA, FL, NR, EB, AC

Methodology: MW, BDP, CLB, HGP, YP, TK, RF, VH, CEP, JTV, AC

Analysis: MW, BDP, CLB

Supervision: MW, CEP, JTV, MZ, AC

Writing: MW, BDP, MZ, AC

Writing - Review & editing: MW, BDP, RF, VH, CEP, JTV, MZ, AC

## Materials and Methods

### Electron Microscopy Data and Reconstruction

The EM volume of the central nervous system (CNS) of the 6 hour old *Drosophila melanogaster* first instar larva used in this study has been previously reported (*38, 40*). Briefly, the genotype of this animal was *Canton S G1 [iso] x w1118 [iso] 5905*. The resulting EM volume contains 4841 z-slices with a x,y,z resolution of 3.8 x 3.8 x 50 nm. This dataset includes the complete CNS, including all neurons, synapses, and accessory structures. Note that only the axons and dendrites of sensory neurons and motor neurons, respectively, are present in the volume. However, the morphology and location of these neurons was sufficient to match them to the respective neurons in whole animal datasets and thereby identify the identities and modalities of sensory axons (*45–47, 49*) or the corresponding muscle targets of motor neurons (*44*).

We identified the boundaries of the brain hemispheres and all brain neurons within using stereotyped landmarks (*202*). Neurons and synapses were manually reconstructed by multiple users using the Collaborative Annotation Tool for Massive Amounts of Imaging Data, CATMAID (*39*). Many previous publications have contributed to the reconstruction of neurons in this CNS (*25, 29, 38, 44–47, 49, 194*), so the completeness of brain neurons was first assessed using review and publication status. A complete census of the brain was conducted by examining each lineage entry point (*202*) to identify all brain cell bodies. Each cell body was then used as a seed point for iterative reconstruction by multiple users until all arbor end-points were identified. The reconstruction process generally followed previous descriptions (*38, 40*), however a targeted review process was used by comparing left-right homologous neuron pairs. Quantification of the results of this methodology suggests it produced neuron reconstructions that are robust across multiple metrics (Fig. S1E, F), although some errors of omission were observed.

### Clustering

We developed a modified spectral clustering procedure to cluster brain neurons based on connectivity. To achieve a clustering in which homologous left/right neuron pairs are likely to be in the same cluster (as opposed to having clusters comprised of left-only or right-only neurons), we developed a technique to perform a spectral embedding which collapses left/right symmetry into a single embedding space. First, the network was split into four subgraphs: connections from neurons on the left side to neurons on the left side (LL), from right to right (RR), from left to right (LR), and from right to left (RL). Each subgraph had its edge weights transformed using a procedure called pass-to-ranks, a regularization scheme which replaces each edge weight with its normalized rank among all edges and is helpful for spectral embedding in the context of outliers or skewed edge weight distributions(*203–205*). We then embed each subgraph into a *d*-dimensional Euclidean space (*d* = 24) using the adjacency spectral embedding (ASE) as implemented in graspologic (*204, 205*). Due to an orthogonal nonidentifiability associated with the latent position estimates from ASE (*204*), we used a joint optimal transport/orthogonal Procrustes procedure (*206, 207*) to align the latent positions of the LL and RR subgraphs, and separately the LR and RL subgraphs. This procedure yields a representation for each node in terms of its ipsilateral (LL or RR) inputs and outputs, as well as its contralateral (LR or RL) inputs and outputs. To achieve a single representation for each node which is amenable to clustering, we concatenated each of these representations per node, and performed another singular value decomposition to further project each node into a lower-dimensional space (*d* = 12). This procedure is inspired by the Multiple ASE (MASE) procedure (*208*). Finally, to ensure that homologous neuron pairs are clustered the same way, we average the embeddings for a left and right node (note that most of these points were already close in this embedded space due to the procedure described above).

With this representation for each neuron, we clustered using a hierarchical approach to Gaussian mixture models (GMM) inspired by past work on hierarchical stochastic block models (*209, 210*). GMM on an ASE embedding was recently shown to be a consistent way of estimating the membership assignments for a statistical network model called the stochastic block model, motivating this approach (*204, 211*). We utilize a Python implementation of GMM with model selection (*212, 213*). In the hierarchical paradigm, all neurons currently under consideration are clustered using a 1-component and 2-component GMM. The fit of both models is evaluated using the Bayesian information criterion (BIC) metric (*214*), which is commonly used to select the number of clusters in a GMM (*215, 216*). If the 2-component model is preferred by the BIC score and the number of neurons is not too small (32 neurons is chosen as the cutoff), then the set of neurons under consideration is split according to this clustering. This procedure was allowed to recurse until the depth of the corresponding “cluster tree” reached eight, yielding a multiresolution clustering of the brain connectivity.

### Walk-Sort

We developed a method based on random walks to order each neuron in the brain based on proximity from user-defined input neurons to user-defined output neurons. First, we selected input neurons (known SNs or ANs from the VNC) or output neurons (neurons projecting to the VNC, SEZ, ring gland and motor neurons) of the brain. We then considered these groups of neurons in (input, output) pairs, and started 256 random walks from each neuron in the input category. The probability of walking from node *i* to node *j* was defined as the number of synapses from node *i* to node *j* divided by the number of output synapses from node *i*. We kept any random walks which went from the given input to output neuron groups in fewer than 16 hops. This process was repeated for each pair of (input, output) neuron groups, as well as for random walks run in reverse (traversing edges opposite to their true orientation). After generating these paths, each neuron in each path was given a rank based its order in the path divided by the path’s number of hops:

Example walk: [(sensory) A, B, C, D, E (output)]

Walk orders: [0, 1, 2, 3, 4]

Normalized walk orders: [A: 0, B: 0.25, C: 0.5, D: 0.75, E:1]

Each neuron was then given a “walk sort” score, defined as its median walk order across all of the walks in which it was visited.

### Pairs via Graph Matching

We employed a family of techniques based on the Fast Approximate Quadratic (FAQ) graph matching algorithm (*58–60, 217*) to predict bilateral neuron pairs on the basis of connectivity. These algorithms seek to find a 1-to-1 alignment of one network’s adjacency matrix with respect to another which minimizes the norm of their difference. In this case, the two adjacency matrices were the induced subgraphs (all connections among a specified subset of nodes) of the left and right hemispheres (i.e. the ipsilateral connections) of the brain. We used 406 ground-truth neuron pairs from previous publications (*25, 31, 45, 162*) as seeds, specifying a fixed, partial alignment between the two networks. The seeded graph matching algorithm (as implemented in iGraphMatch (*218*) or graspologic (*205*)) was randomly initialized 50 times (while preserving the known matching from the ground truth pairs). Predicted pairs from each initialization of the algorithm were recorded. We then ranked potential pairs according to how often they were matched to each other, manually reviewing each potential pair for correctness. This process was iterated multiple times, with newly identified pairs added to the population of seed pairs, until all reasonable pairings were exhausted.

### Pair Ranking by Connectivity Similarity

To quantify the similarity of neuron pairs based on connectivity (Fig. S1E), we used a graph-matching-based metric to avoid biasing our scoring towards the bilateral pairing we sought to evaluate. Let *A* be the adjacency matrix of the left hemisphere induced subgraph, and let *B* be the adjacency matrix of the right hemisphere induced subgraph. For both networks, we use the number of synapses across all edge types as the edge weight. The graph matching problem (GMP) seeks to minimize the objective function

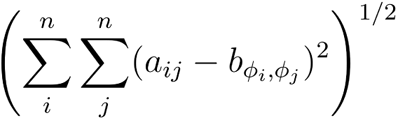

over the set of matchings (or permutations) *ϕ*. A matching *ϕ* is an *n*-length vector, where *n* is the number of nodes, and if *ϕ_i_* = *k*, indicating that node *i* in *A* is matched with node *k* in *B*. In other words, this objective function measures a squared error between the adjacency matrices of the two networks under some 1-to-1 mapping of the nodes from one network to the other. Many modern approaches to solving the GMP relax the permutation constraint to the convex hull of the permutation matrices. Intuitively, this relaxation can be thought of as a *soft* matching between nodes in the two hemispheres. We ran a previously developed FAQ graph matching algorithm (*60*), using a maximum of 30 iterations, 20 initializations, and *α* = 0.1 (see original publications for algorithm details). Note that the annotated pairs were not used as seeds for this analysis and the initializations were random; thus, these annotations did not bias the graph matching. Instead of projecting to the set of permutation matrices at the end of the algorithm, we instead take a weighted sum of the doubly stochastic matrices over each initialization in order to provide a soft matching between potential neuron pairs that can be used for ranking. Let *s_k_* be the objective function value after optimization for initialization *k*, and let

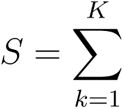

where *K* = 20 is the number of initializations. Let *D_k_* be the final doubly stochastic matrix at the end of optimization for initialization *i*. Then, we use the weighted sum

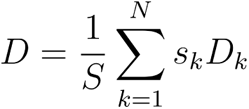

to find a final doubly stochastic matrix, *D*. We then assessed how well bilateral pairs were matched by the soft assignment matrix *D*. To do so, we ranked the elements of each row *i* of *D* (settling ties using the average) and then assessed the rank of the known bilateral neuron pair in this connectivity ranking.

### Multiplexed Edge Analysis

#### Signal flow

To roughly order the network from sensory neurons to output neurons without specifying a start or end of the network (Fig. S6), we applied the “signal flow” algorithm (*82, 83, 219*). This algorithm uses an energy function based on the sum of edge weights which point from a lower-scored neuron (closer to output) to a higher-scored neuron (closer to sensory), and minimizes this function over the set of possible 1-dimensional scores for each neuron. We ranked these signal flow scores for each neuron independently for each network type (axon to dendrite, axon to axon, etc.). For pairwise comparisons of all network orderings, we computed the rank correlation (Spearman’s *ρ*) between the signal flow rankings for each network.

#### Edge reciprocity

Reciprocity is a commonly used metric in network science which quantifies the probability that two nodes in a directed network are connected via mutual edges in each direction (*220*). Specifically, it is defined as the number of reciprocal edges divided by the total number of edges, where a reciprocal edge means that both *A_ij_* and *A_ji_* are present in the adjacency matrix *A*. Here, we generalize this notion to multigraphs. With *A^source^* representing the unweighted, loopless adjacency matrix for the source network, and *A^target^* defined likewise for the target network, we define the edge reciprocity *r*(*A^source^*, *A^target^*) as

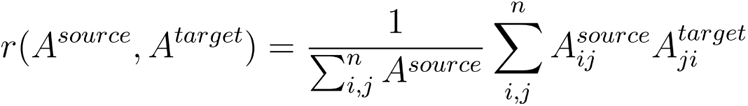

In other words, averaged over the entire network, this is the conditional probability of observing edge 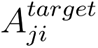 conditioned on observing edge 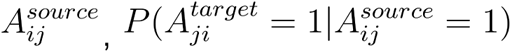.

#### Multiplex edge probabilities

To examine the likelihood of various multiplexed edge combinations, we counted the number of (*i, j*) pairs with each possible combination of edge type occurrences in the measured networks (e.g. an axo-dendritic edge with no other type present, axo-dendritic and axo-axonic but no other edge types, etc.) (Fig. S5D). To calibrate expectations for these counts, we used a simple null model of multiplex edge overlaps. This model assumed that each of the four edge type graphs was generated independently, and modeled each network as a completely random (Erdos-Renyi) network. To compute the parameters of this model, we first simply calculate the global connection probability *p* for each network *A*^(*k*)^ as

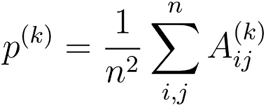

Under the assumptions above, the expected number of (*i, j*) pairs which have only axo-dendritic (*AD*) edges (denote this *m*([1, 0, 0, 0])) is:

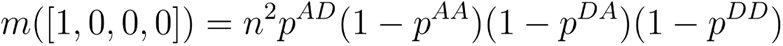

More generally, we denote to be a 4-dimensional binary vector, which indicates the presence (1) or absence (0) of the *AD*, *AA*, *DA* or *DD* edge types, respectively. Then, we can write the expected number of edges under edge type pattern as:

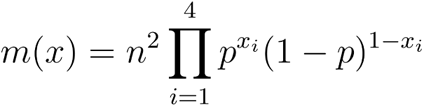

Under this definition, we calculated the expected number of edges for each combination of the four edge types, and used this to compare to the observed counts.

### Signal Cascades

We applied a technique for modeling information propagation through a network based on the independent cascade model, which has been used to study epidemic and social information transmission through networks (*221, 222*). Briefly, the algorithm (which we call the signal cascade) starts with a set of active nodes which are then allowed to propagate their active state to other nodes according to a set of probabilistic rules based on the number of synapses from active to inactive neurons. At each time step, a new set of nodes becomes active, and previously active nodes enter a deactivated state in which they cannot be activated again during that experiment. We modified the original independent cascade model to include a set of “stop” nodes from which the cascade does not proceed further, allowing us to isolate effects from feedback from a specific set of neurons during a given experiment. This tool allows one to determine how much signal from a given set of starting neurons could reach other sets of neurons in the brain, and after how many timesteps (hops). Our approach differs from some previous models of signal propagation across a connectome in that we only allow activation from neurons which were active at the last timestep, rather than from neurons which were activated at any previous timestep (*155, 223*), allowing us to assess the temporal ordering of the potential flow of information through the brain.

To elaborate on the details of the model, the algorithm starts with a set of user-defined nodes which are initially in an active state at time *t* = 0, and all other nodes in an inactive state, meaning they are susceptible to activation. We denote the set of active, inactive, and deactivated nodes at timepoint *t* as 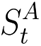, 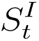, and 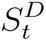, respectively. Our modified cascades algorithm also includes a set of nodes *S^E^* which are “end” nodes from which the cascade no longer continues - these nodes can become active, but then do not propagate their signal at the next timepoint. To determine which nodes become active at the next timepoint *t* + 1, each synapse is assigned an equal probability *p* of transmission, with *p* = 0.05. For each outgoing synapse (*i* → *j*) from each active node which is not a stop node 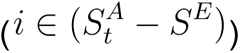 to each previously unactivated node 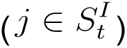, we conduct an independent Bernoulli trial with probability *p* to determine whether that synapse activates node *j* at the next timepoint. Nodes that had at least one successful activation of an upstream presynapse are included in the set 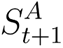. Every node that was active at time *t* is moved to the set 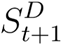, the deactivated nodes which cannot be activated again during the current cascade. This process was repeated for *T* timesteps, where *T* could vary depending on the particular question of interest. These cascades were run 1,000 times for the same set of start and end nodes 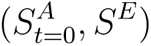. To understand how signals could propagate through the brain based on this model, we tracked the probability that a node was active at a given time over these 1000 independently run cascades. When analyzing groups of neurons, signal cascade data was aggregated by averaging these activation probabilities across neurons in a group.

### Morphological similarity calculation within neuron groups

To quantify the similarity between neuron morphologies within clusters (Fig. 3B; Fig. S7A, B), we applied the NBLAST algorithm (*224*) as implemented in navis (*225*), computing NBLAST scores between all pairs of neurons on the same hemisphere. To make NBLAST scores symmetric (same score between neurons (*i, j*) as between (*j, i*) we set the NBLAST scores for (*i, j*) and (*j, i*) to be the geometric mean of their original scores. We then apply a normalization scheme to each pairwise NBLAST similarity matrix, in which scores are converted to their pairwise ranks in the similarity matrix (*205*). With these normalized NBLAST scores, we define a simple score of morphological similarity within each cluster. First, we computed the mean of all pairwise similarity scores between neurons in a hemisphere of a specific cluster. Then, we took the mean of those average scores between left and right hemispheres to compute the final score for a given cluster.

### Code

Many analyses relied on NumPy (*226*), SciPy (*227*), Pandas (*228*), NetworkX (*229*), navis (*225*), and python-catmaid (pypi.org/project/python-catmaid/). Plotting was performed using matplotlib (*230*), Seaborn (*231*), and Blender (https://www.blender.org/). UpSet plots were used to visualize complex intersections (*232*).

## Supplementary Materials

**Fig. S1.**
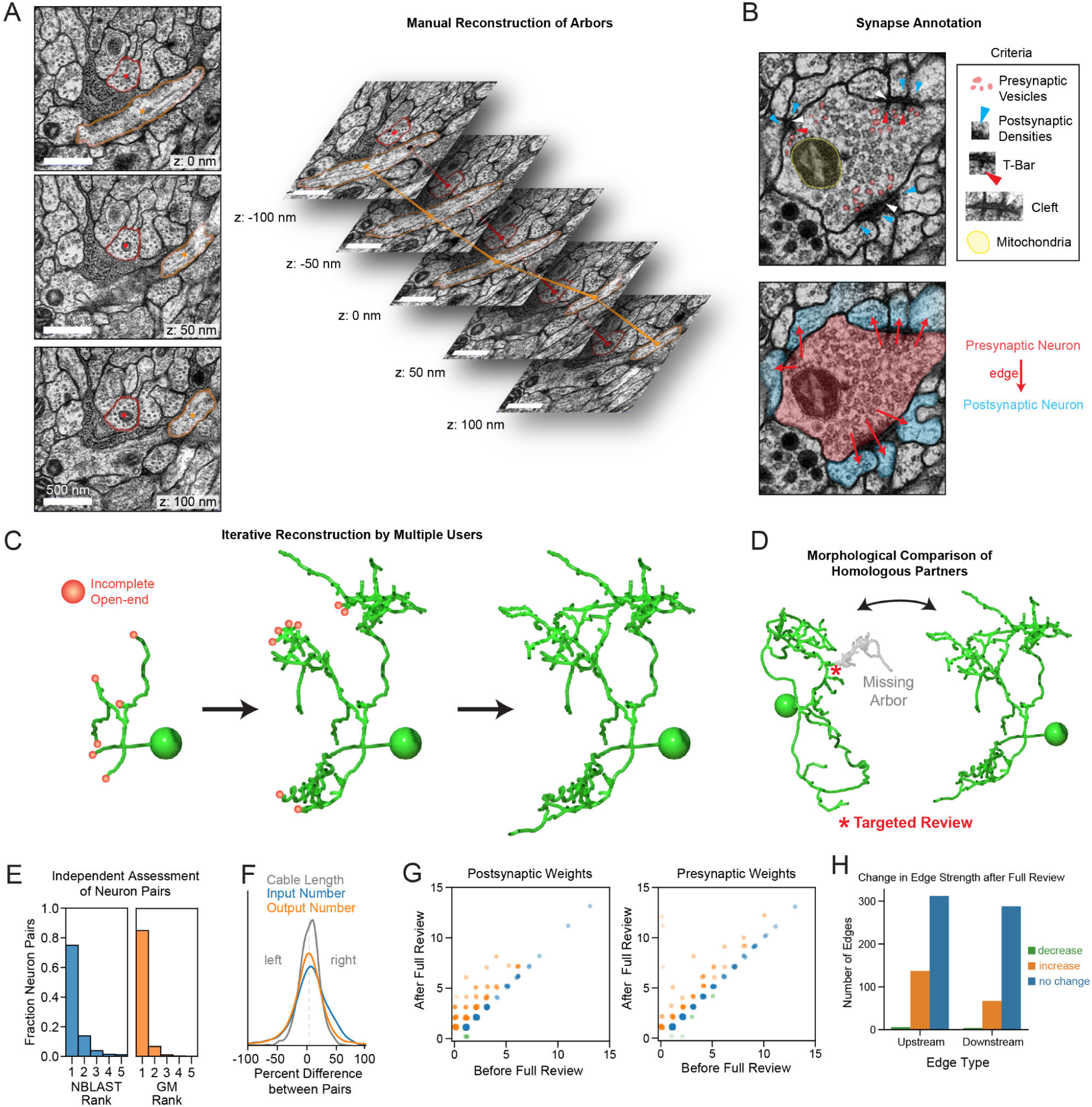
Reconstruction methodology. (**A**) Slices from a whole CNS ssTEM (serial section transmission electron microscopy) volume of the first-instar *Drosophila melanogaster* larva. The membranes of two example neurons are highlighted. To reconstruct neuronal arbors, users annotate the center of each neuron in ssTEM slices using the CATMAID software. These nodes are automatically connected to generate three-dimensional skeleton representations of each neuron. (**B**) Criteria to identify pre- and postsynaptic sites in the ssTEM volume. Synaptic sites are only annotated if they: 1) display presynaptic vesicles, 2) postsynaptic densities in postsynaptic cells, 3) contain a presynaptic T-bar structure that can be observed over multiple z-slices, 4) a synaptic cleft, evident as a subtle black-white-black pattern between the membranes of synaptic partners, and 5) nearby mitochondria. (**C**) Methodology for whole brain reconstruction. All neuron cell bodies were identified in the brain and reconstructed from these seed points. Incomplete open-ends, or sections of arbor that were incomplete, were reconstructed by multiple users in an iterative process until the tips of all arbor were accounted for. (**D**) Left-right hemisphere homologous neuron pairs were morphologically compared to target review to regions that were likely to have reconstruction issues or errors of omission (missing branches for example). (**E**) The quality of left-right homologous pairs were assessed morphologically and using network-based connectivity metrics. Left and right hemisphere neurons were transformed into a shared space, followed by NBLAST comparison. The rank of NBLAST-predicted pairings was compared to the expert-annotated left-right pairings. A similar rank comparison was performed using connectivity information via a graph-matching-based vertex nomination scheme, which evaluated how likely neurons were to be matched to their annotated pair (see Methods for details). (**F**) The cable length, number of inputs (postsynaptic sites), and number of outputs (presynaptic sites) were compared between left and right homologs. The similarity between left-right pairs suggests that the reconstruction methodology generates reproducible results. (**G**) Assessment of targeted review methodology compared to traditional full review of all neuron arbor. A random set of 10 brain interneurons were selected for full review. The connectivity of these neurons before and after full review was compared. (**H**) The majority of edges in these neurons did not change after full review. Edges that did change tended to increase, suggesting that most errors were of omission.

**Fig. S2.**
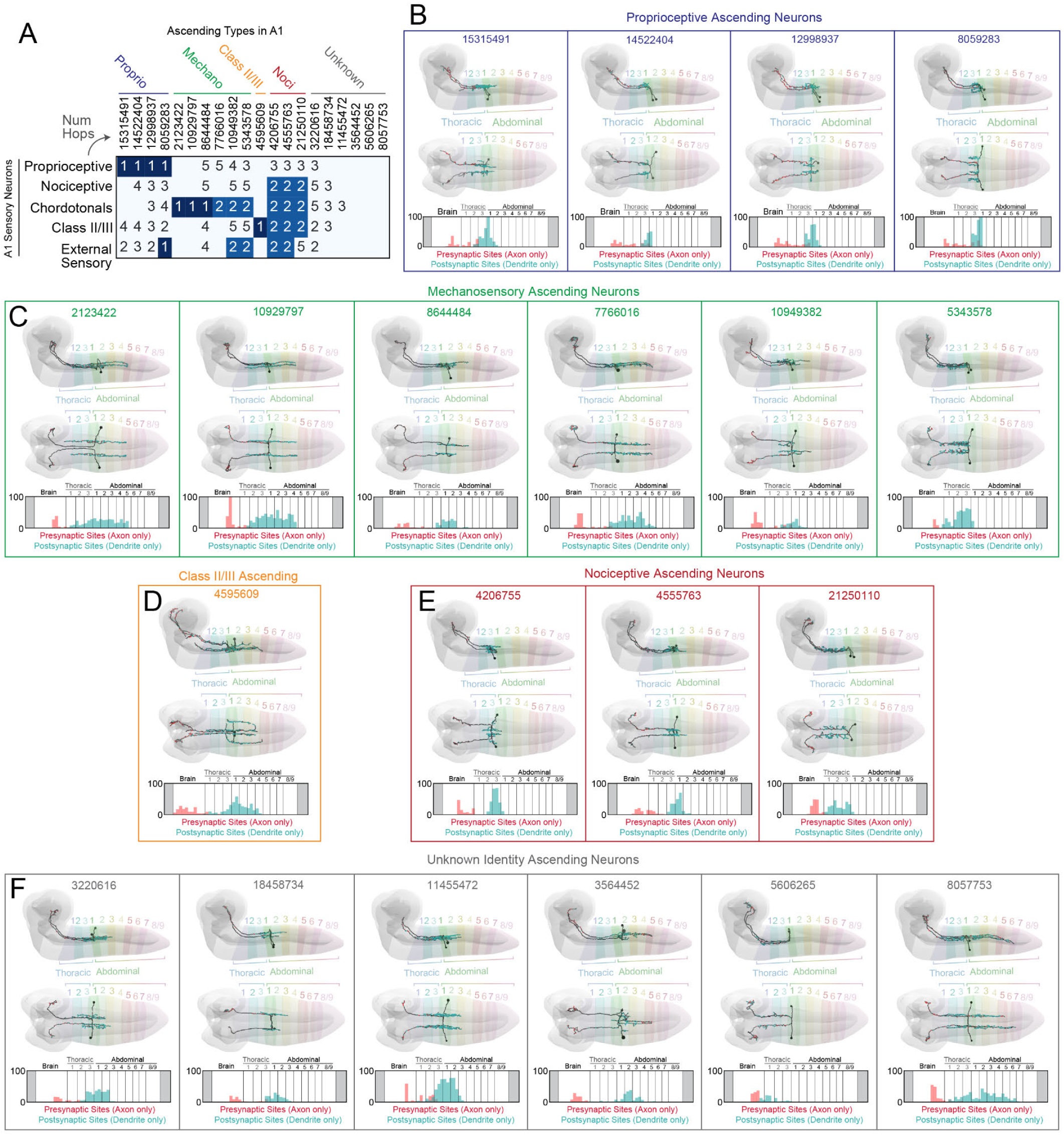
Overview of ascending neurons from A1. (**A**) Multihop connectivity matrix between SN types in A1 (rows) and individual AN pairs (columns, IDs correspond to left-side neuron ID in CATMAID). Numbers in this matrix indicate the number of hops between row and column neurons (i.e. 1 indicates a direct connection, 2 indicates a 2-hop connection, etc.). Each hop must be a strong, bilaterally symmetrical connection with ≥1% input onto the dendrite to be reported. We considered 1- and 2-hop connections to be salient and assigned putative modalities based on these connections. (**B** to **F**) Morphology of individual ascending neuron pairs grouped by sensory modality. The thoracic and abdominal segments are indicated in each plot, as well as the location of presynaptic sites in the axon and postsynaptic sites in the dendrite for each neuron in the anterior-posterior axis.

**Fig. S3.**
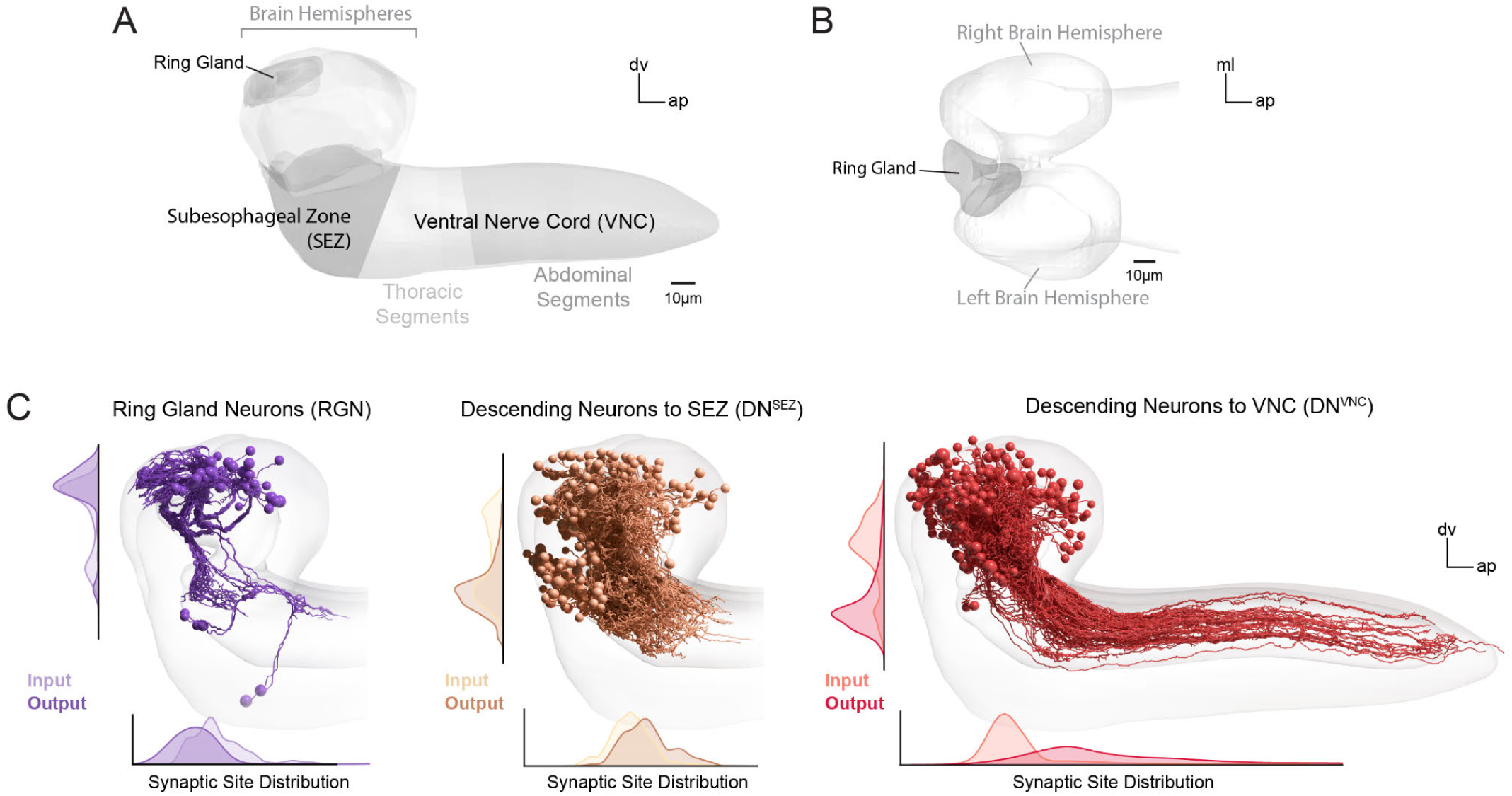
Overview of brain output neurons. (**A**) Rendered CNS regions from the EM volume, including both brain hemispheres, the ring gland, the SEZ, and the VNC. The boundary between the brain and SEZ was defined using cell bodies of ventral brain neurons. The brain itself was defined according to stereotyped lineage entry points (see Methods). The boundary between SEZ and VNC was defined based on neurohemal organs at the boundary (Fig. S22A). The image represents a side view of the CNS. dv=dorsoventral axis, ap=anteroposterior axis. (**B**) A top-down view of the ring gland in the brain, which is positioned between the two hemispheres on the dorsal side. ml=mediolateral axis, ap=anteroposterior axis. (**C**) The three brain output types were categorized based on location of axon presynaptic sites. RGNs displayed axonal outputs in the ring gland, DN^SEZ^ displayed axonal outputs in the SEZ, and DN^VNC^ displayed axonal outputs in the VNC. Distributions of all postsynaptic sites (input) and presynaptic sites (output) in both axon and dendrites are plotted for each output type.

**Fig. S4.**
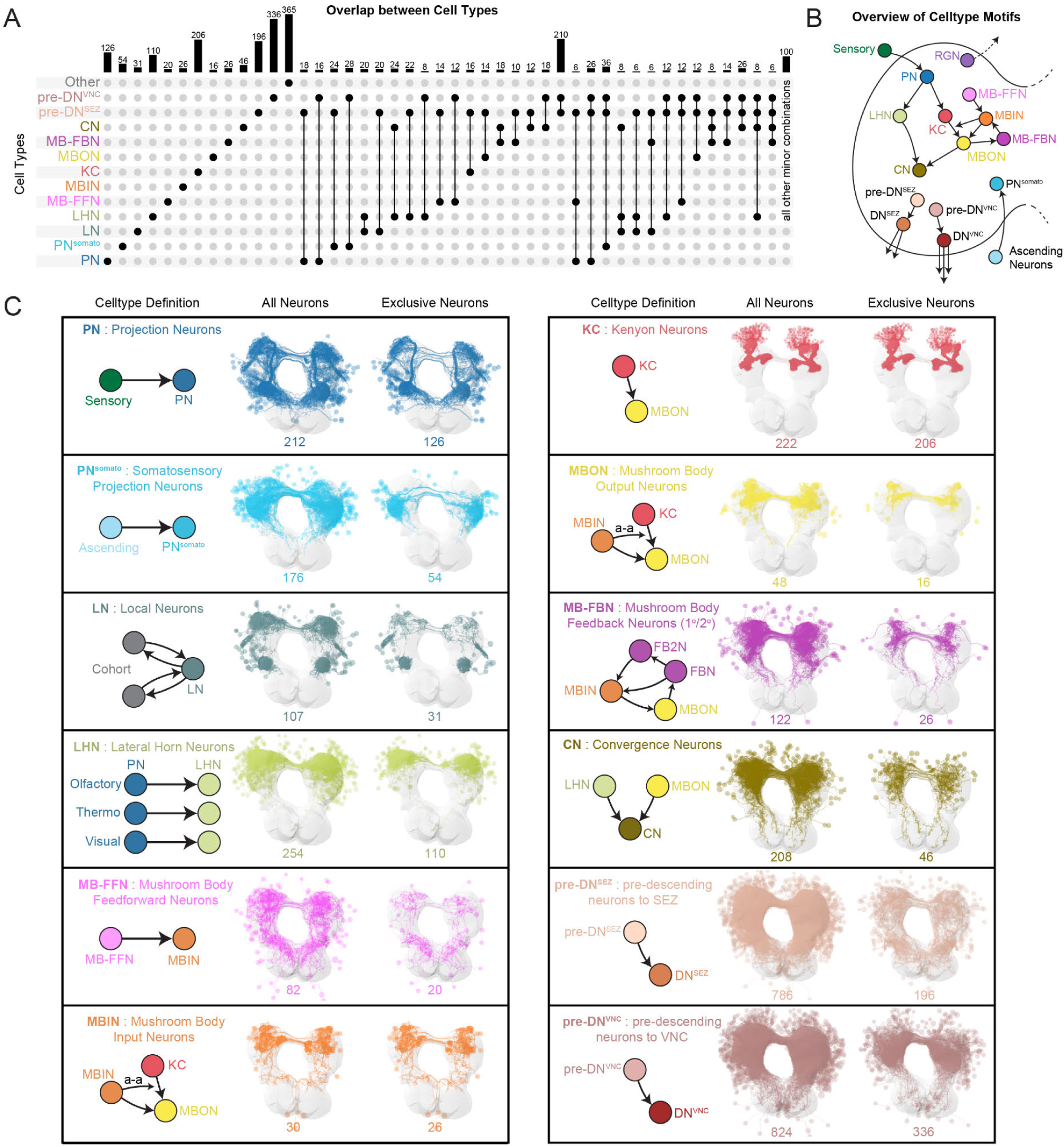
Overview of brain interneurons. (**A**) Overlap between cell-type classifications. Cell-type classifications of many neurons were unique, but neurons with multiple classifications also occurred. Throughout this paper, interneuron classes are displayed as mutually exclusive for plotting expedience, based on the following priority: [SN, AN, LN, MBIN, KC, MBON] > [PN, DN^VNC^] > [MB-FBN] > [LHN] > [CN] > [DN^SEZ^, PN^somato^] > [RGN, MB-FFN] > [pre-DN^VNC^] > [pre-DN^SEZ^]. For example, a neuron that is both MBON and pre-DN^SEZ^ is plotted as a MBON in future figures. Cell classes within brackets displayed no overlap. (**B**) Schematic of connectivity between cell classes in the brain, based on prior studies (*25, 31, 45, 92*) and cell classes defined in the present study, including LN, PN^somato^, pre-DN^SEZ^, and pre-DN^VNC^. (**C**) Cell classes and their connectivity definitions (left columns). All brain neurons that met each cell class definition were identified. Neuron morphologies of each cell type are displayed (center columns) or only the neurons that exclusively belong to the respective cell type (right columns). Arrows indicate a-d connections, unless labeled a-a. LNs were identified using definitions explained in Fig. S9A, B.

**Fig. S5.**
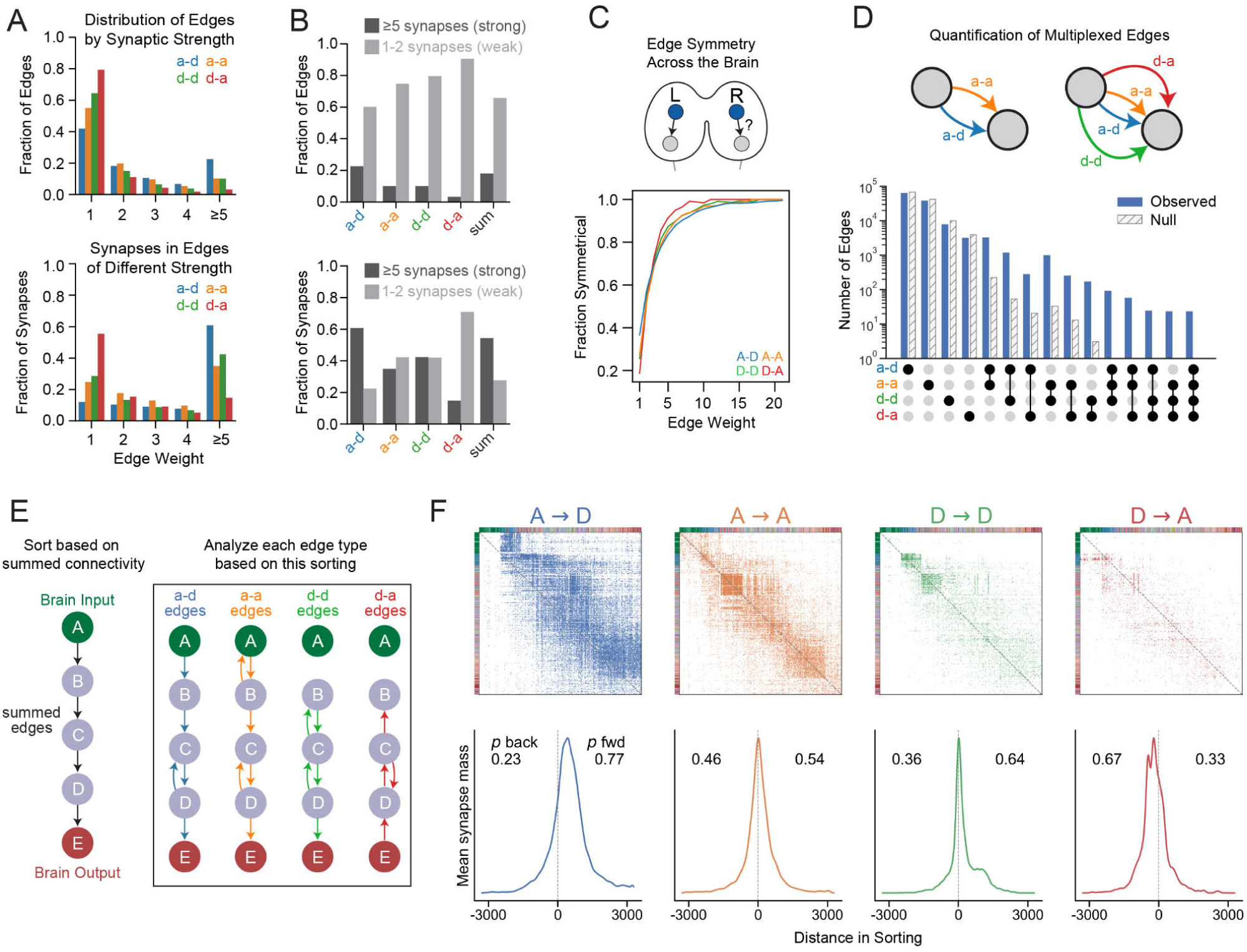
Detailed analysis of four connection types. (**A**) Distribution of edges and synapses based on edge weight (number of chemical connections between neurons). Axo-dendritic (a-d) connections displayed stronger edge weights than other edge types. (**B**) Fraction of edges or synapses in strong (≥5 synaptic strength) or weak edges (1-2 synaptic strength), per edge type (sum = the summed graph, all edge types together). (**C**) When an edge was observed in one brain hemisphere, we determined whether a homologous edge existed in the opposite hemisphere. This edge symmetry was quantified across synaptic strengths. If matching edges existed in both hemispheres, they were considered symmetrical regardless of edge strength. Stronger edges were more likely to have a homologous edge in the opposite hemisphere than weak edges. (**D**) Number of multiplexed edges in the connectome, i.e. two or more edges that both originate from one neuron and terminate onto another neuron. Most edges were singletons, also observed in a Erdos-Renyi null model (see Methods, Multiplexed Edge Analysis). However, there were many multiplexed edge types, including a-d/a-a edges and rare 4-type edges, and these multiplexed edges were observed more frequently than expected by the null model. Comparisons between the observed and the expected number of edges under the null model were significantly different (p < 10^-37^, binomial test) for each of the edge type combinations. (**E**) Methodology to identify feedforward and feedback edges in each connection type. The Walk-Sort algorithm was applied to the summed graph (all edge types together), resulting in a sorting from SNs to DNs. This sorting was then applied to all graph types. Feedforward and feedback edges were quantified based on their anterograde or retrograde direction in respect to the overall graph sorting. (**F**) Adjacency matrices of each connection type with sorting from the summed graph applied. The mean synaptic mass above the diagonal (feedforward) or below the diagonal (feedback) was quantified. The dotted line on the diagonal of each adjacency matrix corresponds to the dotted line in each line plot directly below. The a-d graph displayed the most feedforward synapses, while the d-a graph displayed the most feedback synapses.

**Fig. S6.**
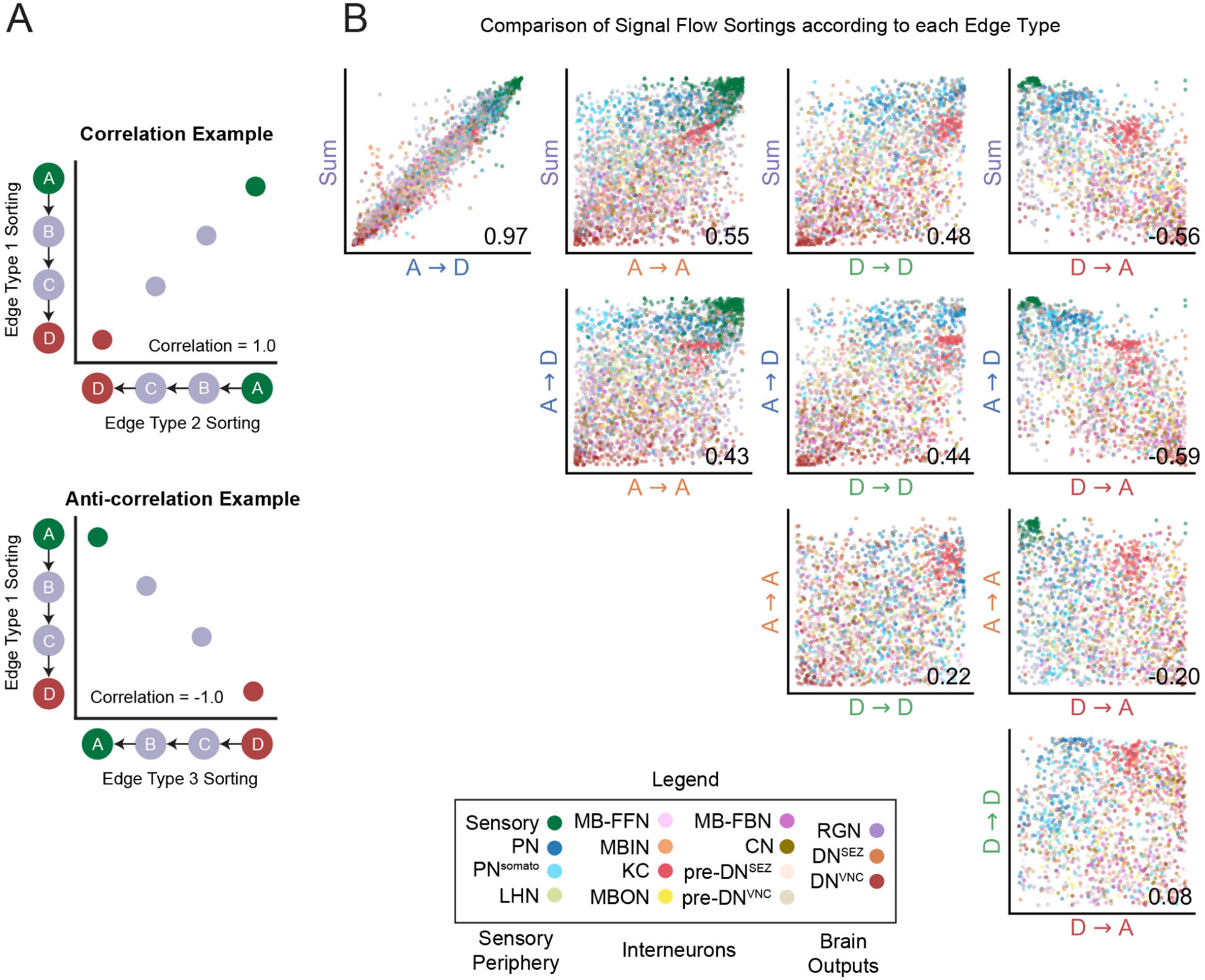
Signal flow sorting for each edge type in the connectome. (**A**) Schematic examples of comparison of signal flow sortings between graphs with different edge types (in a hypothetical graph with 4 nodes/neurons). Graphs comprising each edge type are sorted independently using the signal flow algorithm. These sortings are compared and the correlation between sortings is quantified. An example of correlation (*top*) and anti-correlation (*bottom*) are displayed. (**B**) Comparison of signal flow sortings for each graph type. Individual dots correspond to single cells, colored based on cell classes. The rank correlation between these node sortings was quantified for each pairwise comparison and displayed in the bottom right of each plot. We found that the axo-dendritic sorting best matched the summed graph (all edge types together), sorted from sensory neurons (green; top right) to output neurons (red, bottom left). The dendro-axonic sorting was negatively correlated with the axo-dendritic (−0.63) and summed sortings (−0.55).

**Fig. S7.**
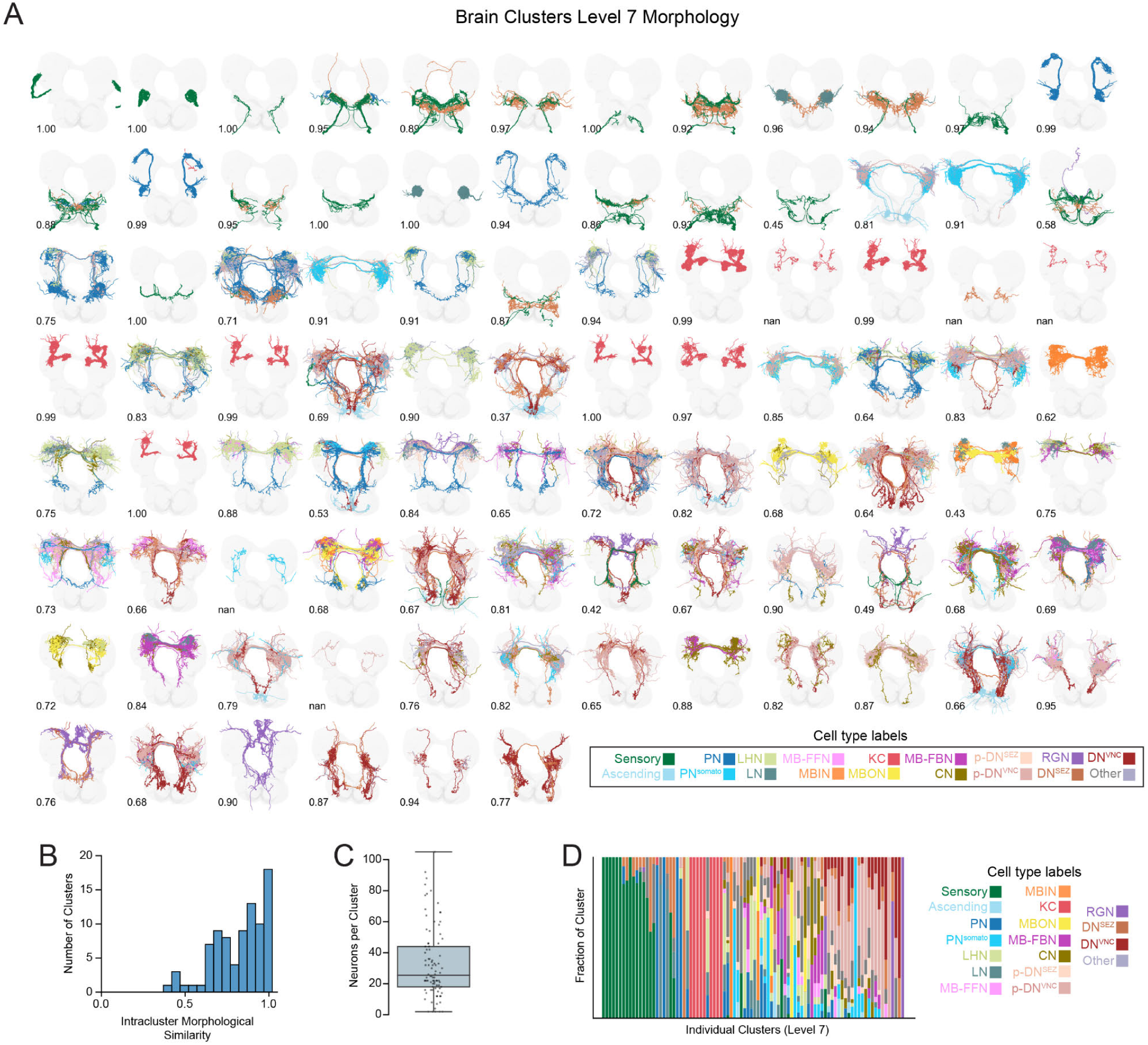
Overview of level 7 brain clusters. (**A**) Morphology of level 7 brain clusters. Neurons within each cluster are colored based on cell class (legend, bottom right). The intracluster morphological similarity was quantified using averaged within-cluster NBLAST scores (see Methods) and is displayed on the left bottom of each cluster plot. Note that similarity cannot be calculated (nan) for a small number of clusters, because they only contain a single neuron pair. (**B**) Distribution of intracluster morphological similarity. Most clusters display a high level of morphological similarity between their constituent neuron members. Note that using this metric, 0.5 corresponds to the expected similarity between any two randomly chosen neurons. (**C**) Distribution of neuron counts per cluster. (**D**) Fraction of neuron classes per level 7 cluster. Clusters are sorted from SNs to DNs.

**Fig. S8.**
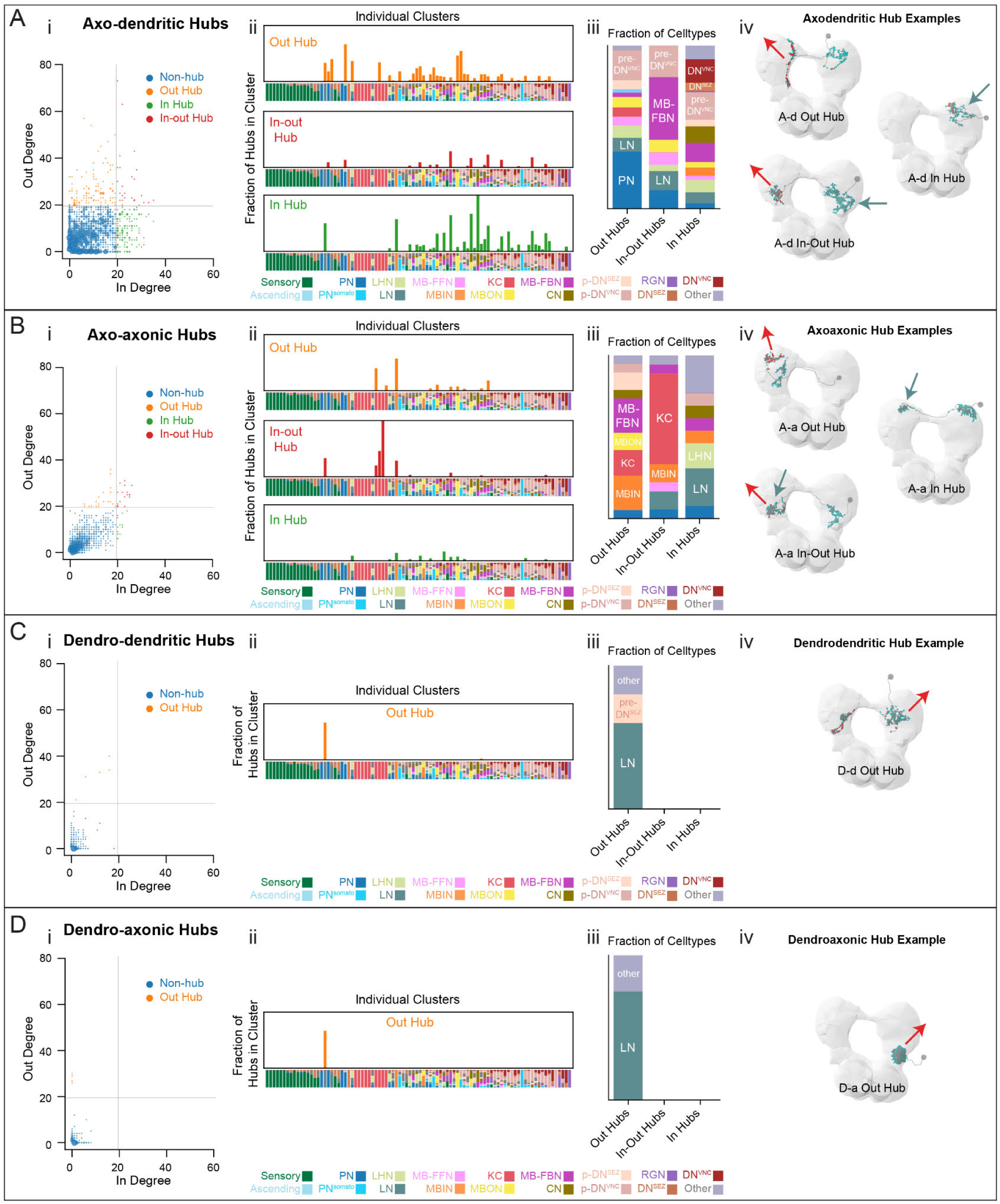
Characterization of all brain hub neurons. (**A**) Out hubs (≥20 out-degree), in-out hubs (≥20 out-degree and ≥20 in-degree), and in hubs (≥20 in-degree) in the a-d graph (i). In degree is the number of strongly connected presynaptic partners, while out degree is the number of strongly connected postsynaptic partners. The locations of hubs in the brain cluster structure (ii), their neuron class identities (iii), and some example morphologies (iv) are depicted. (**B**) Out hubs, in-out hubs, and in hubs in the a-a graph. (**C, D**) Out hubs in the d-d and d-a graphs.

**Fig. S9.**
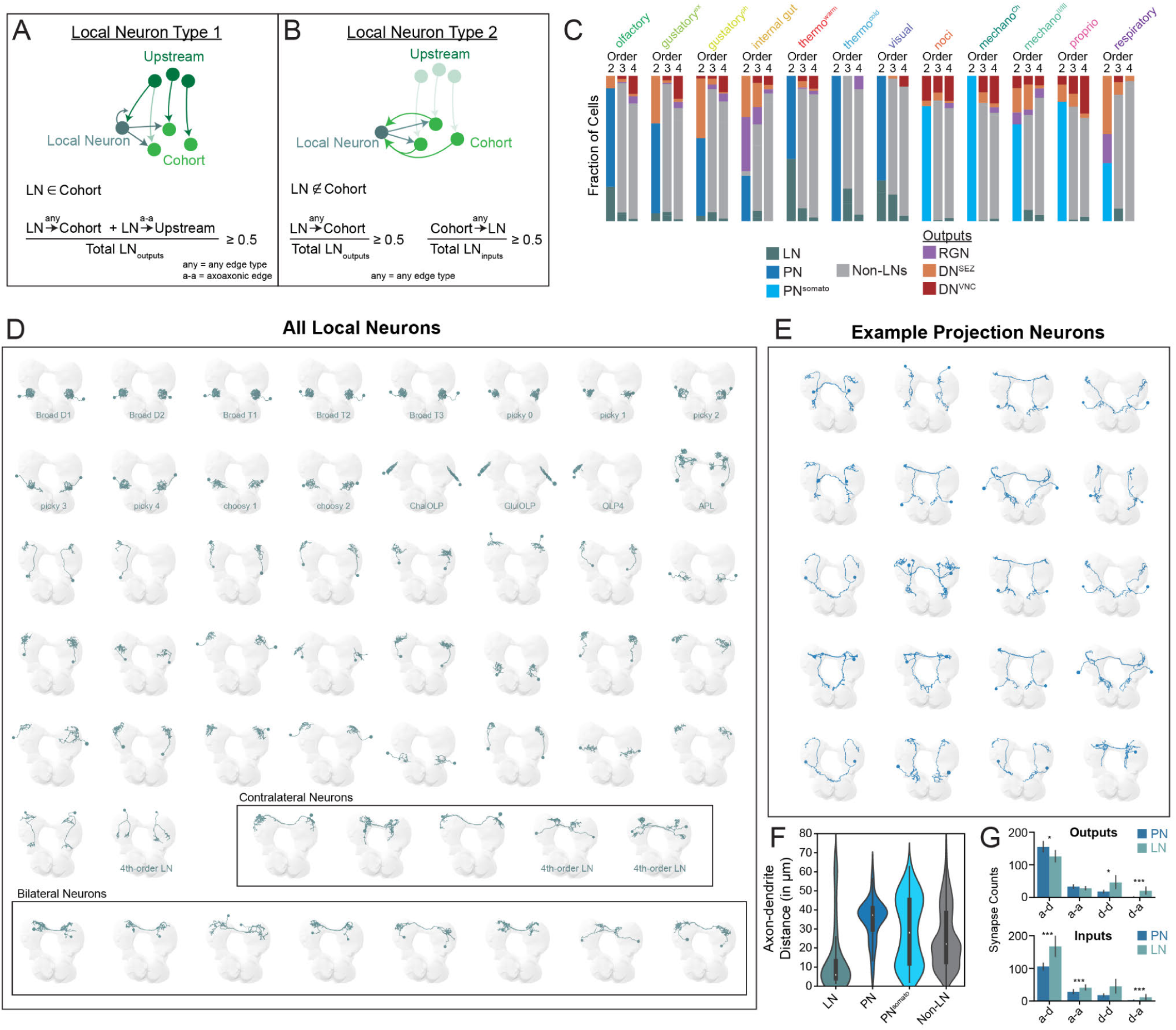
Identification of local and projection neurons in the brain. (**A**) Definition of a local neuron (LN) within a cohort (type 1), based on LNs observed in olfactory and visual neuropils (*45, 46*). Type 1 LNs sent most of their output to neurons within their own sensory cohort (i.e. neurons that are the same number of hops away from SNs of a particular sensory modality, see Fig. 4A-C, Fig. S10A) or to neurons upstream of that cohort via a-a connections. (**B**) Definition of a LN outside of a cohort (type 2), based on LNs observed in the mushroom body, specifically APL and its interactions with KCs (*25*). Type 2 LNs received most of their input and sent most of their output to a cohort of neurons, to which they do not belong. Sensory circuits were again defined as cohorts (Fig. 4A-C, Fig. S10A). (**C**) LN types were identified in all sensory circuits. Most sensory circuits contained LNs, but higher-order circuits and somatosensory circuits displayed less LNs than circuits directly downstream of brain sensory neurons. Because there is strong overlap in 4th-order neuropils, 4th-order LNs were shared between different modalities. Projection neurons (PN) were defined by exclusion in 2nd-order circuits (all neurons that were not local or brain output neurons). PNs were postsynaptic of SNs, while PNs^somato^ were postsynaptic of somatosensory ANs. (**D**) Morphology of all LN pairs. Previously identified LNs are annotated (Broad LNs, picky LNs, choosy LNs, ChalOLP, GlulOLP, OLP4, and APL), as well as one 4th-order LN pair. (**E**) Morphology of example PNs. (**F**) The axon-dendrite distance was longer in PNs and PNs^somato^ compared to LNs, using the centroids of axon presynaptic sites and dendrite postsynaptic sites. (**G**) Connection types observed in PNs and LNs. LNs exhibited significantly more noncanonical outputs and inputs. Mann Whitney U test, p-values: * < 0.05, *** < 0.001

**Fig. S10.**
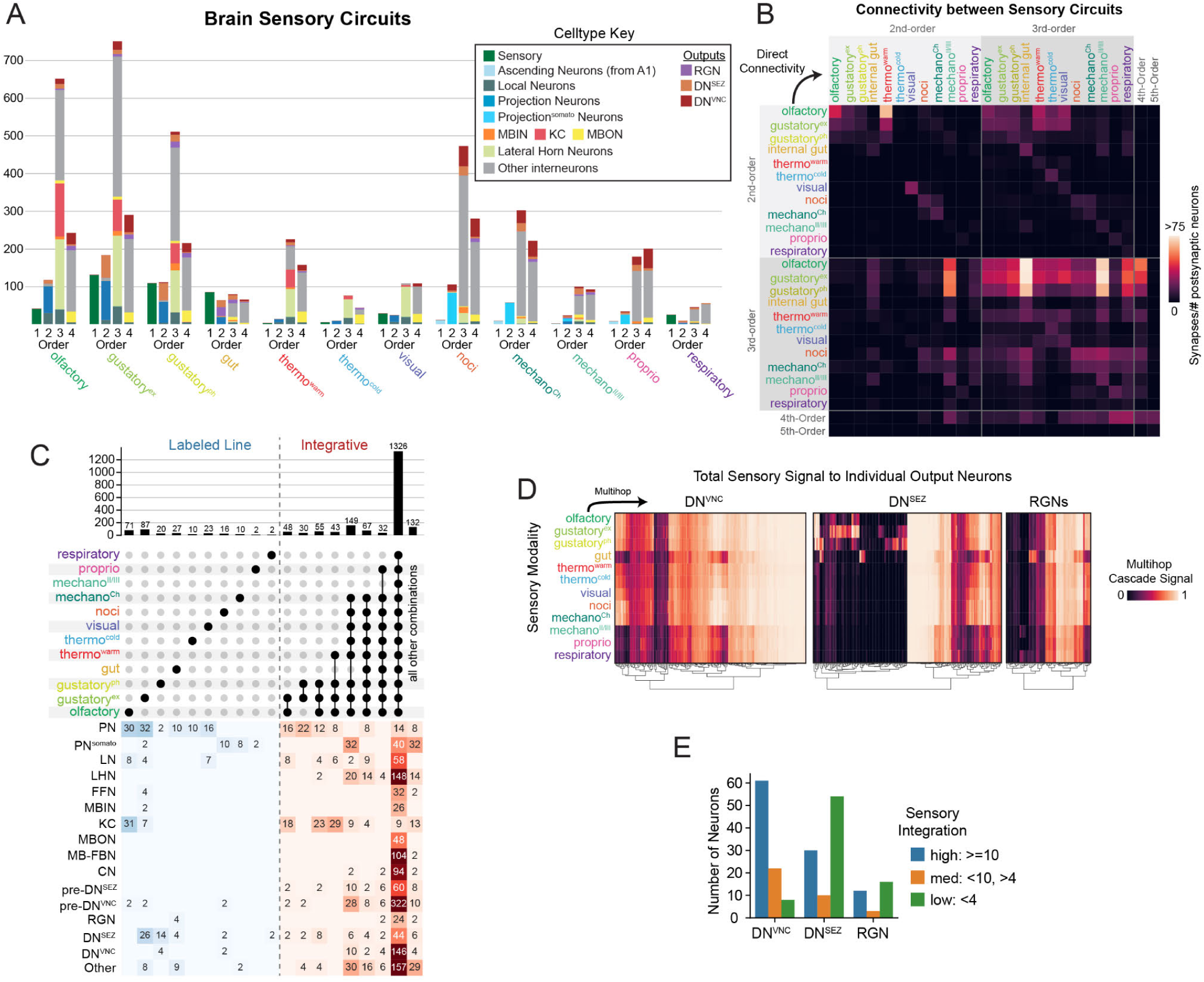
Detailed analysis of multimodal integration. (**A**) Overview of input neurons, 2nd-, 3rd-, and 4th-order neurons for each sensory modality (identified using a-d connectivity). The number of neurons in each category is quantified and color-coded by neuron classes. (**B**) Direct a-d connectivity between sensory circuits, displayed as the summed number of synapses between neuron groups, normalized by the total number of postsynaptic cells. (**C**) Celltype breakdown of each combination of input from multihop a-d sensory cascades. (**D**) Multihop a-d signal from each sensory modality to individual brain output neurons. (**E**) Fraction of different output types that display high, medium, or low sensory integration. Most DNs^VNC^ were highly integrative, while many DNs^SEZ^ and RGNs displayed low sensory integration.

**Fig. S11.**
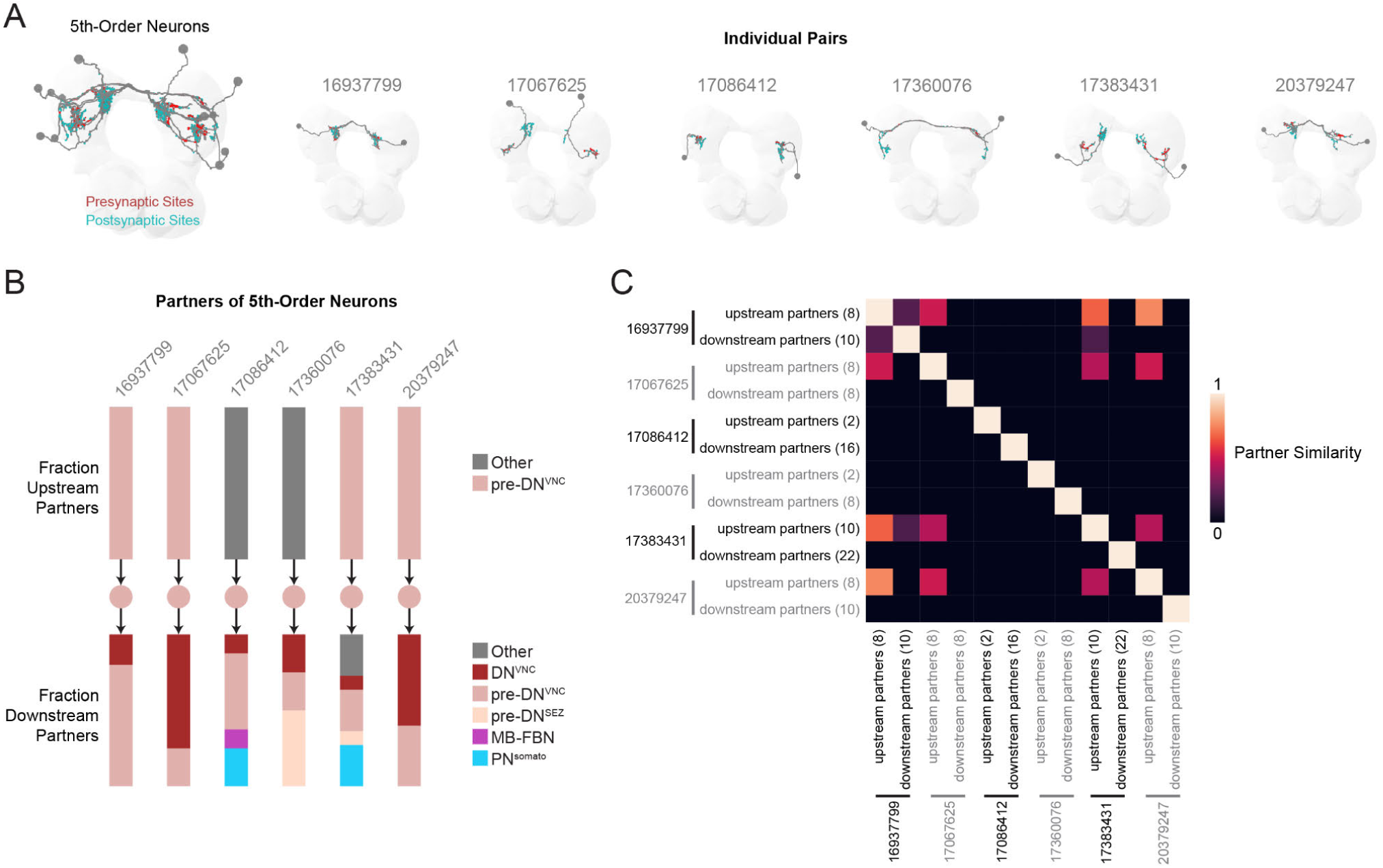
Overview of 5th-Order Neuron Morphology and Connectivity. (**A**) Morphology of all 5th-order neurons, plotted together or as left/right homologous pairs. (**B**) Neuron classes of each 5th-order neurons’ presynaptic and postsynaptic partners (using a ≥1% input threshold of a-d connectivity). All 5th-order neurons were pre-DN^VNC^ neurons and therefore synapsed onto DNs^VNC^. Connections were also observed onto other pre-DN^VNC^, pre-DN^SEZ^, and occasionally PN^somato^ neurons. One 5th-order neuron pair directed synapse onto FBN-7, an a-d in-out hub (Fig. 3F) involved in feedback in the larval learning and memory center. (**C**) The similarity between each 5th-order neuron’s downstream and upstream partners was compared using the Dice Coefficient, such that 0 indicates no shared partners and 1 indicates that partners are exactly the same.

**Fig. S12.**
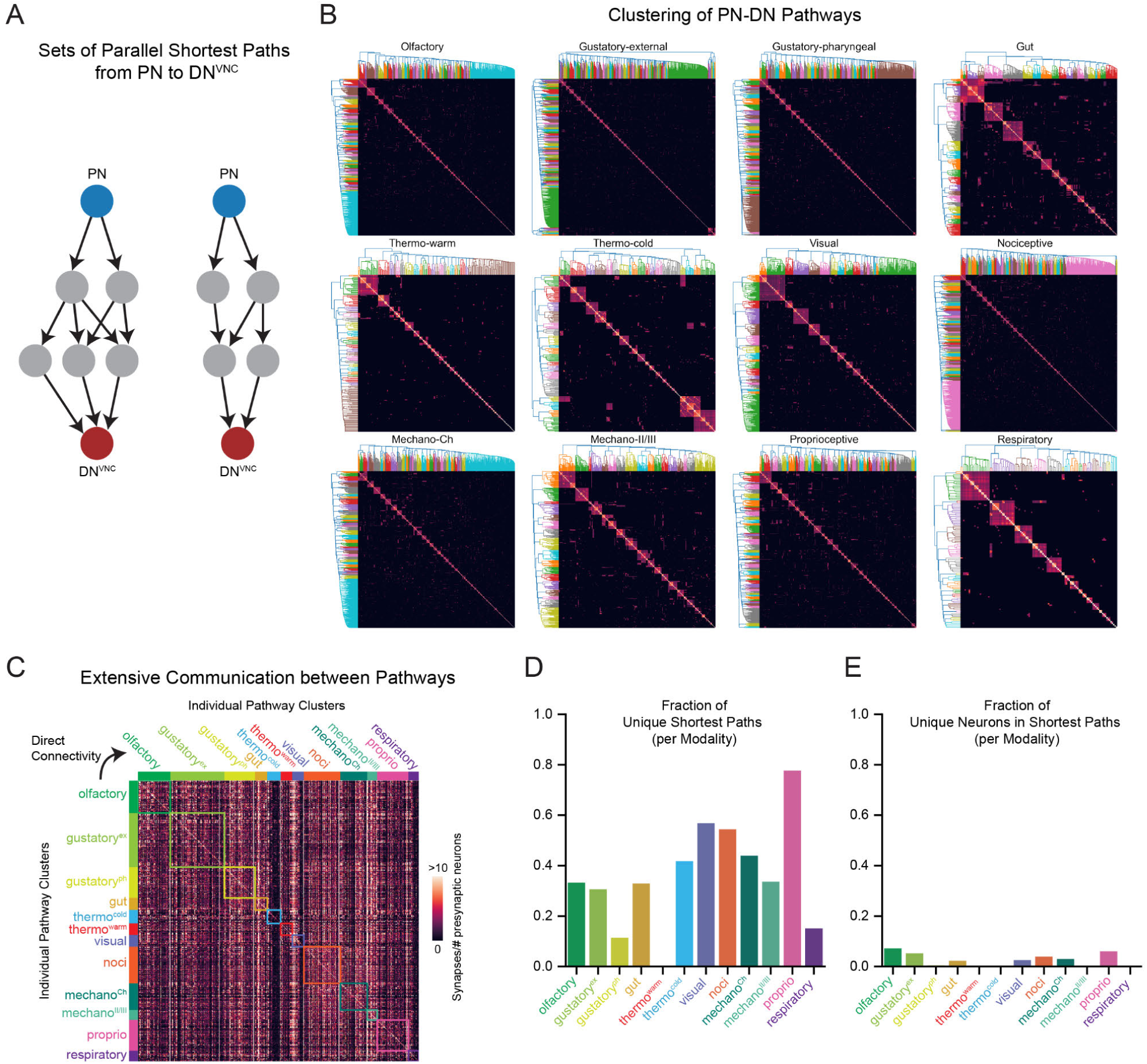
Shortest Pathways from Projection to Descending Neurons for each Sensory Modality. (**A**) Parallel shortest pathways from PN to DNs^VNC^ for a hypothetical sensory modality, using a-d connectivity. Pathways often shared neurons, but it was possible to cluster groups of related parallel pathways from the same sensory modality. (**B**) Clustering of PN-DN^VNC^ shortest pathways based on Dice Coefficient comparison of neuron members within each pathway. Each sensory modality displayed a variety of pathway clusters, which may be thought of as parallel pathway types. (**C**) Extensive a-d connectivity was observed between neurons within pathway clusters for each sensory modality (boxed areas in matrix), as well as between pathway clusters across sensory modalities (all areas outside of boxes). (**D**) Unique pathways were identified across sensory modalities (i.e. pathways only observed in a single modality). We found many unique pathways, but in 9/12 sensory modalities, most pathways were not unique. (**E**) We found that the vast majority of neurons found in PN-DN^VNC^ pathways were not unique across modalities, consistent with the finding that most neurons are multimodal (Fig. 4G).

**Fig. S13.**
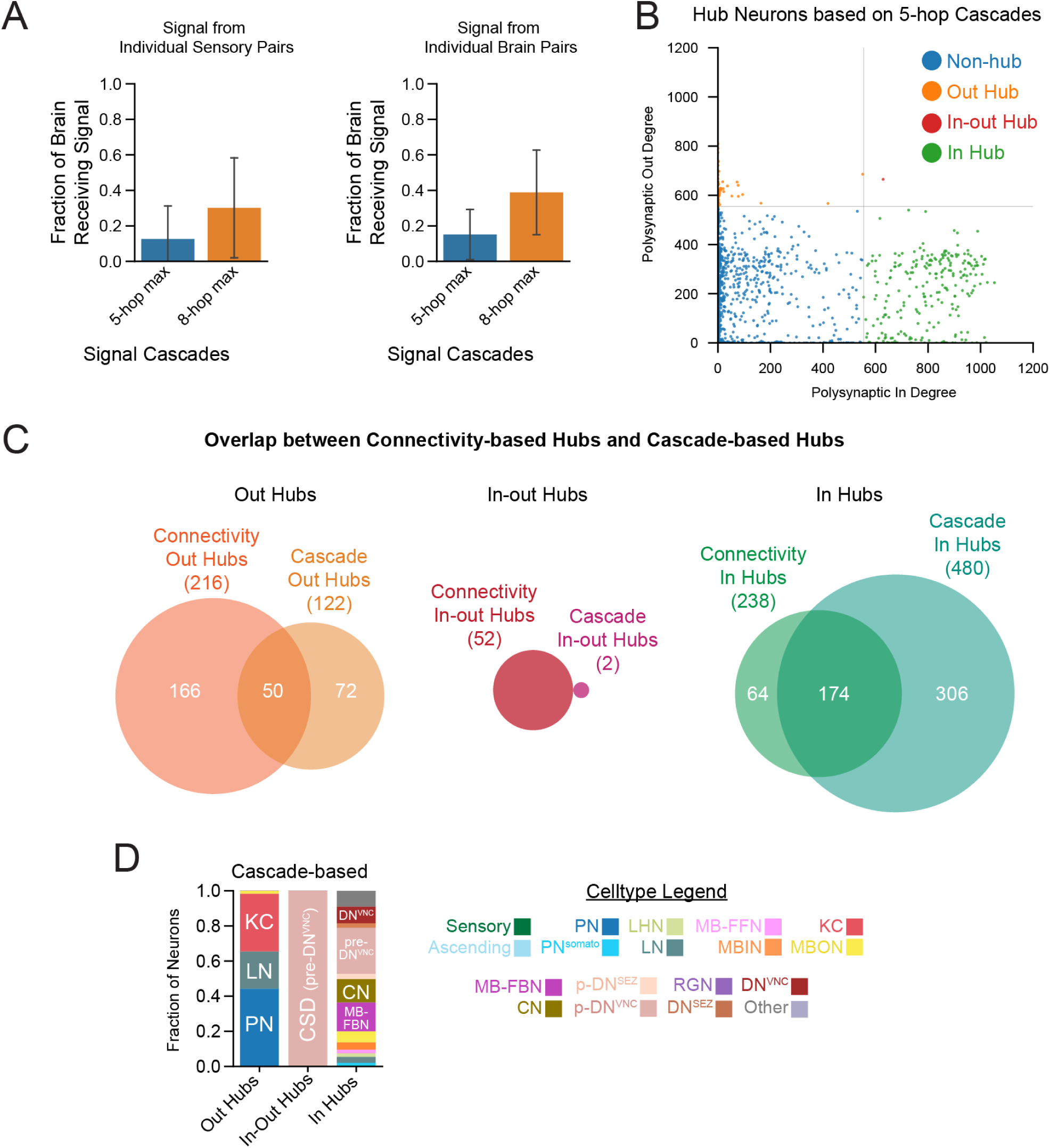
Overview of signal cascade brain coverage and cascade-based hubs. (**A**) Quantification of the fraction of brain neurons that receive 5-hop or 8-hop max cascade signals using a-d connectivity, starting from individual sensory pairs (*left*) or individual brain neuron pairs (*right*). (**B**) Cascade-based hub neurons were identified using a polysynaptic in or out degree threshold (mean degree plus 1.5 standard deviations). Polysynaptic in and out degrees refer to the number of upstream or downstream neurons using 5-hop signal cascades. (**C**) Set intersections between connectivity-based hubs and cascade-based hubs. (**D**) Celltype identities of cascade-based hubs. Note that there is only one pair of cascade-based in-out hubs, namely the serotonergic neuron CSD (also a pre-DN^VNC^).

**Fig. S14.**
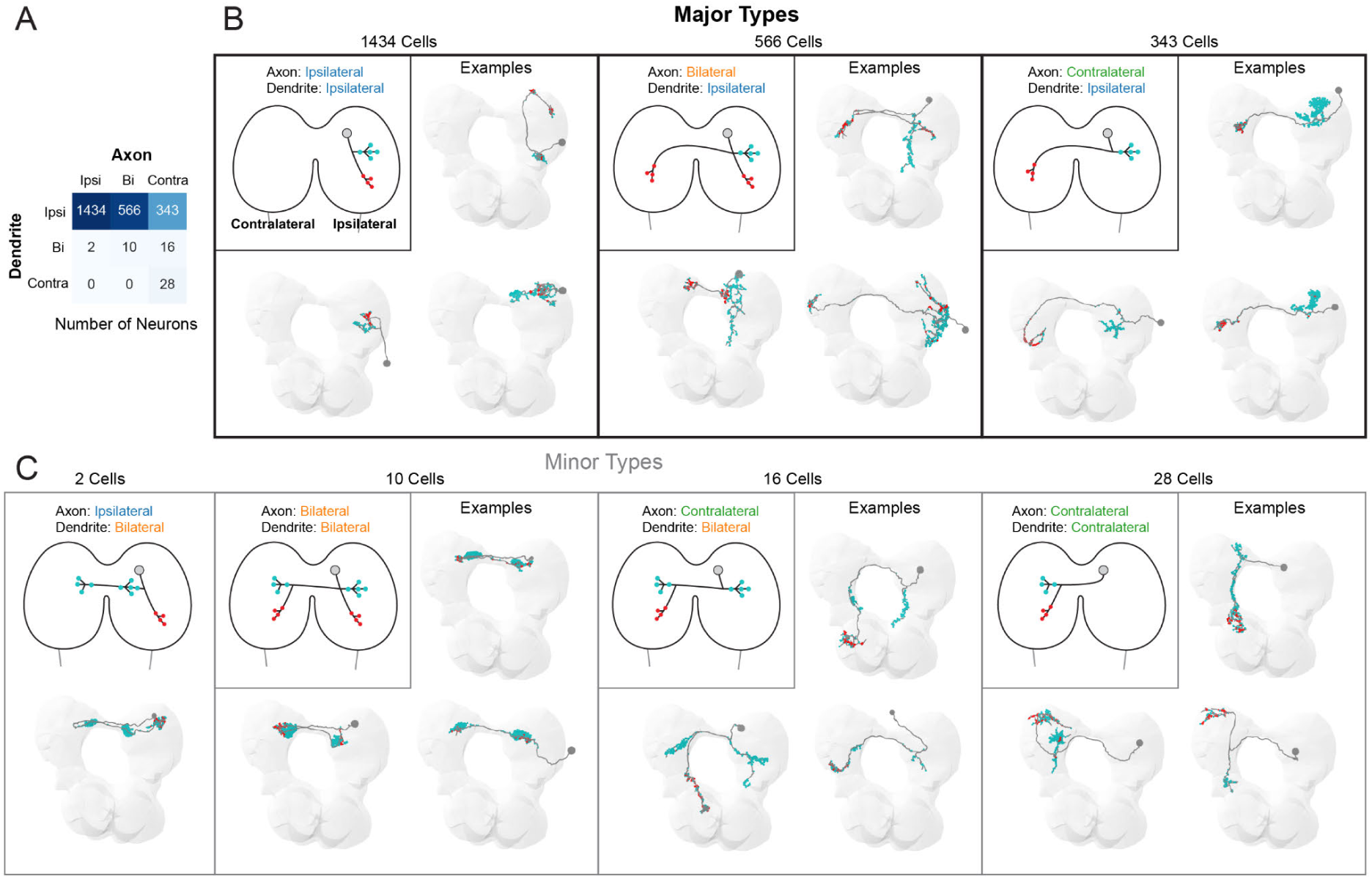
Interhemispheric characterization of all axons and dendrites. (**A**) Axons and dendrites were annotated as ipsilateral, bilateral, or contralateral, based on their relation to the brain hemisphere which contained the cell body. A majority of brain neurons displayed ipsilateral dendrites with a mixture of axon types. However, a small number of neurons displayed bilateral and contralateral dendrites. (**B**) Morphology of the major classes of neurons, which all contain ipsilateral dendrites. Schematic diagrams and the morphology a few examples are depicted for each category. (**C**) Morphology of the minor classes of neurons, containing bilateral or contralateral dendrites. Schematic diagrams and the morphology a few examples are depicted for each category. Neurons with contralateral axons and contralateral dendrites were treated as ipsilateral-ipsilateral neurons for interhemispheric analyses throughout this study, despite their unusual morphology.

**Fig. S15.**
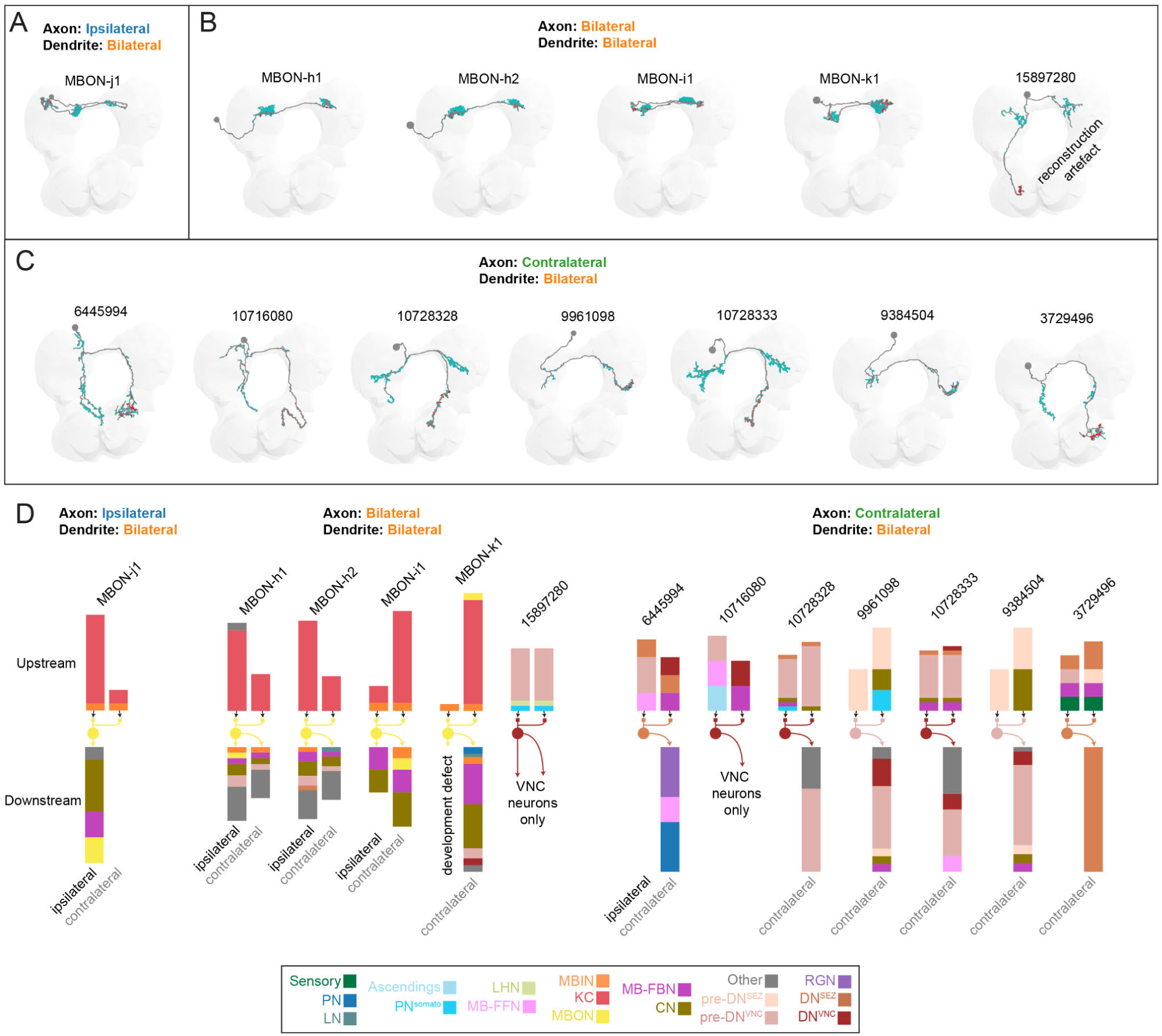
Characterization of neurons with bilateral dendrites. (**A**) Morphology of MBON-j1, the only example of an ipsilateral axon / bilateral dendrite neuron. (**B**) Morphology of all bilateral axon / bilateral dendrite neurons. Note that most are mushroom body (MB) output neurons (MBONs). (**C**) Morphology of all contralateral axon / bilateral dendrite neurons. Note that 5/7 pairs are DNs and extend axons into the SEZ or VNC. (**D**) Summary of direct a-d upstream and downstream partners. Barplots above each neuron indicate the fraction of neurons upstream of either the ipsilateral or contralateral dendrite of each neuron, such that both bars sum to 1. Barplots below each neuron indicate the fraction of neurons downstream of ipsilateral or contralateral portions of the axon. The name of the neuron of interest is displayed above each bar plot and the color of the neuron indicates its cell type. Note that the downstream VNC targets of DNs^VNC^ are not currently reconstructed in these cases, so only downstream neurons in the brain are displayed.

**Fig. S16.**
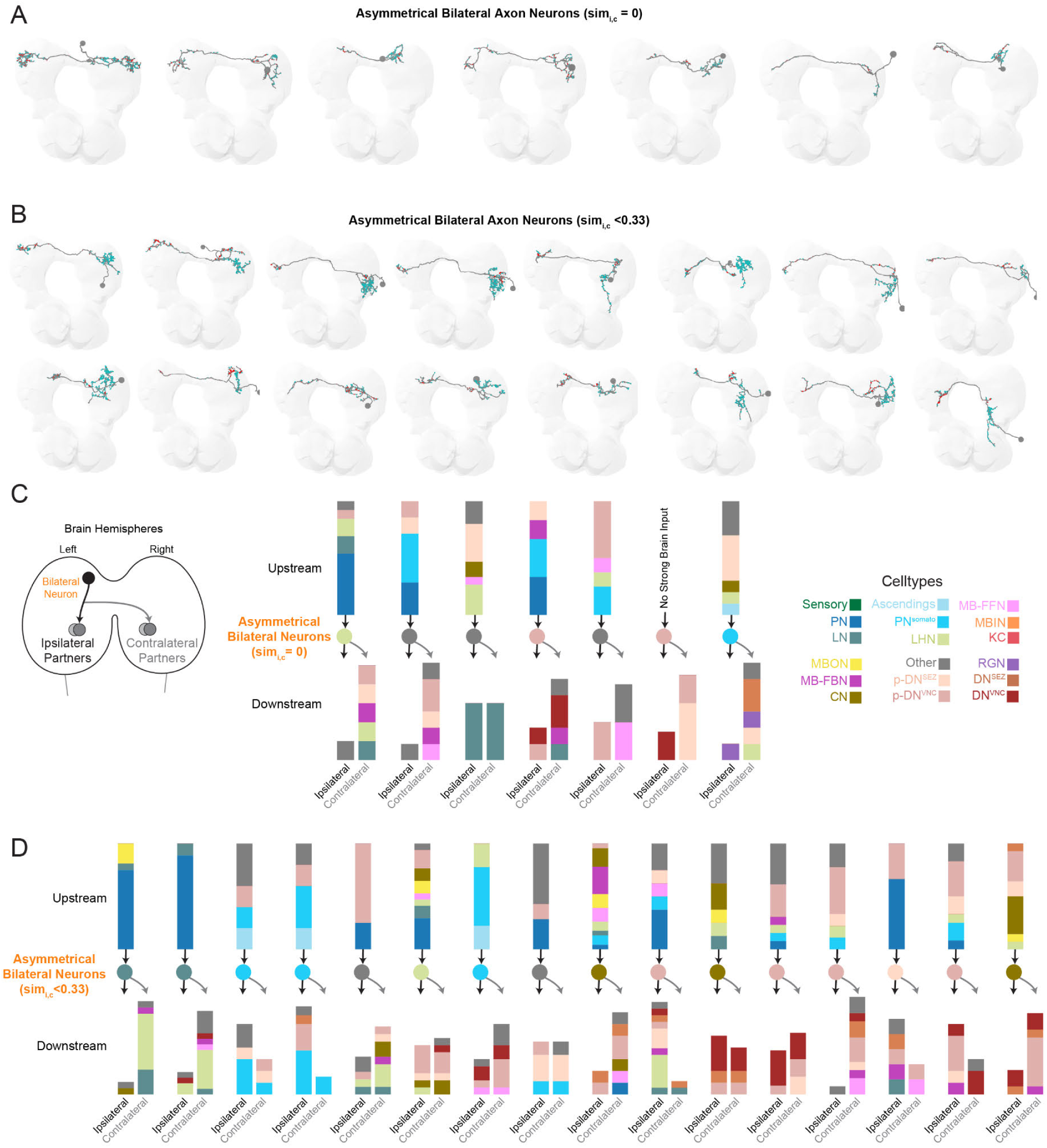
Characterization of neurons with asymmetrical bilateral axons. (**A**) Morphology of asymmetrical bilateral axon neurons. (**B**) Morphology of partially asymmetrical bilateral axon neurons with a cosine similarity <0.33 between ipsilateral and contralateral edges. (**C**) Summary of all direct a-d upstream and downstream partners of asymmetrical bilateral axon neurons. All downstream ipsilateral and contralateral partners have different identities and here we demonstrate that many of the class identities are also different. (**D**) Summary of all direct a-d upstream and downstream partners of partially asymmetrical bilateral axon neurons. Many class identities were different between downstream ipsilateral vs. contralateral partners.

**Fig. S17.**
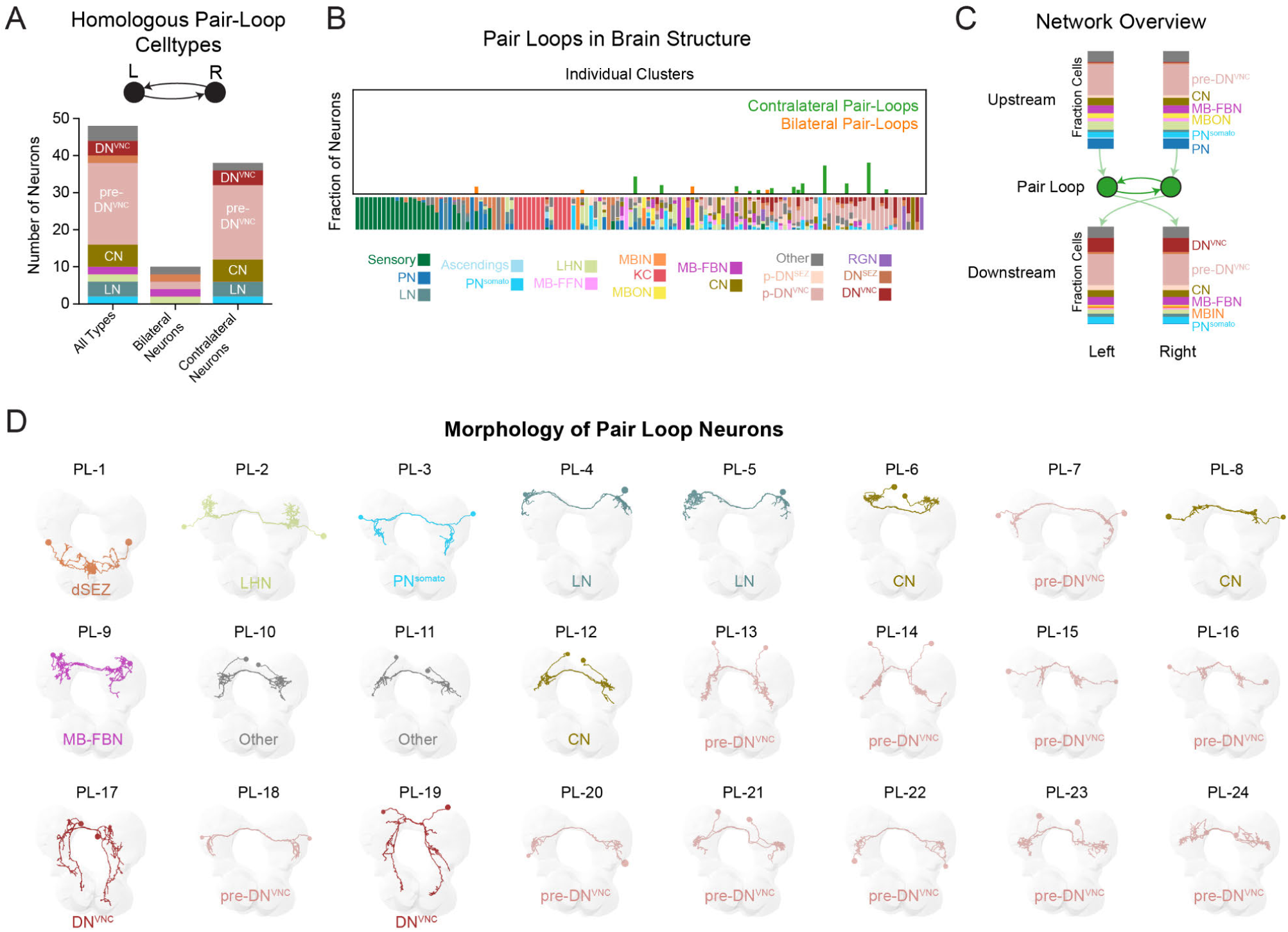
Overview of all reciprocal pair loops in the brain. (**A**) Cell types of homologous pair loops. (**B**) Location of pair loops within the brain cluster structure. Pair loops tend to be in deep brain regions. (**C**) Overview of all direct a-d upstream and downstream partners for all pair loops combined. (**D**) Morphology of each pair loop (PL) in the brain, colored and labeled by cell-type identity.

**Fig. S18.**
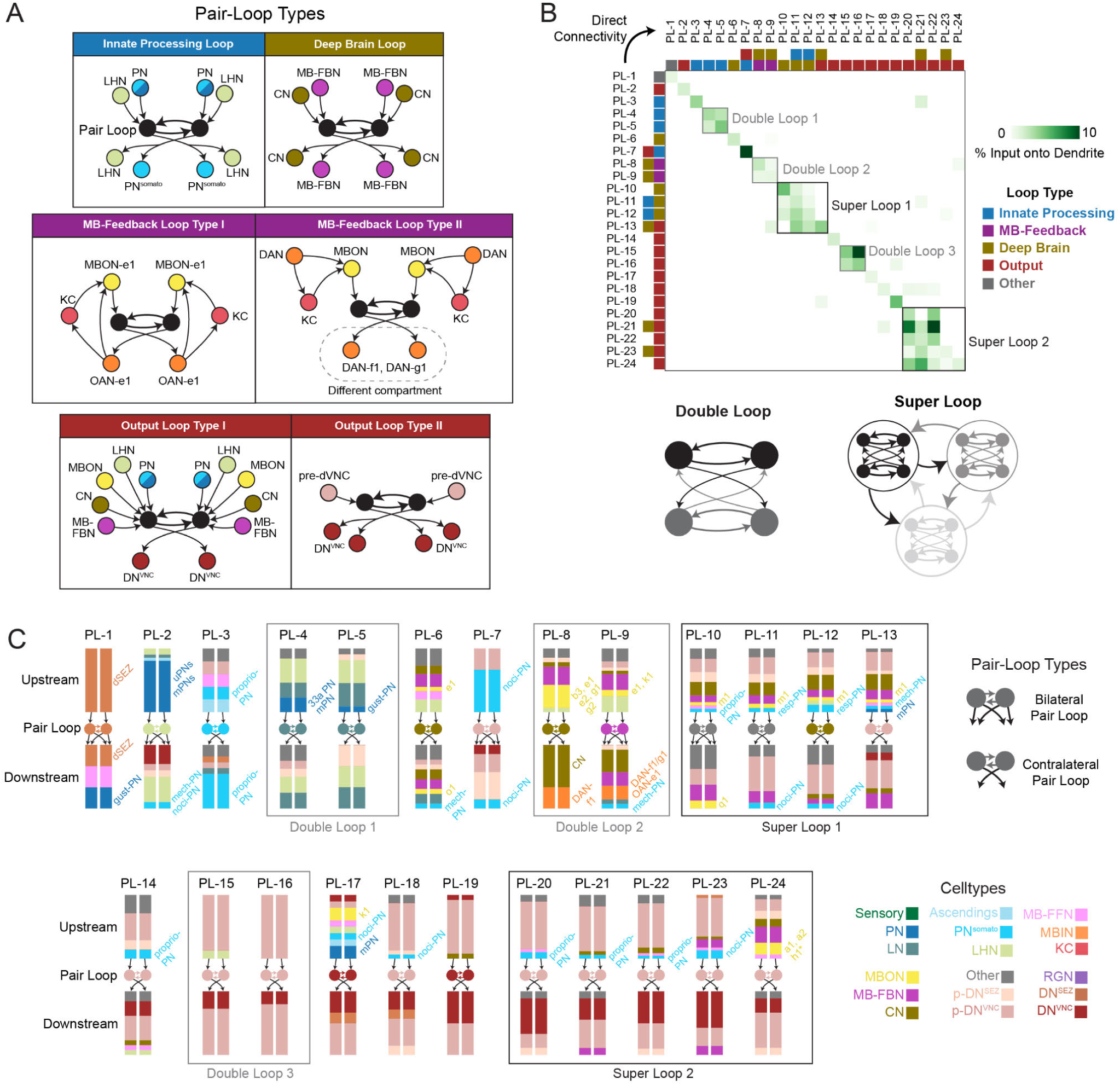
Characterization of motifs observed in reciprocal pair loops. (**A**) Common motifs observed across pair loops. Generally, pair loops propagated signals within sub-systems (innate processing loop, deep brain loop, and MB-feedback loops I/II) or propagated signals to the brain output system (output loop types I/II). (**B**) Adjacency matrix displaying a-d connectivity between all pair loops. We observed direct connectivity between multiple pair-loop types, including double loops (where two pair loops formed reciprocal connectivity between the four neurons within two pair loops) and super loops (where 4-5 pair loops formed complex reciprocal structures). (**C**) Summary of direct a-d upstream and downstream connectivity for each pair loop in both ipsilateral and contralateral hemispheres. Barplots indicate the fraction of neurons of a particular class in each upstream or downstream network. The double loops and super loops are boxed and correspond to those identified in B.

**Fig. S19.**
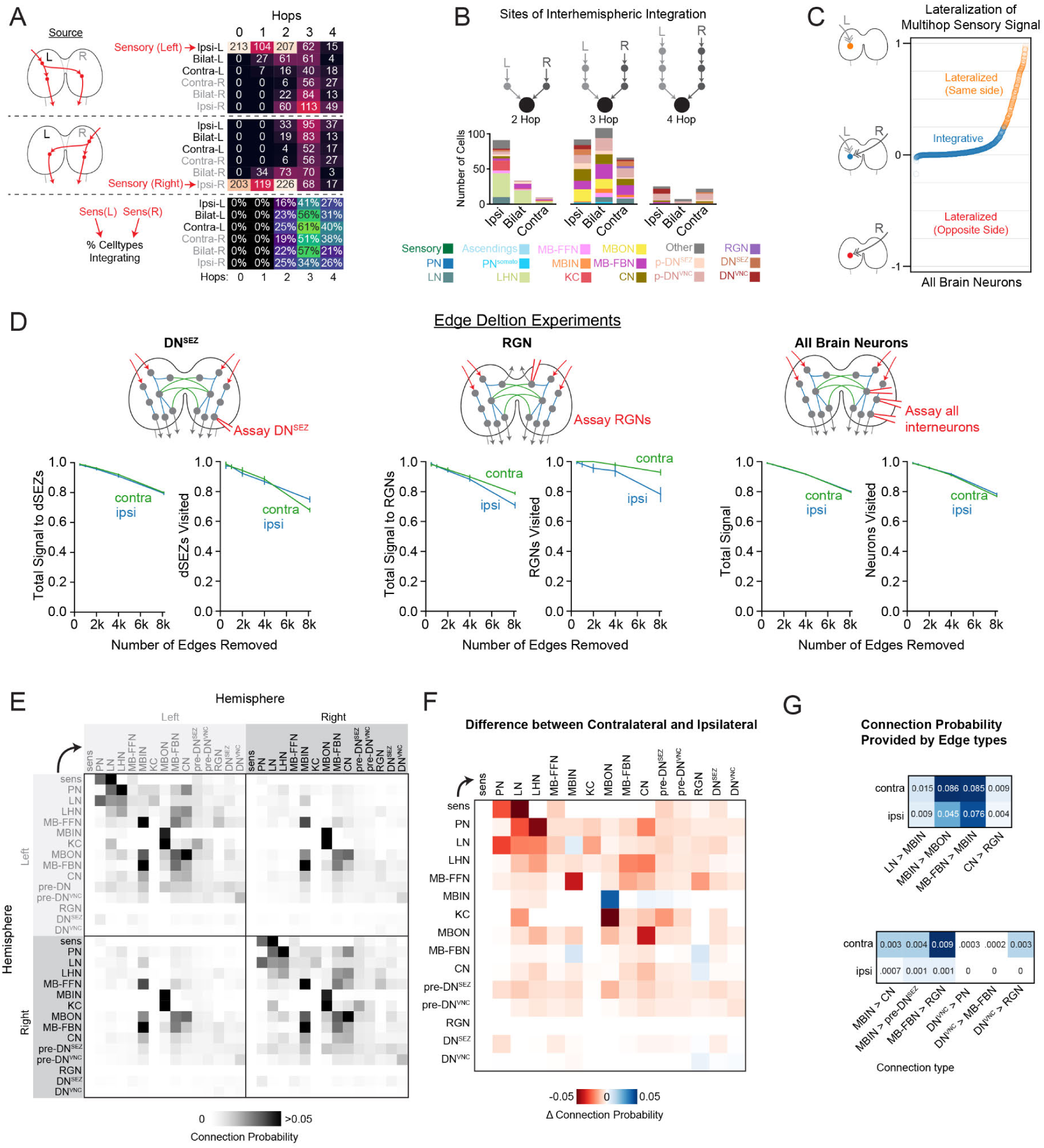
Characterization of contralateral edges in interhemispheric communication. (**A**) Signal cascades were generated from SNs on either the left or right hemisphere of the brain (using a-d connectivity). Left- and right-side ipsilateral, bilateral, and contralateral neurons were identified that received these signals within 4 hops (*top, middle*). Substantial signal to the contralateral hemisphere was already observed by hop 2. The percent of neurons that simultaneously integrate both left- and right-hemisphere sensory signals was quantified (*bottom*). Integration occurs as early as hop 2 but peaks at hop 3. (**B**) Cell types that integrate both left- and right-side sensory signals simultaneously at 2-, 3-, or 4-hops in the majority of signal cascade iterations (N=1000). (**C**) Quantification of sensory signal lateralization per neuron. Left- and right-signal cascades were generated from SNs using a-d connectivity. The left or right sensory signal each neuron received was quantified (regardless of whether these signals were coincident). The difference between ipsilateral and contralateral cascade signals was calculated for each neuron, such that 0 indicates equal signal from ipsilateral/contralateral sources, while +1 or −1 indicates signal only from ipsilateral or contralateral SNs, respectively. The majority of the brain integrates both left and right sensory signals, while some neurons receive lateralized signal from ipsilateral SNs. No neurons received only contralateral signal. (**D**) The role of ipsilateral and contralateral edges was tested by generating multihop a-d signal cascades from left and right brain inputs simultaneously. Random sets of ipsilateral or contralateral edges were removed from the graph before running these cascades. Total signal reaching each indicated neuron type and the total number of neurons visited were then assayed. Both contralateral and ipsilateral edges were important for signal to reach DNs^SEZ^. The same was true for RGNs, but ipsilateral edges appeared to be more important than contralateral edges. Ipsilateral and contralateral edges were both important for sensory signal to reach brain interneurons generally. (**E**) The connection probability (a-d) between different cell types is displayed both within a hemisphere and across hemispheres. Left-left and right-right as well as left-right and right-left patterns of connectivity appeared very similar. However, ipsilateral patterns of connectivity (left-left, right-right) appeared different from contralateral patterns (left-right, right-left). (**F**) Adjacency matrix displaying the difference between contralateral and ipsilateral connection probability between neuron classes. Most types of inter-class connection probability were reduced across hemispheres as compared to within a hemisphere (red). However, there were some examples of increased connection probabilities across hemispheres vs. within a hemisphere (blue). (**G**) All class connection probabilities that were stronger contralaterally are displayed. Some connection types also existed within the ipsilateral hemisphere, but were strengthened contralaterally (top panel). Other connection types were either very weak or didn’t exist ipsilaterally, but were observed in contralateral connections (bottom panel). Many of these connections involve feedback from DNs^VNC^.

**Fig. S20.**
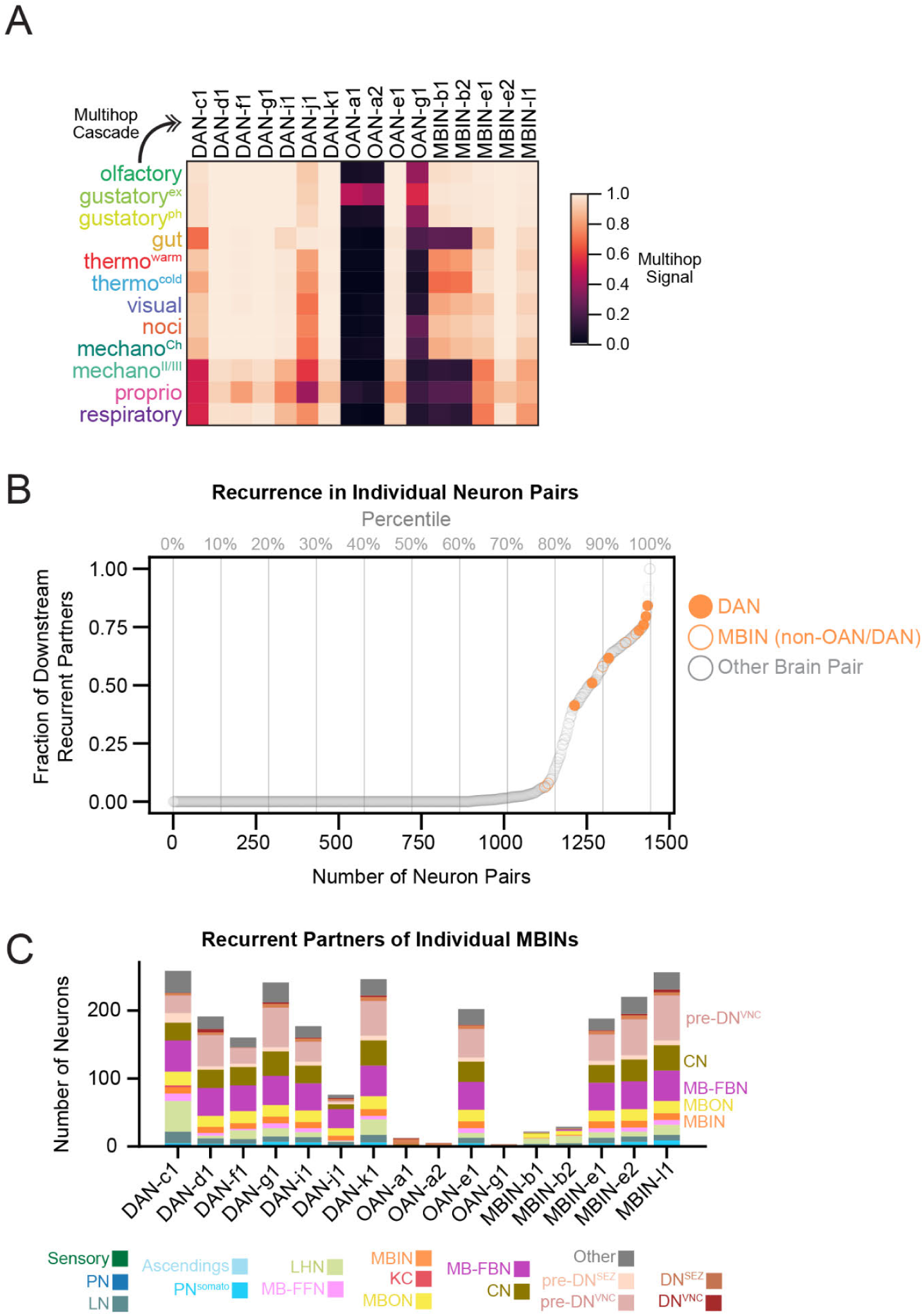
Recurrence in individual mushroom body modulatory neurons. (**A**) Signal cascades up to 5-hops maximum (a-d) to each modulatory input neuron in the learning center. Dopaminergic neurons (DANs) all integrated signal from every sensory modality. Only 33% of OANs and 60% of non-DAN MBINs integrated signal from every sensory modality. (**B**) Recurrence in individual neuron pairs, sorted from least to most recurrent. The locations of DANs and non-OAN MBINs are depicted with orange circles or hollow orange circles, respectively. Note that 4/7 DANs are more recurrent than 95% of brain neuron pairs, while the remainder are more recurrent than 90% or 80% of brain neuron pairs. Recurrence is based on a-d connectivity. (**C**) Characterization of the recurrent partners downstream of individual MBINs (a-d). Many MB-FBNs, CNs, MBONs, MBINs, and pre-DN^VNC^ neurons were observed.

**Fig. S21.**
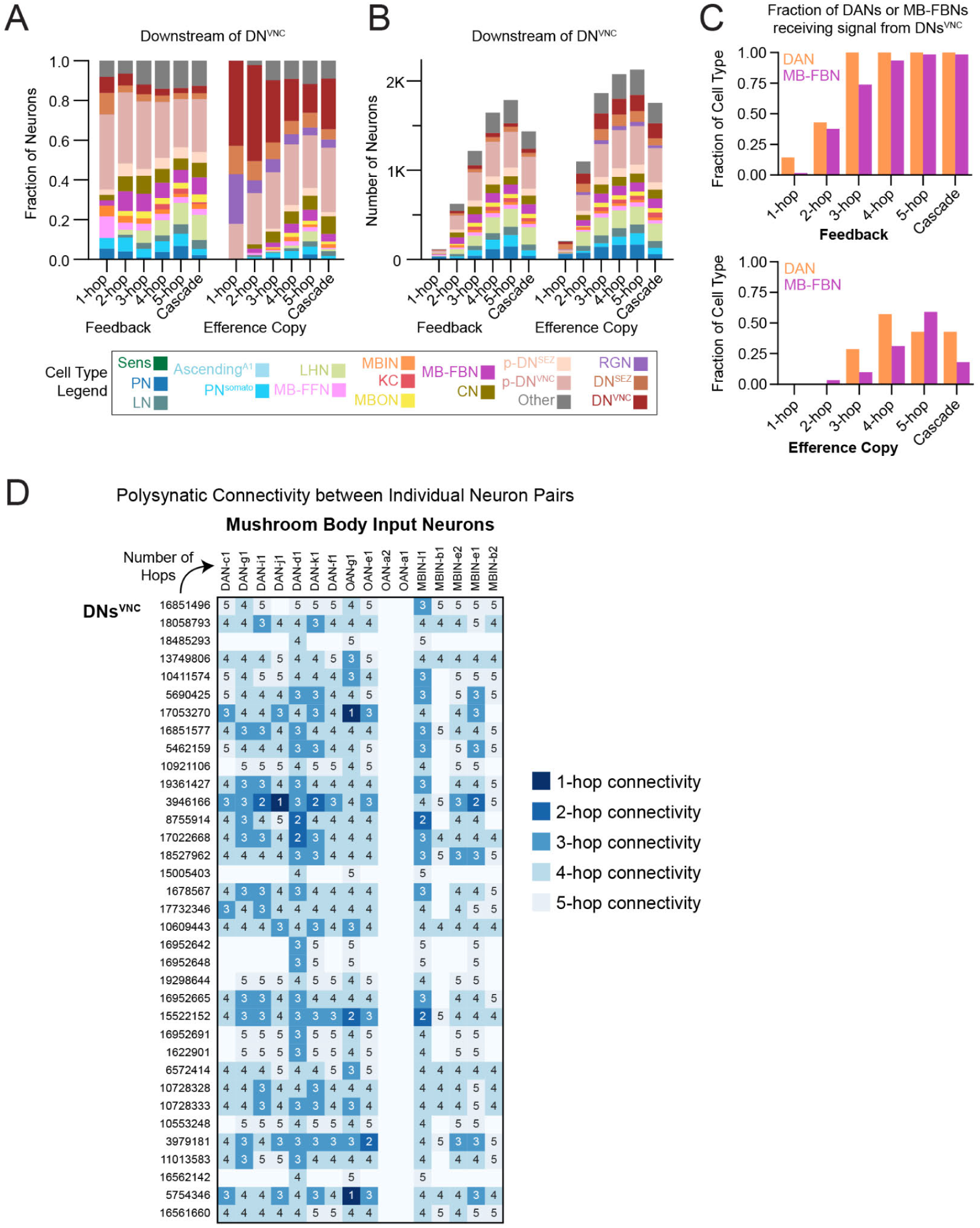
Feedback and parallel efference copy signal from descending neurons. (**A**) Cell types receiving feedback or parallel efference copy signal from DNs^VNC^, with 1-5 hops using ≥1% a-d input thresholds or 8-hops maximum using signal cascades. (**B**) Number of cell types receiving either feedback or parallel efference copy signal from DNs^VNC^. (**C**) Fraction of DANs and MB-FBNs that received polysynaptic (1-5 hops) or cascade signal from DNs^VNC^. A direct connection was observed from a DN^VNC^ to DAN-j1, as well as 2-hop feedback onto DAN-i and DAN-k1 through MB-FBNs. (**D**) Polysynaptic connectivity matrix (a-d) between individual DN^VNC^ pairs and individual MBIN pairs, indicating the number of hops between the upstream DN^VNC^ and downstream MBIN. All reported connectivity passed a ≥1% axo-dendritic input threshold. DN^VNC^ pairs are labeled by the ID of the left-side neuron.

**Fig. S22.**
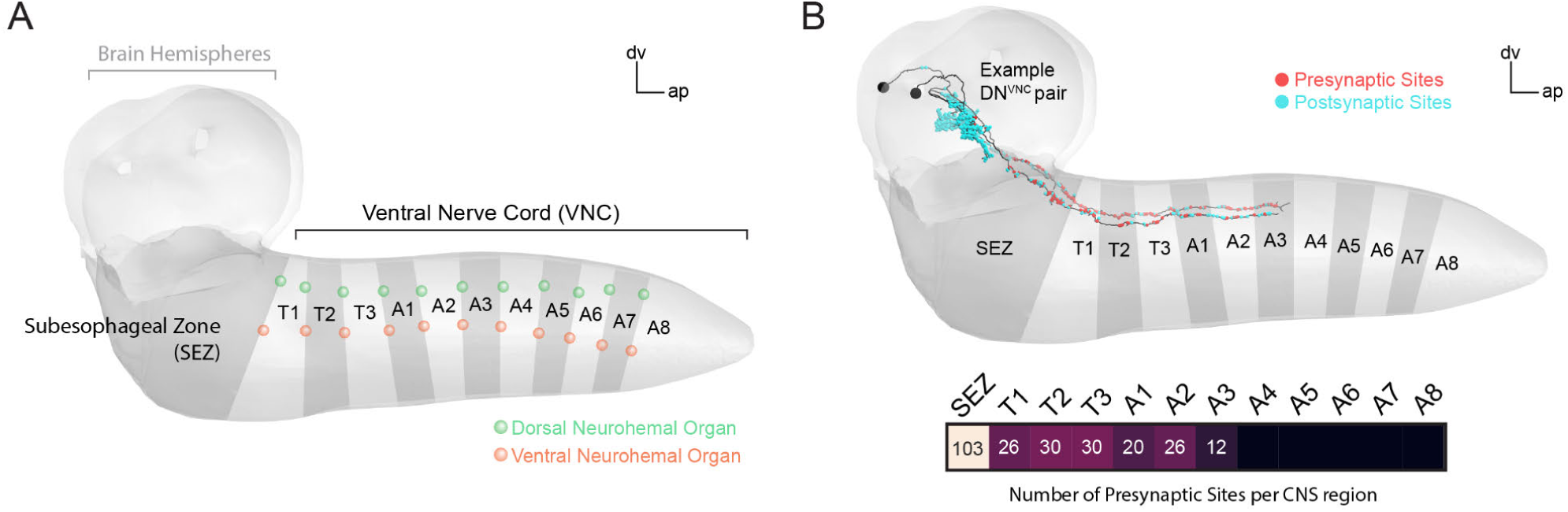
Generating a CNS projectome. (**A**) Subregions of the CNS. The boundaries between the SEZ and VNC segments were determined using stereotyped landmarks, namely dorsal and ventral neurohemal organs. The boundary between the brain and SEZ was defined using the cell bodies of ventral brain neurons. The brain was defined according to stereotyped lineage entry points (see Methods). (**B**) Example of a DN^VNC^ pair displayed within the CNS rendering. Pre- and postsynaptic sites are colored red and cyan, respectively. The number of DN^VNC^ presynaptic sites was quantified within each CNS region.

**Fig. S23.**
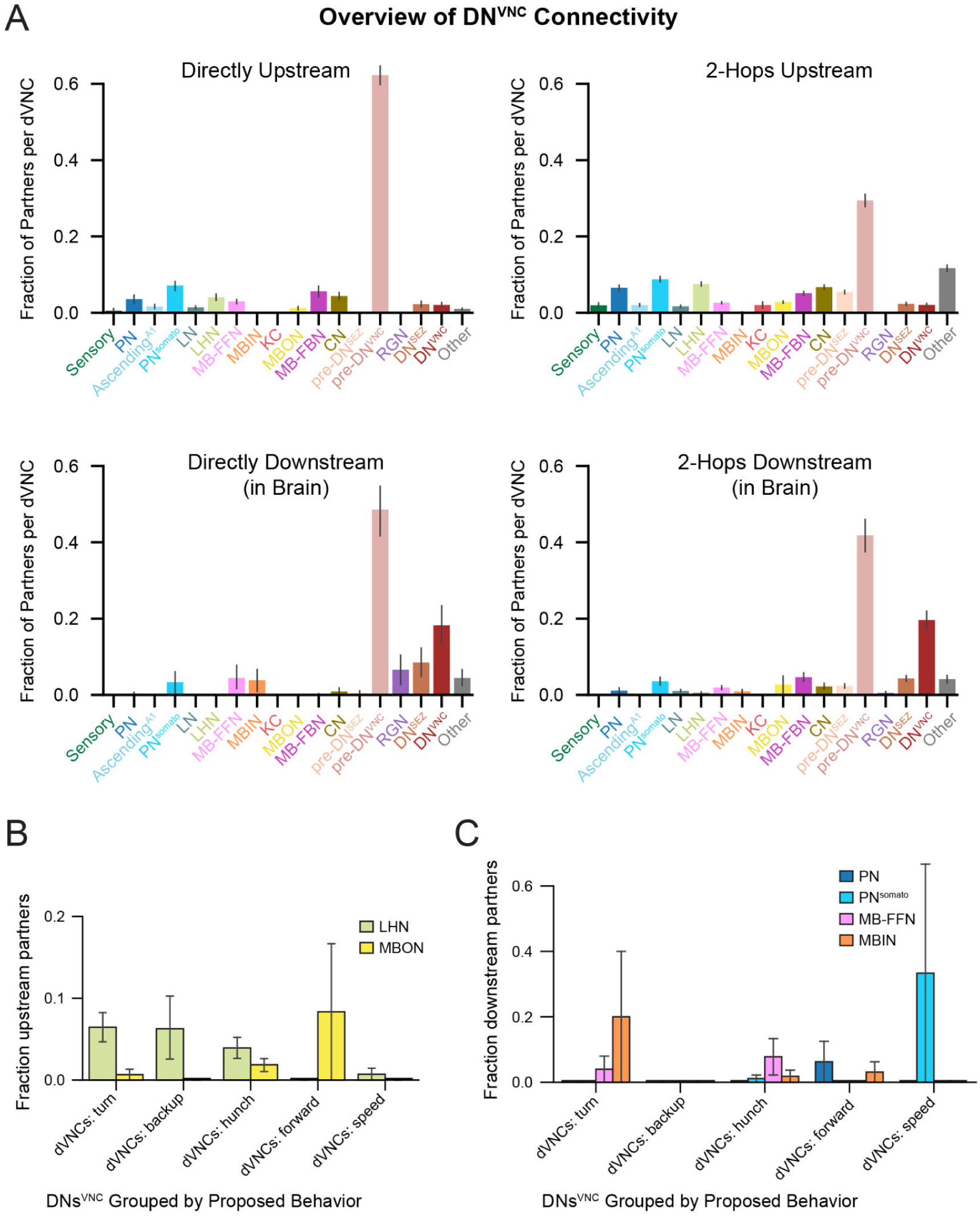
Characterization of upstream and downstream DN^VNC^ partners. (**A**) Fraction of cell types that are 1- and 2-hop upstream or downstream of DNs^VNC^ (a-d), reported as mean ± SEM. The most common partners on average were pre-DN^VNC^ neurons, both upstream and downstream. However, DNs^VNC^ also directly communicated with many different cell types. (**B**) Fraction of LHNs and MBONs that were directly upstream of DNs^VNC^ (a-d), grouped by behaviors proposed based on outputs to VNC segments (Fig. 6J). Groups of DNs^VNC^ putatively grouped by aversive behaviors displayed a higher fraction of upstream innate center neurons (lateral horn neurons, LHN), while DNs^VNC^ putatively grouped by forward crawl (appetitive behavior) displayed a higher fraction of upstream memory/learning center output neurons (MBONs). (**C**) Fraction of projection neurons (PN or PN^somato^) and MB input neuron types (MB-FFN, MBIN) directly downstream of DNs^VNC^ (a-d), grouped by proposed role in behavior.

**Fig. S24.**
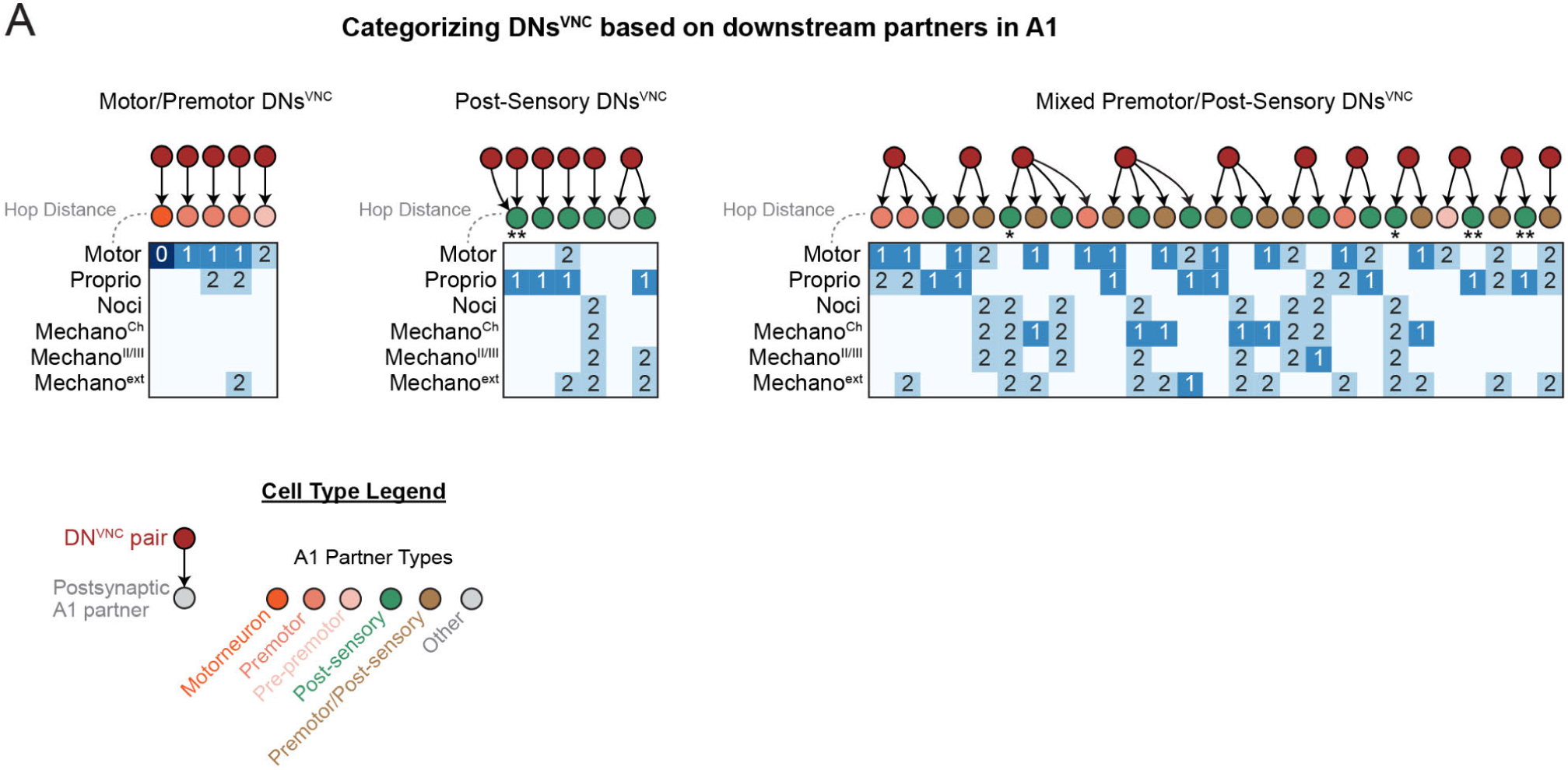
Categorization of DNs^VNC^ based on downstream partners in A1. (**A**) DNs^VNC^ were categorized using their downstream a-d partners in motor/premotor circuits, post-sensory circuits, or a mixture of premotor/post-sensory circuits. Plots depict the distance in hops between motor neurons or A1 SNs and A1 interneurons downstream of DNs^VNC^. For premotor neurons, this indicates the upstream distance (i.e. premotor neurons are 1-hop upstream of motor neurons). For post-sensory neurons, hop distance indicates the number of hops downstream of each sensory neuron modality. Postsynaptic A1 partners of DNs^VNC^ are color-coded based on their premotor or post-sensory status. * and ** indicate the two ascending neuron pairs involved in zigzag motifs (* refers to AF10-like from Fig. 7G).

**Fig. S25.**
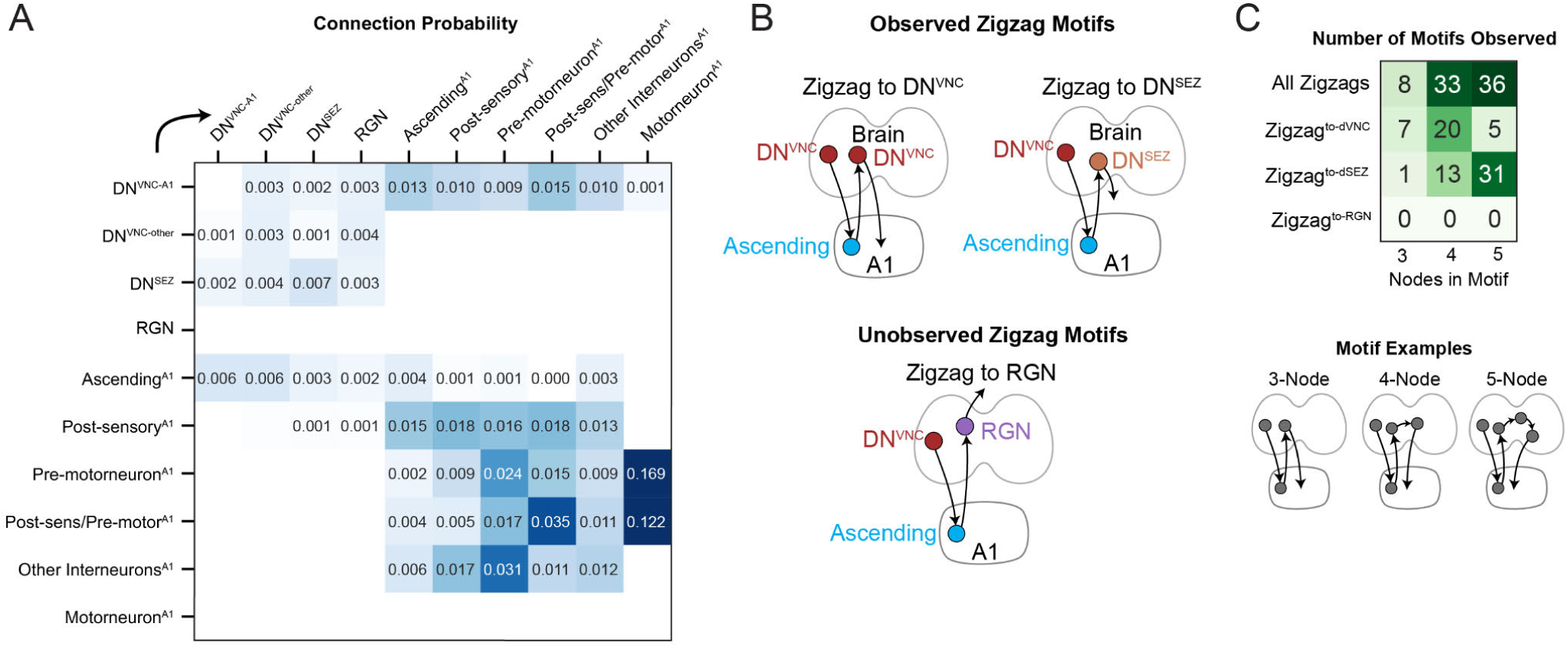
Detailed overview of brain-nerve cord interactions. (**A**) Connection probabilities (a-d) between brain output neurons and nerve cord neurons (A1 segment of the VNC). DN^VNC-A1^ neurons interact with many cell types, including ANs and post-sensory/pre-motor neurons. ANs in A1 also interact with DNs^VNC^. (**B**) Schematic representations of different possible zigzag motifs. Note that the Zigzag^to-RGN^ motif (DN^VNC^-AN-RGN) was not observed. (**C**) Observed occurrences of each motif type (a-d connectivity). The zigzag^to-RGN^ motif was never observed. Note that while diagrams in (B) displays 3-node motifs, we also quantified 4- and 5-node motifs where additional neurons were allowed between the ascending and brain output neuron (e.g. DN^VNC^-ascending^A1^-interneuron^brain^-DN^VNC^, a 4-node motif).

**Table S1.** Description of sensory and ascending neurons by modality. Brain input neurons were divided into groups based on modality. These modalities were previously described for sensory neurons, while ascending neuron modalities are based on connectivity (Fig. S2). The number of input neurons, their cell type, which tract was used to enter the brain, the associated organ for these neurons, and the neuron classes contained in each group are displayed. More detailed or additional roles are also reported. Entry Points: AN = Antennal Nerve, MxN = Maxillary Nerve, VNC = Ventral Nerve Cord. Associated Organ(s): DO = dorsal organ, TO = terminal organ, DPS = Dorsal Pharyngeal Sense Organ, DPO = Dorsal Pharyngeal Organ, PPS = Posterior Pharyngeal Sense Organ, VPS = Ventral Pharyngeal Sense Organ, ENS = Enteric Nervous System. AN-B1, AN-B2, AN-B3, MxN-B1, MxN-B2, and MxN-B3 refer to neuron classes reported in (47). Thermo-WC (warm cells) and Thermo-CC (cold cells) refer to classes reported in (48).

| Modality | Neuron Count | Neuron Type | Brain Entry Point | Associated Organ(s) | Neuron Classes | Detailed/ Additional Roles |
| --- | --- | --- | --- | --- | --- | --- |
| Olfactory | 42 | Sensory | AN | DO | Olfactory Receptor Neurons | Pheromone Sensing |
| Gustatory External | 136 | Sensory | MxN, AN | DO, TO | AN-B3, MxN-B1, MxN-B2 | Quinine, Bitter, Salt Sensing; Potential Role in Mechanosensation |
| Gustatory Pharynx | 110 | Sensory | MxN, AN | DPS, DPO, PPS, VPS | AN-B2, MxN-B3 | Sweet, Bitter, Caffeine Sensing |
| Gut Internal State | 85 | Sensory | AN | ENS | AN-B1 |  |
| Thermo-Warm | 4 | Sensory | AN | DO | Thermo-WC |  |
| Thermo-Cold | 6 | Sensory | AN | DO | Thermo-CC |  |
| Visual | 29 | Sensory | Bolwig Nerve | Bolwig Organ | Photoreceptors | Blue-tuned (Rh5) or Green-tuned (Rh6) |
| Nociceptive | 12 | Ascending | VNC-Brain Tract | VNC | Nociceptive Ascending Neurons |  |
| Mechano-Chordotonal | 10 | Ascending | VNC-Brain Tract | VNC | Chordotonal Ascending Neurons | Chordotonals have additional role in proprioception |
| Mechano-Class II/III | 2 | Ascending | VNC-Brain Tract | VNC | Class II/III Ascending Neurons | Class III sensories have additional role in cold nociception |
| Proprioceptive | 8 | Ascending | VNC-Brain Tract | VNC | Proprioceptive Ascending Neurons |  |
| Respiratory Internal State | 25 | Sensory | VNC-Brain Tract | Tracheal Network | V'td Sensory Neurons |  |

**Table S2.**
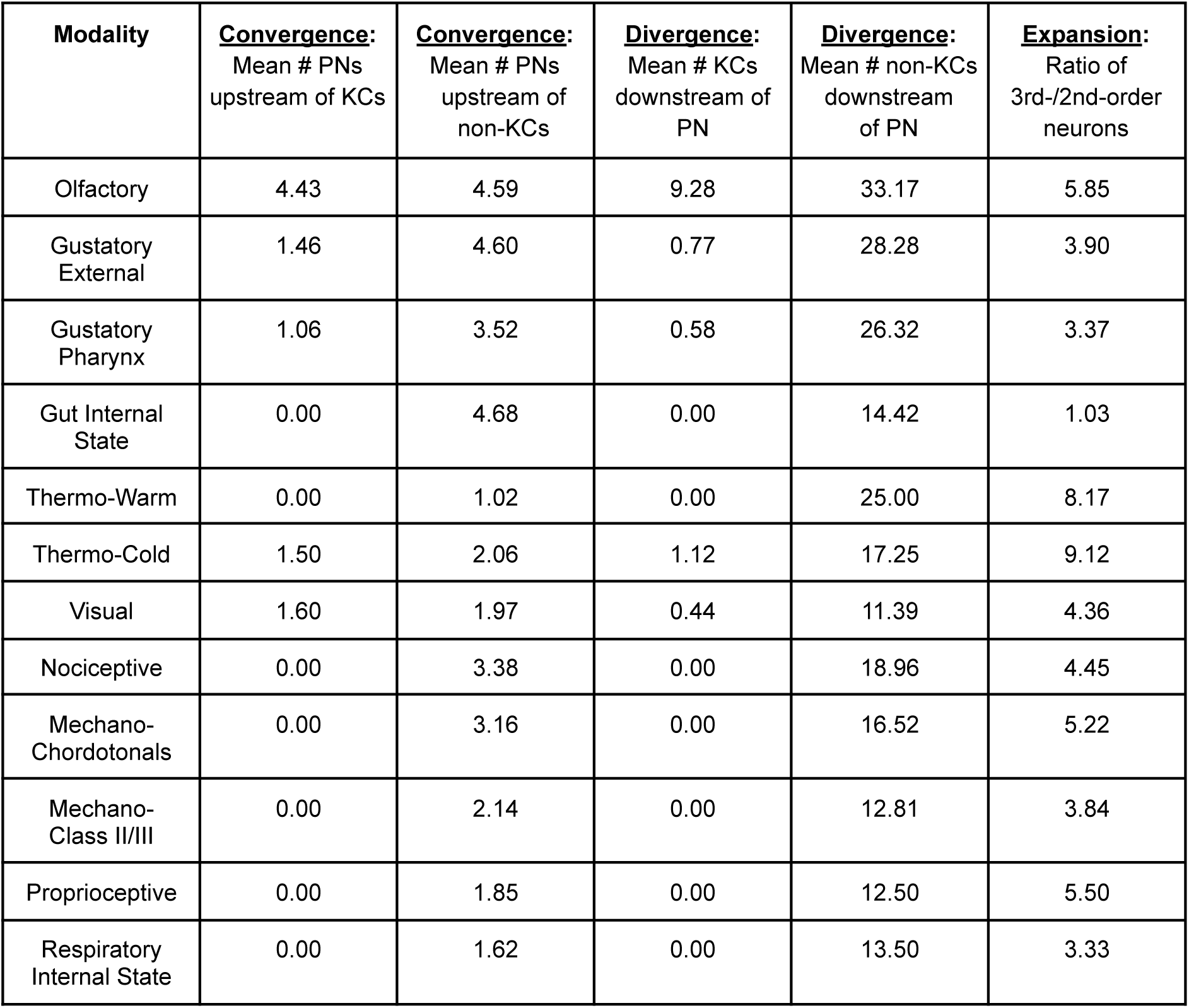
Convergence, Divergence, and Expansion between 2nd-order and 3rd-order Sensory Circuits. The mean number of PNs upstream of individual KCs or non-KCs is depicted in the first two columns (PN signal convergence). The mean number of KCs or non-KCs downstream of individual PNs is depicted in the third and fourth columns (PN signal divergence). The ratio of total number of neurons in 3rd and 2nd-order sensory circuits is depicted in the last column (signal expansion).

